# Advancing RNAi-Based Strategies Against Downy Mildews: Insights Into dsRNA Uptake and Gene Silencing

**DOI:** 10.1101/2025.03.31.646348

**Authors:** Deniz Göl, Emeka Okechukwu, Gizem Ünal, Anne Webb, Tom Woods, Yiguo Hong, Sherif M. Sherif, Theresa Wacker, David J. Studholme, John M. McDowell, Mahmut Tör

## Abstract

Downy mildew (DMs) dieases are caused by destructive obligate pathogens with limited control options, posing a significant threat to global agriculture. RNA interference (RNAi) has emerged as a promising, environmentally sustainable strategy for disease management. In this study, we evaluated the efficacy of dsRNA-mediated RNAi in suppressing key biological functions in DM pathogens of *Arabidopsis thaliana*, pea and lettuce DM pathogens, *Hyaloperonospora arabidopsidis* (*Hpa*), *Peronospora viciae* f. sp. *pisi* (*Pvp*) and *Bremia lactucae* (*Bl*), respectively. We specifically targeted the *cellulose synthase 3* (*CesA3)* and the *beta tubulin (BTUB)* genes. Silencing *CesA3* impaired spore germination and infection across multiple species, while *BTUB* silencing reinforced the potential of dsRNA-mediated inhibition. Reduction in gene expression levels correlated well with the sporulation assays confirming the effectiveness of dsRNA-mediated gene silencing. We used dsRNAs that were chemically synthesized, *in vitro* transcribed (IVT) or produced in *E. coli*. We found that the length and concentration of these dsRNAs significantly affected uptake efficiency, spore germination, and sporulation, with higher concentrations enhancing inhibitory effects. Confocal microscopy using Cy-5-labelled short-synthesized dsRNA (SS-dsRNA) provided direct evidence of spore uptake, confirming the potential of SS-dsRNA for pathogen control. However, species-specific sequence variations influenced dsRNA efficacy, underscoring the importance of target sequence design. Multiplexed RNAi impacted silencing synergisticly, further reducing germination and sporulation in *Hpa*. Additionally, we demonstrated that SS-dsRNA-mediated gene silencing is sustained over time, with a significant reduction in gene expression level at 4, 7, 10 and 11dpi. This indicates the durability and efficacy of this approach. Taken together, these findings demonstrate the potential of dsRNA-mediated gene silencing as a precision tool for managing DM pathogens.

## Introduction

Downy mildews diseases (DMs) are caused by a group of obligate biotrophic oomycetes, and are among the most destructive plant pathogens, causing severe economic losses to crops such as grapevine, lettuce, spinach, and cucurbits (Haile *et al*., 2021); (Govindarajulu *et al*., 2015; Bianca *et al*., 2023; Tör *et al*., 2023). Unlike hemibiotrophic oomycetes such as *Phytophthora* species, DM pathogens are specialized for biotrophy, depending entirely on living plant tissue for growth and reproduction (Fabro *et al*., 2011; Asai *et al*., 2014). These pathogens utilize specialized intracellular structures called haustoria to mediate nutrient uptake and secrete arrays of effector proteins to modulate host immunity. To date, the studies on these pathogens have focused primarily on characterizing their effectors, the evolutionary mechanisms underlying their success, and sustainable strategies to manage their impact on crops.

RNA interference (RNAi) has emerged as a promising tool for managing plant diseases. Host-induced gene silencing (HIGS) and spray-induced gene silencing (SIGS) are advanced RNAi-based strategies that reduce pathogen virulence by targeting genes that are critical during infection (Bilir *et al*., 2022). HIGS involves the stable expression of dsRNA constructs in transgenic plants, which silence pathogen genes through the transfer of small interfering RNAs (siRNAs) into the pathogen. This approach has demonstrated efficacy in several systems; for example, HIGS in lettuce in *Bremia lactucae* significantly reduced infection by targeting the effector *HAM34* (Govindarajulu *et al*., 2015). Similarly, HIGS targeting core pathogenicity genes (e.g., *Dicer-like* -*DCL*- or RxLR effector genes) has shown promise in controlling *Plasmopara viticola* (Haile *et al*., 2021).

SIGS, a non-transgenic RNAi-based strategy, has shown promise for controlling plant pathogens by spraying double stranded RNAs (dsRNAs) directly on to crops (Amanda *et al*., 2023). This method induces gene silencing in pathogens by targeting essential genes in pathogens, reducing disease severity. SIGS offers significant advantages as an environmentally friendly alternative to chemical pesticides, particularly against eukaryotic plant pathogens like oomycetes. Effective application of SIGS requires careful design of dsRNA constructs, stability against environmental degradation, and efficient uptake by the target pathogen.

Multiple studies have demonstrated the feasibility of using dsRNA for controlling oomycete pathogens. Ivanov and Golubeva (2023) showed that exogenous dsRNA targeting *Phytophthora infestans* elicitin genes reduced severity of lesions on potato explants. Similarly, Kalyandurg *et al*. (2021) demonstrated significant mitigation of disease symptoms by spraying potato leaves with dsRNA targeting key developmental and pathogenicity-related genes in *P. infestans* significantly mitigated disease symptoms. Sundaresha *et al*. (2022) also achieved enhanced disease resistance by targeting multiple *P. infestans* genes involved in infection and sporulation using *in vitro* synthesized dsRNAs.

Despite the successes, the broader adoption of SIGS faces challenges related to dsRNA delivery, stability, and uptake by pathogens. Studies by Wang *et al* (2023) and Zheng *et al*. (2024) addressed these issues by utilizing advanced carriers such as functionalized carbon dots and biomimetic nanovesicles, which significantly improved dsRNA stability and uptake efficiency. The addition of nanoclay carriers (Sundaresha et al., 2022) enhanced pathogen resistance and duration of stability of dsRNAs under field conditions. Mechanistic investigations revealed variability in dsRNA uptake efficiency across *P. infestans* cell types and developmental stages, highlighting the need for improved delivery strategies (Qiao *et al*., 2021).

While most research has focused on controlled laboratory conditions, environmental factors influencing dsRNA efficacy remain underexplored. Studies like Hoang *et al*. (2022) emphasized the need for understanding the barriers to dsRNA uptake in real-world settings, while Sundaresha et al. (2022) provided preliminary simulations of field applications with promising results. Taken together, these studies demonstrate the potential of SIGS as a viable strategy against oomycete pathogens. However, further research is required to optimize delivery systems, assess environmental persistence, and validate efficacy through extensive field trials to enable practical deployment of SIGS in agriculture.

In our previous studies, we targeted the *Hpa-CesA3* gene using small RNAs (sRNAs) in sense and antisense forms and found that double-stranded sRNAs were more effective at inhibiting pathogen infection than antisense sRNAs alone (Bilir *et al*., 2019). Here, we further investigated the use of short-synthesized dsRNAs (SS-dsRNAs) to target two genes across three DM pathogens, demonstrating their effectiveness for silencing essential structural genes and their broader applicability to other DM species. We also assessed the efficacy of SS-dsRNAs in silencing dual and triple gene targets, confirming that multiple genes can be effectively suppressed in parallel. Additionally, we evaluated dsRNAs of varying lengths and identified limitations in dsRNA uptake by DM spores. Using confocal microscopy, we visualized fluorescent SS-dsRNA internalization in DM spores and germ tubes and compared the gene-silencing effectiveness of chemically synthesized, *in vitro*-transcribed, and *E. coli*-expressed dsRNAs.

## Results

### Silencing *CesA3* gene across different DM pathogens inhibits germination and infection

Previously, we focused on the *Arabidopsis thaliana-Hpa* interactions using *Hpa-CesA3* in our sRNA experiments (Bilir *et al*., 2019). We took this further by investigating whether the same approach could be used in other obligate DM pathogens, such as *Peronospora viciae* f. sp. *pisi* (*Pvp)* and *Bremia lactucae* (*Bl)*, pathogens of pea and lettuce DM, respectively. Using the *Hpa-CesA3* sequence as the query, we conducted searches against genomic sequences of recently assembled *Pvp* (Webb et al, unpublished) and publically available *Bl* genome sequences using BLASTN and BLASTX (Altschul *et al*., 1997). High-confidence orthologs were identified based on significant sequence similarity and conserved domain architecture. The identified orthologs were designated as *Pvp-CesA3* for pea-infecting DM and *Bl-CesA3* for lettuce-infecting DM. *Hpa-CesA3* was 83% to *Pvp-CesA3* and 81% identical to *Bl-CesA3*, while *Pvp-CesA3* was 79% identical to *Bl-CesA3* (Supplemental Figure 1).

Previously, we targeted the *Hpa-CesA3* gene using sense and antisense sRNAs individually and found that sRNA duplexes (dsRNAs) were more effective than antisense sRNAs alone (Bilir *et al*., 2019). However, in this current study, only 30 bp SS-dsRNAs were used. We first checked the original *Hpa-CesA3*-specific 30 bp SS-dsRNA against the *Bl-CesA3* and *Pvp-CesA3* genes, and the sequence alignment in this area showed 73.3% to 76.6% identity, respectively (Figure 1A). Secondly, we also designed a new SS-dsRNA, designated *CesA3*-common, from a highly conserved region of the alignment (Supplemental Figure1) where the sequence identities were 100% for *Pvp* and 83.3% for *BI* (Figure 1B). Next, we checked for off-target matches by subjecting the SS-dsRNA sequences to BLAST searches against the genomic sequences of respective pathogens (*Hpa, Pvp,* and *Bl*) and the host plant (*Arabidopsis*, pea, and lettuce) genomes. The sequences did not show any detectable similarity to non-target genes and host-plant genomes.

**Figure 1.**
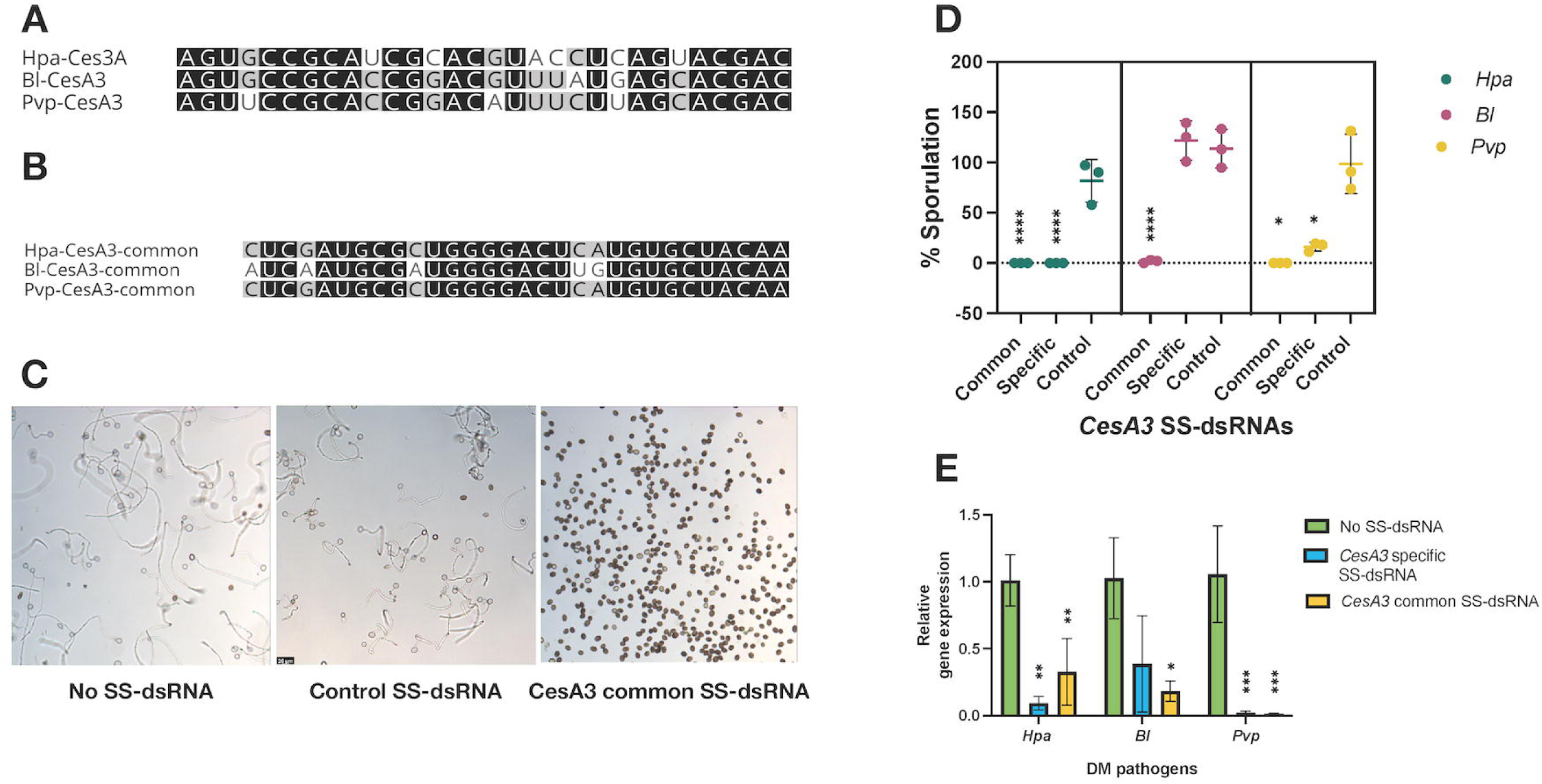
Targeting *CesA3* genes in *Hpa, Bl* and *Pvp* with SS-dsRNAs inhibit spore germination and infection. **A)** *Hpa-CesA3* specific SS-dsRNA targeting *Hpa-CesA3* was designed based on sequences from (Bilir et al., 2019), and alignment with *Bl-CesA3* and *Pvp-CesA3* sequences was performed, **B)** *Hpa-CesA3* common SS-dsRNA targeting a conserved region of *CesA3* across *Hpa*, *Bl*, and *Pvp* was designed using alignment data, **C)** Spore germination assays were conducted using the designed SS-dsRNAs. Representative images of *Pvp* spore germination are shown, **D)** Sporulation assays were performed with the SS-dsRNAs, assessing at 7 dpi for *Hpa* and *BI*, and at 8 dpi for *Pvp*, expressed as a percentage of the control, **E)** Expression levels of *CesA3* genes in all three pathogens were measured by qRT-PCR at 4 dpi. Statistical significance was analyzed using a one-way ANOVA, comparing the means of SS-dsRNA-treated and control samples. Values marked with an asterisk (*) indicate significant differences from the control group. Bars represent the standard deviation of three independent replicates. Adjusted p-values for sporulation are as follows: *Hpa*: **** (p < 0.0001); *BI*: *CesA3* common **** (p < 0.0001),; *Pvp*: *CesA3*-common * (p=0.0169), *Hpa*-*CesA3*-specific * (p=0.0432). Adjusted p-values for qRT-PCR are as follows: *Hpa*: *Hpa*-*CesA3-* specific **(p=0.0016), *CesA3*-common **(p=0.0071). *BI*: *CesA3*-common *(p=0.0167). *Pvp*: *Hpa*-*CesA3*-specific **(p=0.0015), *CesA3*-common **(p=0.0014).

To evaluate the effectiveness of the designed duplexes, we conducted*, in vitro* germination and *in planta* sporulation assays were conducted with SS-dsRNAs targeting the *CesA3* gene in three DM pathogens to evaluate the effectiveness of the designed duplexes. For analysis of germination and sporulation inhibition, percentages were calculated relative to the control group. Spores from *Hpa*, *Pvp*, and *Bl* in the untreated control (no SS-dsRNA) or those treated with a control SS-dsRNA germinated well. In contrast, treatment with the *Hpa-CesA3*-specific or the *CesA3*-common SS-dsRNAs significantly inhibited spore germination across all pathogens (Figure 1C for a representative image). For the *in planta* assays, sporulation varied by pathogen. In *Hpa*, both the *CesA3*-common (\*\*\*\**, p < 0.0001) and Hpa-CesA3-*specific *(\**, p = 0.0432) SS-dsRNAs significantly inhibited sporulation, while the negative control SS-dsRNA allowed full sporulation (Figure 1D). Similarly, in *BI*, the *CesA3*-common SS-dsRNA significantly reduced sporulation (****, p < 0.0001), whereas the *Hpa-CesA3*-specific and control SS-dsRNAs allowed full sporulation, highlighting the effects of single-nucleotide mismatches between the SS-dsRNAs and the pathogen target. In *Pvp*, the *CesA3*-common SS-dsRNA completely inhibited sporulation (*, p = 0.0169), while the *Hpa-CesA3*-specific SS-dsRNA significantly reduced sporulation (*, p=0.0432). The negative control SS-dsRNA permitted full sporulation (Figure 1D).

We then assessed the expression levels of *CesA3* genes in infected tissues using qRT-PCR for each DM pathogen separately, following treatment with the *Hpa-CesA3*-specific and the *CesA3*-common SS-dsRNAs in all three DM pathogens. There were statistically significant reductions in the expression levels of *CesA3* genes in all tested pathogens (Figure 1E). In *Hpa*, both *Hpa-CesA3*-specific (**, p = 0.0016) and *CesA3*-common (**p = 0.0071) SS-dsRNAs significantly reduced gene expression. Similarly, in *BI*, the *CesA3*-common SS-dsRNA (*, *p = 0.0167*) led to a significant reduction. In *Pvp*, both *Hpa-CesA3*-specific (**, p = 0.0015) and *CesA3*-common (**, p = 0.0014) SS-dsRNAs also significantly reduced expression. These results align with the sporulation assays, confirming that SS-dsRNA application effectively silences the targeted genes.These results suggest that *CesA3* genes from different DM pathogens can be targeted using a single SS-dsRNA. Additionally, SS-dsRNAs can be designed either for specific pathogens or for a broader class of pathogens by targeting conserved gene regions.

### Targeting *BTUB* gene in *Hpa, Pvp*, and *Bl* further confirms gene silencing in downy mildews

To confirm our results with *CesA3* and demonstrate that conserved genes can be used as targets in different DM pathogens, we applied the same approach to the *BTUB* gene in *Hpa*, *Pvp*, and *Bl*. We used reciprocal BLASTN and BLASTX (Altschul et al., 1997) searches of *Hpa-BTUB* against genomic sequences and identified its orthologs in *Hpa, Pvp,* and *Bl*; and the identified orthologs were designated as *Pvp-BTUB* and *Bl-BTUB*. Multiple sequence alignments (Supplemental Figure 2) indicated that *Hpa-BTUB* was 87% and 84% identical to *Pvp-BTUB* and *Bl-BTUB*, respectively, and that the amino acid sequences of these three genes contained the beta-tubulin domain IPR002453. Then, we designed a 30-nt SS-dsRNAs from a highly conserved region of the alignment (Supplemental Figure 2) where the sequence identities were 83.3% for BI and 90% for *Pvp* (Figure 2A) and the BLAST searches against the pathogens’ respective host genomes did not identify any detectable sequence similarity to non-target genes.

**Figure 2.**
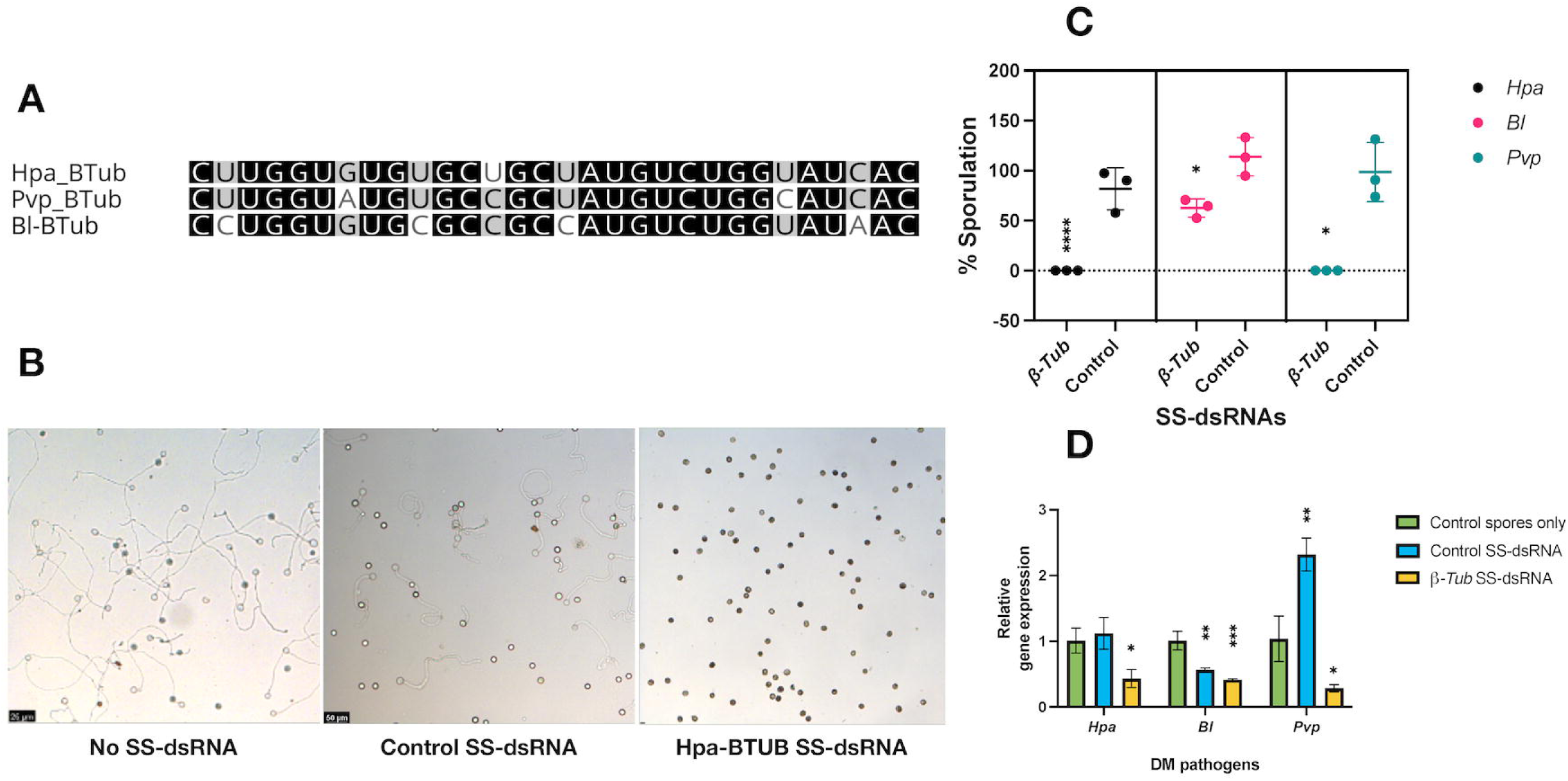
Silencing *BTUB* genes in *Hpa, Bl* and *Pvp* inhibits spore germinations and alter sporulation. **A)** A 30 bp SS-dsRNA was designed form a conserved region identified through sequence alignment, **B)** Spore germination assays were carried out using the designed SS-dsRNAs. Representative images of *Hpa* spore germination are shown, **C)** Sporulation assays were performed with the SS-dsRNAs, assessing *Hpa*, *Bl*, and *Pvp* sporulation at 7 dpi, expressed as a percentage of the control. **D)** Expression levels of *BTUB* genes in all three pathogens were measured by qRT-PCR at 4 dpi. Statistical significance was assessed using a one-way ANOVA for sporulation and t-test for qRT-PCR, comparing the means of SS-dsRNA-treated and control samples. Values marked with an asterisk (*) indicate significant differences from the control group. Error bars represent the standard deviation of three independent replicates. Adjusted p-values for sporulation: *Hpa*: **** (p < 0.0001); *BI*: * (p=0.0273); *Pvp*: * (p=0.0169). Adjusted p-values for qRT-PCR are as follows: *Hpa*: *(p=0.0130). *BI*: **(p=0.0019). *Pvp*: *(p=0.0207).

*In vitro* spore germination assays showed that a single SS-dsRNA that targets *BTUB* genes inhibited spore germination in all three DM pathogen spores (Figure 2B). In planta sporulation assays with SS-dsRNA resulted in total inhibition of sporulation in both *Hpa*-(****, p < 0.0001) and *Pvp*-infected (*, *p = 0.0169*) *Arabidopsis* and pea plants, respectively. Additionally, after treatment with SS-dsRNA, *Bl* spores showed significantly reduced sporulation on lettuce plants (*, *p = 0.0273*) (Figure 2C), indicating the effect of sequence variation within the SS-dsRNA on silencing.

Gene expression of *BTUB* in all three DM pathogens was quantified using qRT-PCR after *Hpa-BTUB* SS-dsRNA treatment. A significant reduction in *BTUB* gene expression levels was observed across all tested pathogens (Figure 2D), with *Hpa* (*, *p = 0.0130*), *BI* (**, p = 0.0019), and *Pvp* (*, *p = 0.0207*). These results confirm that specific genes can be targeted across multiple DM-causing species, and the observations with *BTUB* SS-dsRNA align with those observed for *CesA3* SS-dsRNA, demonstrating broader applicability rather than being limited to a single target.

### The lengths of SS-dsRNAs influences their uptake by spores

Previously, we used 24-, 25-, and 30-nt single-stranded antisense sRNAs to investigate their effects on pathogen sporulation (Bilir et al., 2019). To advance this work and explore size constraints on dsRNA uptake by spores, we expanded our analysis to include a broader range of dsRNA lengths. Chemically synthesized *CesA3* SS-dsRNAs (21- to 75 bp) were tested for their impact on spore germination rates in *Hpa* and *Pvp* (Table 1). Significant differences were observed in the effects of SS-dsRNA lengths on spore germination for both pathogens.

**Table 1.** Effect of *CesA3* common SS-dsRNA length on spore germination rates of *Hpa* and *Pvp*.

| Target<br>Gene | SS-dsRNA length (bp)<sup>1</sup> | % *Hpa* spore<br>germination<sup>2</sup> | % *Pvp* spore<br>germination<sup>2</sup> |
| --- | --- | --- | --- |
| *CesA3* | 21 | 154.72 (± 13.66) | 73.90 (±1.20) |
|  | 22 | 114.33 (± 12.17) | 73.05 (±1.25) |
|  | 23 | 115.66 (± 11.45) | 77.46 (±1.19) |
|  | 24 | 153.11 (± 65.69) | 75.64 (±0.99) |
|  | 25 | 247.10 (± 45.88) | 76.69 (±1.00) |
|  | 30 | 0 | 0 |
|  | 50 | 0 | 0 |
|  | 75 | 0 | 0 |

1- bp: base pair
2-Spore germination percentages are presented as means of three replicas with standard deviations (±).

For *Hpa*, shorter dsRNAs (21–25 bp) exhibited variable spore germination rates, with the highest rate at 25 bp (247.10%) and the lowest at 22 bp (114.33%). In contrast, longer dsRNAs (30, 50, and 75 bp) completely inhibited spore germination. Similarly, for *Pvp*, dsRNAs in the 21–25 bp range resulted in germination rates ranging from 73.05 to 77.46%, with minimal variation among these lengths. As with *Hpa*, dsRNAs of 30, 50, and 75 bp entirely inhibited spore germination.

Since it was not feasible to synthesize dsRNA longer than 75 bp, we utilized *in vitro*-transcribed (IVT) or *E. coli*-produced dsRNAs of various lengths. DNA fragments of the *Hpa-CesA3* (285 bp), *Hpa-BTUB* (273 bp), and *E. coli GUS* (255 bp, as a control) were cloned and expressed in *E. coli* (Supplemental Figure 3). Germination assays with *Hpa* spores showed that the germination was not totally inhibited by E. coli produced dsRNAs (Supplemental Figure 4), but germination rates with dsRNAs were reduced for the *Hpa-CesA3*, *Hpa-BTUB* and *GUS* with 79.52%, 66.61%, and 48.65%, respectively. However, when these dsRNA fragments were digested with RNase III and retested, all RNase III-digested dsRNAs completely inhibited *Hpa* spore germination, confirming that dsRNA uptake by spores is size-dependent. Interestingly, GUS dsRNA, derived from the *E. coli GUS* gene and lacking sequence similarity to the *Hpa* genome, revealed that not all dsRNAs are suitable as negative controls.

Next, we tested 50- and 75-bp *CesA3* SS-dsRNAs in sporulation assays with both DM pathogens *Hpa* and *Pvp*. As expected, in pea plants, uninfected leaves were green (Figure 3A), and control leaves infected with spores only showed some chlorosis (Figure 3B) as expected. However, 30, 50 and 75 bp SS-dsRNA completely inhibited pathogen sporulation on pea plants (Figure 3 C-E). However, 30 bp SS-dsRNA did not induce any symptoms. In contrast, 50 and 75 bp dsRNAs caused chlorosis and curliness on pea leaves (Figure 3C-E), but excessive necrosis was not observed, likely due to the resilience of the leaves. Similar results were obtained with the *Arabidopsis-Hpa* pathosystem; uninfected (Figure 4A) and control dsRNA treated seedlings (Figure 4B) sporulated, 30 bp SS-dsRNA did not induce symptoms in plants (Figure 4C), but both 50 and 75 bp SS-dsRNAs caused chlorosis on *Arabidopsis* cotyledons and caused seedling necrosis at 4 dpi (Figure 4D).

**Figure 3.**
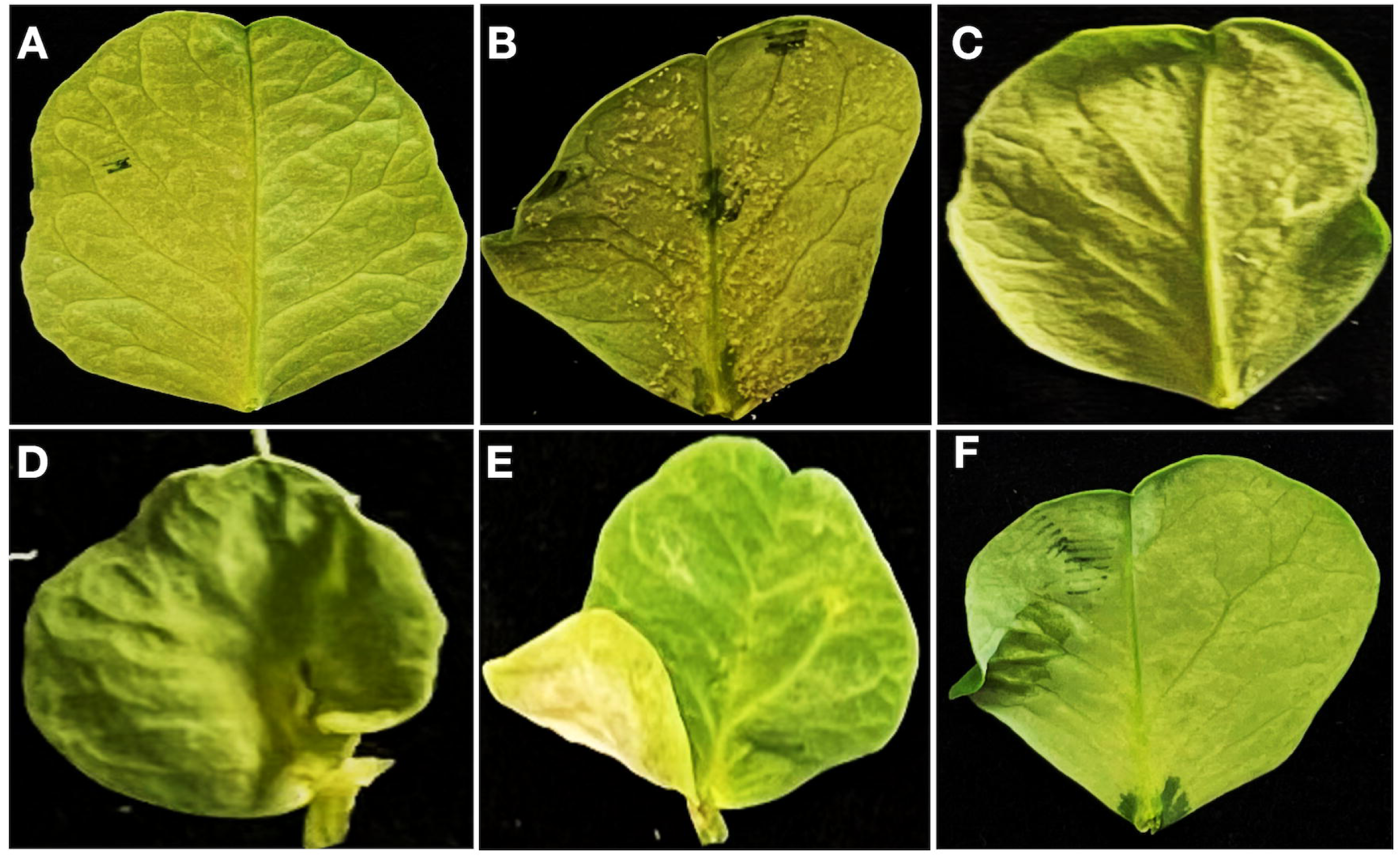
Effects of dsRNA length on sporulation in pea leaves. Pea leaves were drop-inoculated with *Pvp* spores mixed with *CesA3* dsRNAs of varying lengths, and sporulation was assessed at 7 dpi. **A)** Uninoculated control leaf, **B)** Control leaf inoculated, without dsRNA, **C)** Leaf inoculated with 30 bp SS-*CesA3* common dsRNA, **D)** Leaf inoculated with 50 nt SS-*CesA3* dsRNA, **E)** Leaf inoculated with 75 bp SS-*CesA3* dsRNA, **E)** Leaf inoculated with 100 bp IVT-produced *CesA3* dsRNA.

**Figure 4.**
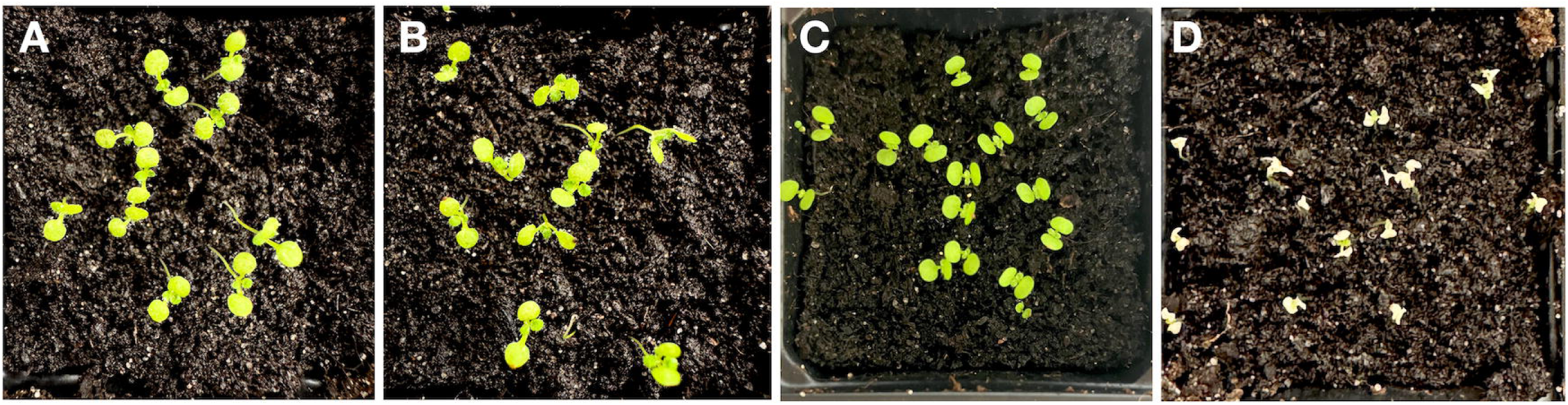
Impact of *Hpa-CesA3* dsRNA length on *Arabidopsis* responses. Seven-day-old *Arabidopsis* cotyledons were drop-inoculated with *Hpa* spores mixed with either 30-nt or 75-nt *CesA3* dsRNAs and evaluated at 4 dpi. **A)** Control seedlings treated with dsRNA only, **B)** Seedlings treated with 30-nt control scrambled dsRNA, **C)** Seedlings treated with 30 bp *Hpa-CesA3* dsRNA, **D)** Seedlings treated with 75 bp *Hpa-CesA3* dsRNA.

Building on this, we used 100 bp IVT-produced *CesA3* dsRNA and tested its effect on *Pvp* spore germination and infection. The 100 bp dsRNA completely inhibited spore germination. In sporulation assays with *Pvp*, the 100 bp dsRNA produced results similar to the 50 and 75 bp dsRNAs, with no sporulation observed. However, chlorosis was again noted on pea leaves (Figure 3E), further highlighting possible off-target effects in the host rather than in the pathogen.

### The concentration of SS-dsRNAs affects both the germination rate and the sporulation rate of *Hpa*

We have established a high-throughput screening protocol in *Hpa* to decipher gene functions using SS-dsRNA. Our previous study (Bilir *et al*., 2019) demonstrated the dose-dependent effects of dsRNAs on sporulation inhibition. Building on this, we targeted 11 genes (Table 2 and Supplemental Table 3) and optimised the concentration to use in our programme. Since dsRNAs were found to be more effective than antisense sRNAs, we tested lower concentrations (1 and 5 µM) for SS-dsRNAs, compared to the higher concentrations (5, 10, and 20 µM) used for antisense sRNAs.

**Table 2.** Germination and sporulation rates of *Hpa* following gene targeting using two SS-dsRNA concentrations.

| Target<br>Transcript<br>IDs | % Germination<sup>2</sup> |  |  | % Sporulation<sup>2</sup> |  |  |
| --- | --- | --- | --- | --- | --- | --- |
|  | SS-dsRNA<sup>1</sup> Concentrations |  |  |  |  |  |
|  | 1μM | 5μM | Statistical<br>Significance<sup>3</sup> | 1μM | 5μM | Statistical<br>Significance<sup>3</sup> |
| HpaT814170 | 0 | 0 | N/A<sup>4</sup> | 152.24 | 0 | **** |
| HpaT812660 | 33.82<sup>2</sup> | 0 | * | 87.76 | 0 | **** |
| HpaT811651 | 117.74 | 0 | * | 139.30 | 0 | ** |
| HpaT800673 | 183.44 | 0 | ** | 75.90 | 33.76 | * |
| HpaT801549 | 145.32 | 0 | * | 56.61 | 35.71 | **** |
| HpaT809280 | 70.74 | 0 | ** | 106.97 | 50.79 | ** |
| HpaT813100 | 129.52 | 0 | ** | 65.75 | 70.65 | n.s |
| HpaT810056 | 283.43 | 244.39 | n.s | 199.50 | 22.02 | *** |
| HpaT802074 | 249.61 | 37.09 | ** | 79.25 | 14.35 | **** |
| HpaT802307 | 117.19 | 184.11 | * | 33.43 | 50.87 | * |
| HpaT813395 | 137.74 | 197.77 | n.s | 124.38 | 42.90 | n.s |

**1.SS-dsRNA:** Short synthesized 30-bp double-stranded RNA.
**2.Normalization:** Germination and sporulation rates are expressed as percentages normalized to untreated *Hpa* controls (set to 100%).
**3.Statistical significance (1μM vs. 5μM):** Results of unpaired t-test; significance levels: ns = not significant, \*p < 0.05, \*\*p < 0.01, \*\*\*p < 0.001, \*\*\*\*p < 0.0001.
**4.HpaT814170 germination:** Both 1μM and 5μM resulted in 0% germination (complete inhibition). Statistical comparison not applicable (N/A).

When targeting gene *HpaT814170*, no germination was observed at either concentration. However, at 1 µM, sporulation increased statistically significantly to 152.24%, whereas no sporulation occurred at 5 µM. In the case of *HpaT812660*, germination was 33.82% at 1 µM, but both germination and sporulation were completely inhibited at 5 µM. Similarly, *HpaT811651* exhibited germination and sporulation rates of 117.74% and 139.30%, respectively, at 1 µM. However, at 5 µM, both processes were entirely suppressed. These findings suggest a concentration-dependent regulatory effect of SS-dsRNA on *Hpa* development, warranting further investigation into the underlying mechanisms. Conversely, some SS-dsRNAs, such as those targeting *HpaT810056*, did not inhibit germination and sporulation; in fact, germination was enhanced at 1 µM (283.43%), though sporulation was reduced statistically significantly at 5 µM (22.02%).

Similarly, silencing *HpaT800673* and *HpaT801549* reduced sporulation to below 36% at 5 µM, but germination remained high at 1 µM, reaching 183.44% and 145.32%, respectively. SS-dsRNAs targeting *HpaT802074* and *HpaT802307* exhibited moderate efficacy, with sporulation being more effectively inhibited at 5 µM, while germination showed either a slight reduction or enhancement.

For *HpaT813395*, the results were more variable, with germination rates increasing to 137.74% at 1 µM and 197.77% at 5 µM compared to the control, whereas sporulation decreased from 124.38% at 1 µM to 42.90% at 5 µM relative to the control (Table 2). Overall, these findings indicate that SS-dsRNA efficacy depends not only on concentration but also on the target gene, as determined by the combined effects on spore germination and inhibition of sporulation. Some SS-dsRNAs enhanced germination or sporulation at lower concentrations while effectively inhibiting them at higher doses. Targeting certain genes, such as *HpaT814170* and *HpaT812660*, resulted in consistent inhibition at both concentrations, whereas targeting others, such as *HpaT810056*, led to substantial variations between the two concentrations tested. In contrast, targeting *HpaT813395* displayed no significant changes in germination or sporulation across different SS-dsRNA concentrations (Table 2).

These results highlight the complexity of gene silencing in *Hpa* and suggest that some genes are more amenable to effective targeting with SS-dsRNAs than others. Additionally, these data also indicate a) the need for optimized concentration and gene selection to achieve consistent pathogen control, and b) germination assays carried out *in vitro* may not always correlate with that of *in planta* sporulation assays.

### Multiplexed SS-dsRNA silencing inhibits *Hpa* germination and sporulation

We wanted to investigate the efficacy of multiplex silencing using multiplexed SS-dsRNAs. We selected three *Hpa* genes (*HpaT802064*, *HpaT802452*, and HpaT803108, see Supplemental Table 3 for their possible functions; also referred to as G1, G2 and G3, respectively in Figure 5), whose targeting did not cause full inhibition of spore germination and pathogen sporulation. The selection of target genes for the multiplexing study was based on the screening data in our research, similar to the approach used in the concentration effect study. In the spore germination assay, although individual SS-dsRNAs did not cause any changes in spore germination rate (Figure 5A), targeting two or three genes simultaneously, fully inhibited spore germination, demonstrating the additive effect of multiplexed silencing.

**Figure 5.**
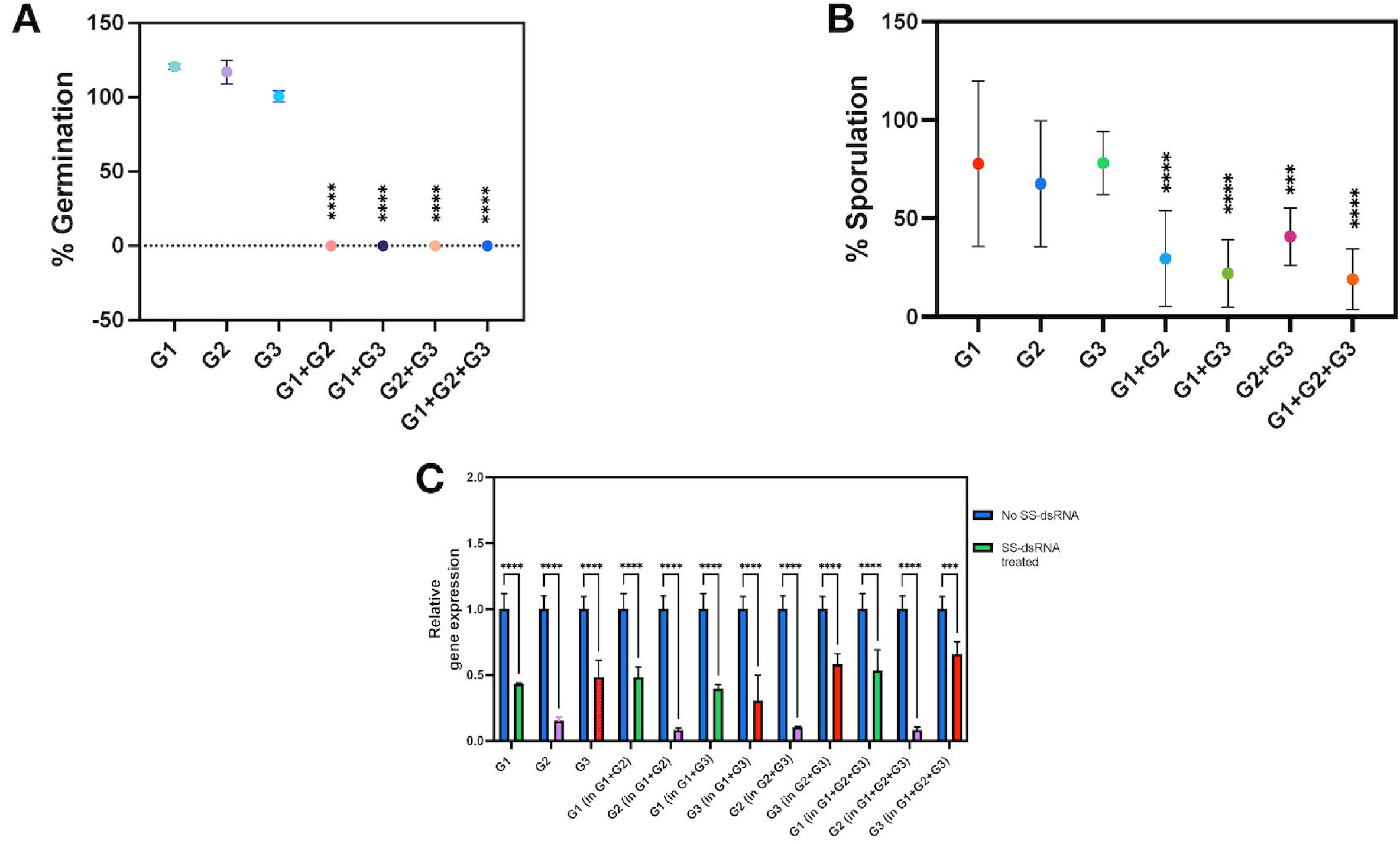
Effects of multiplexed SS-dsRNAs on the germination and sporulation of *Hpa*. Three SS-dsRNAs targeting different genes (802064, 802452, and 803108; G1, G2 and G3, respectively) were tested individually, as well as in pairs and triplets. **A)** Germination rates assessed at 1dpi, **B)** Sporulation rate at 7dpi, **C)** Analysis of gene expression for each gene after SS-dsRNA treatments using qRT-PCR. Gene expression levels were normalised to the control gene (*Actin*). Data are expressed using 2^-ΔΔCt^ method to calculate the relative fold gene expression level of samples. Statistical significance was assessed using a one-way ANOVA, comparing the means of SS-dsRNA-treated and control samples. Values marked with an asterisk (*) indicate significant differences from the control group. Bars represent the standard deviation of three independent samples. **** (p < 0.0001) and *** (p=0.0009).

Similarly, in the sporulation assay, while individual SS-dsRNAs caused only non-significant reductions in sporulation rate, targeting two or three genes simultaneously enhanced suppression of sporulation (Figure 5B), confirming further the additive effect of multiplexing.

The gene expression data provided valuable insights into SS-dsRNA-based silencing, particularly in the multiplex experimental design. qRT-PCR analysis confirmed the efficacy of SS-dsRNA treatments (Figure 5C), showing a significant reduction in expression levels for all three target genes. The strongest knockdown was observed in multiplex treatments targeting all three genes (****, *p* < 0.0001).

Among the targets, gene G2 (*HpaT802452*) exhibited the most pronounced inhibitory effect when silenced individually, consistently showing greater expression reduction than other genes in both pairwise and triple multiplex treatments. Additionally, G1 and G2 displayed consistent expression changes across all conditions, whether targeted individually or in combination. This consistency suggests that SS-dsRNA-based silencing is highly target-specific, with the inclusion of multiple nucleotide sequences not affecting specificity.

Taken together, these results highlight the potential of multiplexed SS-dsRNA treatments to enhance silencing efficiency, effectively disrupting germination, sporulation, and gene expression in *Hpa*.

### Confocal microscopy detects uptake of fluorescently labelled SS-dsRNA

We observed significant alterations in germination and sporulation rates upon targeting numerous genes with SS-dsRNAs. To further validate these findings and provide mechanistic insight, we investigated whether spores of DM-causing pathogen actively internalize dsRNAs, thereby ruling out potential indirect effects. To investigate this, we employed confocal microscopy to visualize the uptake and cellular localization of Cy5-labeled SS-dsRNAs in *Hpa* and *Pvp* spores. This approach allowed us to directly assess dsRNA uptake efficiency and variability in absorption among spores, strengthening the link between dsRNA exposure and the observed phenotypic effects. Pathogen spores were incubated with Cy-5-labelled dsRNA for 6 hours and imaged using a Nikon Ti2-E AX NSPARC laser point-scanning confocal microscope. Fluorescent signals corresponding to Cy-5-labelled dsRNA were detected in the spores and germ tubes, confirming internalization.

Single-plane Z-stack images (Figure 6A) and 3D-rendered projections (Figure 6B) of *Hpa* spores revealed discrete RNA aggregates within the spore cytoplasm. Similarly, in *Pvp* spores, Cy-5-labelled dsRNA was observed in germinated spores, with maximum-intensity projections and 3D-rendered views providing evidence of dsRNA accumulation along germ tubes and within spore bodies (Figures 6C-G). These findings demonstrate that spores of DM-causing pathogens can actively take up exogenous dsRNA, which subsequently localizes and moves along germ tubes during spore germination.

**Figure 6.**
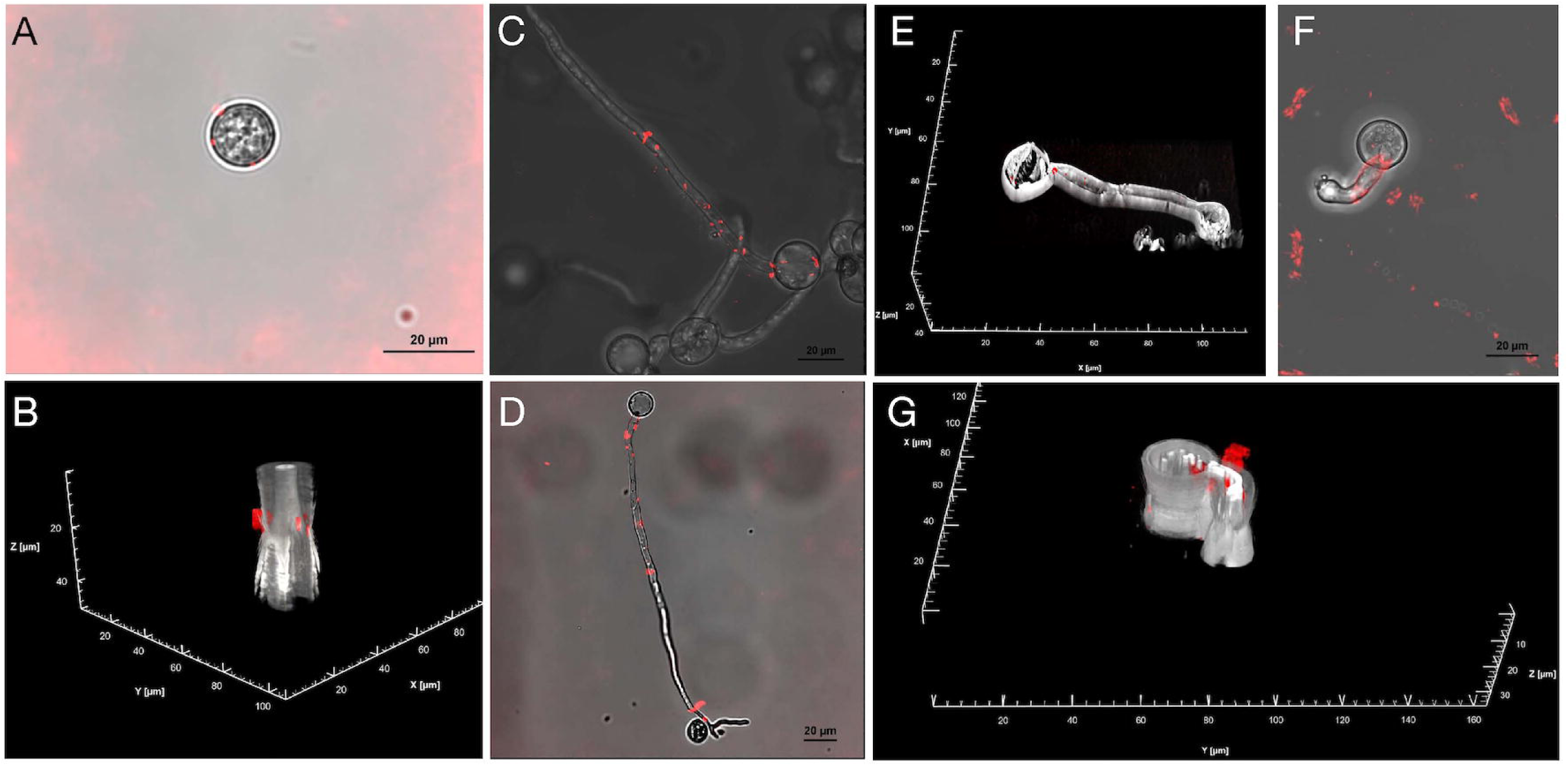
Internalisation of Cy-5 labelled SS-dsRNAs by downy mildew spores. *Hpa* and *Pvp*, were mixed with Cy-5 labelled dsRNA and examined with a Nikon Ti2-E AX NSPARC laser point-scanning confocal system 6h after incubation. In all images, Cy-5 labelled dsRNA is color-coded in red and superimposed over images of spores captured simultaneously with a transmitted detector mounted in the diascopic light path using 640nm laser (gray color-coded). **A)** *Hpa* spores, view from a single plane of a Zstack (77 slices @ 0.659um each step), **B)** Same spores as A but 3D cropped rendered view. RNA particles (red coloured) inside spores, **C)** Germinated *Pvp* spores view from a maximum intensity projection of a Zstack (61 slices @ 0.589um each step), **D)** Germinated *Hpa* spores in a single plane view from a Zstack (77 slices @ 0.659um each step), **E)** Germinated *Pvp* spores, view as a 3D cropped rendered projection of a Zstack (69 slices @ 0.589um each step). RNA particles imaged with NSPARC super resolution detector, **F)** Germinated Pvp spores, view from a maximum intensity projection of a Zstack (61 slices @ 0.589um each step), **G)** Same spore as F but in a 3D cropped rendered view of the Zstack. All images were captured with either a Nikon Plan Apochromat Lambda D 20X (0.8NA) or a Nikon Lambda S LWD water immersion 20X (0.95 NA) objective.

### SS-dsRNA-mediated gene silencing is sustained over time

To assess the efficacy and longevity of SS-dsRNA-mediated gene silencing, we targeted *HpaT809066*, a 1575 bp gene encoding a 525-amino-acid protein. Analysis using InterProScan (Blum et al., 2024) and UniProt (UniProt Consortium, 2025) revealed that this protein contains a signal peptide and a protein disulfide-isomerase domain, suggesting a potential role in pathogen virulence or host interaction. A 30 bp SS-dsRNA was designed to silence *HpaT809066*, and its expression was quantified using qRT-PCR at 4, 7, 10, and 11 dpi.

As shown in Figure 7, level of gene expression remained high in untreated samples (no SS-dsRNA), whereas SS-dsRNA-treated samples exhibited a significant and consistent reduction in expression at all time points. The most pronounced decrease occurred at 4 dpi (94% reduction), followed by reductions at 7 dpi (89%), 10 dpi (80%), and 11 dpi (53%). The observed suppression was statistically significant (****, p < 0.0001), confirming the robustness of SS-dsRNA-induced gene silencing.

**Figure 7.**
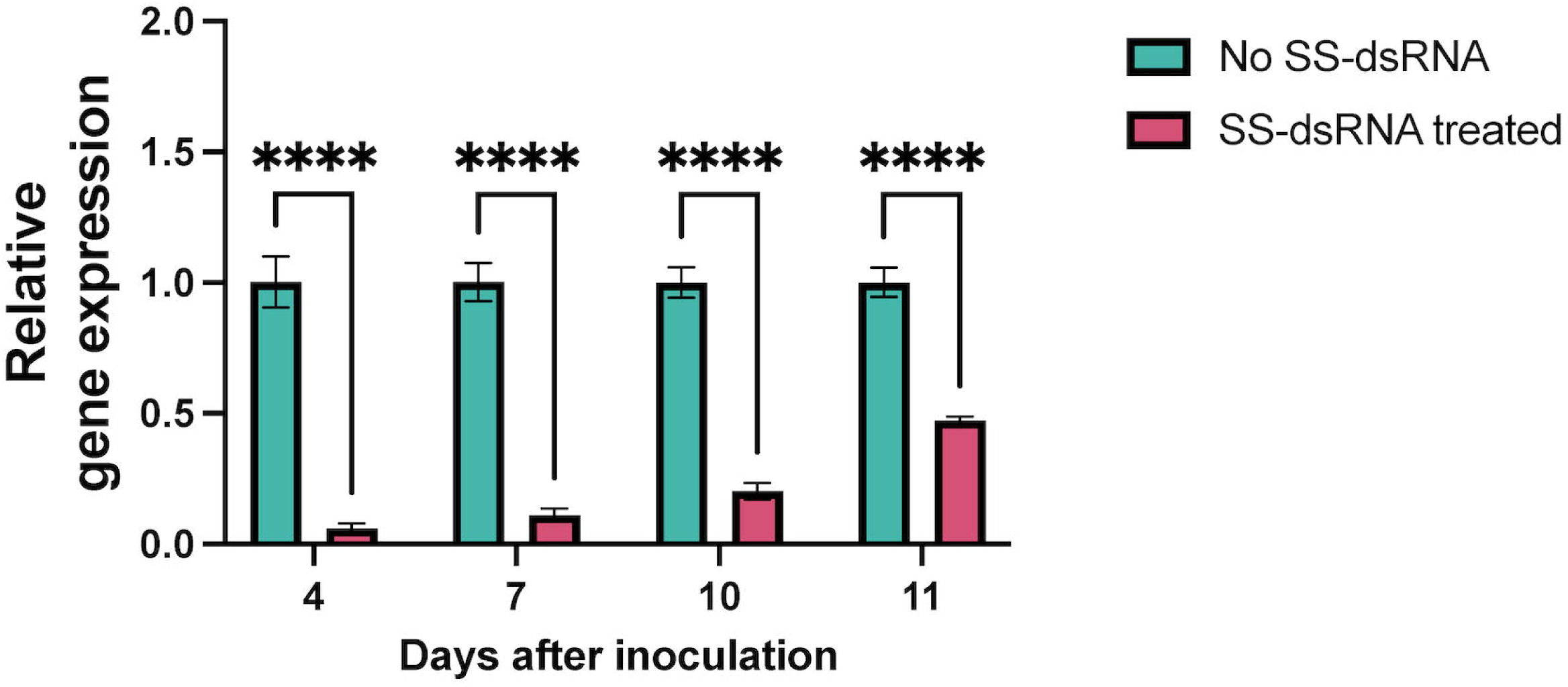
SS-dsRNA-mediated silencing of *HpaT809066* reduces gene expression over time. *H. arabidopsidis* (*Hpa*) spores were treated with 5 μM SS-dsRNA, while untreated samples served as controls. Gene expression levels were assessed by RT-qPCR at 4, 7, 10, and 11 dpi. A significant reduction in gene expression was observed in SS-dsRNA-treated samples compared to untreated controls at all time points (\*\*\**, p < 0.0001).* Statistical significance was assessed using a two-way ANOVA, comparing SS-dsRNA-treated and untreated samples. Values marked with an asterisk (*) indicate significant differences. Bars represent the standard deviation of three independent samples.

## Discussion

We used three host-DM pathosystems—*Arabidopsis-Hpa*, Pea*-Pvp*, and Lettuce*-Bl*— to investigate factors influencing dsRNA-mediated pathogen control. Through *in vitro* spore germination assays, *in vivo* sporulation assays, and gene expression analyses, we demonstrated that exogenously applied SS-dsRNAs targeting the *CesA3* and *BTUB* genes effectively reduced gene expression, thereby inhibiting spore germination and plant infection. The uptake efficiency of dsRNAs by spores was influenced by dsRNA length, while dsRNA concentration significantly affected spore germination and infection rates. Additionally, we demonstrated the feasibility of simultaneously targeting multiple genes using SS-dsRNAs. Using confocal microscopy, we provided direct evidence that *Hpa* and *Pvp* spores can take up fluorescently labelled SS-dsRNAs. Additionally, we showed gene silencing could be sustained for at least 11 days within *Hpa*. These findings highlight the potential of SIGS as an effective strategy against DM pathogens.

The oomycete cell wall is primarily composed of β-glucans and cellulose (Raaymakers & Van den Ackerveken, 2016), with cellulose biosynthesis mediated by cellulose synthase (CesA) enzyme complexes (Grenville-Briggs *et al*., 2008). The CesA3 subunit, essential for cellulose biosynthesis, is a key target for oomycete-targeting compounds such as carboxylic acid amides (CAAs), which disrupt cell wall formation in oomycetes (Blum *et al*., 2012). CAAs exhibit activity against oomycetes, including *P. infestans, P. capsici,* and *Plasmopara viticola* (Blum & Gisi, 2012), although specific amino acid changes in CesA3 can confer insensitivity (Blum et al., 2012).

Microtubules, composed of α- and β-tubulin subunits, are crucial cytoskeletal components involved in cell division and plant-microbe interactions (Hardham, 2013) (Piquerez, 2011). β-tubulin has been targeted in fungicides such as methyl benzimidazole carbamates (MBCs), which inhibit tubulin polymerization but are ineffective against oomycetes (Vela-Corcia *et al*., 2018). However, β-tubulin inhibitor zoxamide is effective against oomycete pathogens, including *Phytophthora* and *Pythium spp.* (Cai *et al*., 2016).

Previous studies have demonstrated the role of *CesA3* and *β-tubulin* genes in oomycete pathogenicity through mutant analyses and fungicidal treatments. However, our results demonstrate that these genes can also be effectively targeted using dsRNA-based biopesticides. Until now, many pathogen genes have been targeted using SIGS-based methods (Bilir *et al*., 2022). Similarly, dsRNA shows promise for pest management by silencing target genes. For example, dsRNAs targeting tubulin genes (*Msalpha-tubulin* and *Msbeta-tubulin*) in *Mythimna separata*, the oriental armyworm effectively reduced target gene mRNA levels, leading to reduced body weight and increased mortality phenotypes (Wang *et al*., 2018). In our study, silencing of *BTUB* genes in three DM-causing pathogens not only supports tubulin as a promising candidate for SIGS but also highlights that pathogen-specific regions within these genes can enhance the specificity of dsRNA targeting. A unique aspect of our study is the use of 30 bp SS-dsRNAs to target both the *CesA3* and *BTUB* genes in *Hpa*, *Pvp*, and *Bl*, emphasizing the importance of allelic variations within the targeted regions. For instance, while sporulation results for *Hpa* and *Pvp* were similar, those for *Bl* differed, likely due to the dsRNA mismatches in the target region.

The term “RNAi” describes the overarching mechanism of gene silencing mediated by dsRNAs through the association of sRNAs, such as siRNAs and miRNAs, with Argonaute effector proteins (Chou *et al*., 2024). Initially, RNAi was considered to be a sequence-specific mechanism requiring perfect complementarity between the siRNA guide strand and the target mRNA sequence (Elbashir *et al*., 2001). However, subsequent studies revealed that the specificity of siRNA-mediated silencing depends on both the position and identity of nucleotide mismatches (Qiu *et al*., 2005). For instance, certain mismatches, such as A:C pairs and G:U wobble base pairs, are well tolerated, enabling silencing efficiency comparable to fully matched sequences (Du *et al*., 2005). This tolerance to mismatches might explain the observed effects in our spore germination and sporulation assays, where mismatches at the target region could have been accommodated.

Recent studies have highlighted the importance of dsRNA length and concentration in determining the efficacy of RNAi-mediated silencing (Höfle *et al*., 2019; Baldwin *et al*., 2018a; Koch *et al*., 2019; He *et al*., 2020; Soumya *et al*., 2024). Studies with longer dsRNAs (400–1500 bp) on *Fusarium graminearum* have shown greater effectiveness due to their ability to generate a more diverse pool of siRNAs (Höfle et al., 2019). However, researchers working on *Sclerotinia sclerotiorum* have argued that longer constructs (>2 kb) are prone to degradation, reducing their applicability under field conditions (Soumya *et al*., 2024). In the present study, we used different length of chemically synthesised, IVT- or *E. coli-*produced dsRNAs of different lengths and determined their impacts on *Hpa* and *Pvp* spore germination and sporulation on their host plants. While 21-25 bp did not inhibit spore germination entirely, 30, 50, 75 and 100 bp long dsRNAs inhibited spore germination, indicating that spores could take up dsRNAs as long as 100 bp. When we pushed beyond the limits of chemically synthesis to longer fragments using *E-coli* produced dsRNA fragments *Hpa-CesA3* (285 bp) and *Hpa-BTUB* (273 bp), we did not see total inhibition of spore germination. However, digesting these long fragments with RNase III allowed full inhibition of spore germination. This clearly indicates that DM-causing pathogen spores are limited in their uptake of dsRNA in respect of its length. We also note the importance of properly choosing control dsRNA, as they can show off-target effects. For example, when we used *E. coli*-produced dsRNA from E. coli *GUS* gene (255 bp) as a negative control, we observed reduced germination and after digestion with RNase III, we observed full inhibition of spore germination.

Efficient dsRNA uptake occurs in the fungal plant pathogens including *Botrytis cinerea*, *Sclerotinia sclerotiorum*, and *Verticillium dahliae*, but no uptake in the fungal pathogen *Colletotrichum gloeosporioides* and weak uptake in the fungus, *Trichoderma virens* (Qiao *et al*., 2021). Interestingly, with the oomycete plant pathogen *Phytophthora infestans*, RNA uptake was limited and varied across different cell types and developmental stages (Qiao et al., 2021). Similarly, studies on wheat-*Zymoseptoria tiritici* pathosystem revealed that *Z. tritici* is incapable of dsRNA uptake (Kettles *et al*., 2019). In some cases, comparisons of dsRNA fragments targeting different genes have shown that shorter dsRNAs (∼250–350 bp) are less effective in generating diverse siRNAs but can achieve gene-specific effects under certain conditions (Baldwin *et al*., 2018b). In addition, these uptake efficiencies are linked to the functionality of fungal RNAi pathways, including *Dicer* and *Argonaute* genes, which have been shown to control RNAi effectiveness (Gaffar *et al*., 2019; Werner *et al*., 2019). However, longer dsRNAs are not always preferable, as they may increase the likelihood of off-target effects in the host plant. This is underscored by our observation of chlorosis and necrosis in *Arabidopsis* and chlorosis with leaf curling in pea when longer dsRNAs were used to target *CesA3* genes in sporulation assays. Thus, design of dsRNAs should consider trade-offs between dsRNA lengths and potential off-target effects.

The concentration of applied dsRNA is crucial and the optimum varies between pathosystems. Moderate concentrations (∼30–100 ng/µL) typically achieve optimal silencing, whereas higher concentrations (>200 ng/µL) may induce non-specific stress responses or degradation (Qiao *et al*., 2021). Conversely, concentrations below 10 ng/µL are generally ineffective for pathogen gene silencing (Qiao *et al*., 2021; Koch *et al*., 2016). In our studies, we tested 10, 100, and 200 ng/µL (equivalent to 1, 5, and 10 µM) on *Hpa* and *Pvp* spore germination and infection. We found 100 ng/µL to be optimal for *Hpa* and *Pvp* spore germination assays. However, for sporulation, 100 ng/µL was optimal for the *Arabidopsis-Hpa* pathosystem, whereas 200 ng/µL was required for the Pea*-Pvp* pathosystem. These findings highlight the necessity of optimizing dsRNA concentration for each pathosystem.

Recent studies highlight the efficacy of dsRNA constructs that target multiple genes in fungal pathogens. For example, simultaneous silencing of multiple virulence and growth-related genes in *Fusarium graminearum* (including *FgSGE1*, *FgSTE12*, and *FgPP1)* enhanced wheat resistance to Fusarium head blight (Wang *et al*., 2020) (Yang *et al*., 2021). Similarly, targeting ergosterol biosynthesis genes in *Botrytis cinerea* effectively suppressed fungal proliferation (Danielle *et al*., 2021). In our multiplexing experiments, targeting specific individual genes did not inhibit spore germination. However, simultaneous silencing of two or three genes resulted in complete germination inhibition. Likewise, sporulation assays showed a significant reduction when multiple genes were targeted together. These findings reinforce the power of multiplex approaches in disrupting complex virulence networks, highlighting their potential for pathogen control across diverse pathosystems.

Several studies have demonstrated the uptake of fluorescently labelled dsRNAs in fungal and oomycete pathogens using confocal microscopy, confirming its localization within targeted cells. For example, *Cercospora zeina* spores were shown to internalize fluorescein-labelled dsRNA, leading to RNAi-mediated silencing of GFP expression and fungal viability reduction (Marais, 2024). Similarly, fluorescein-labelled dsRNA targeting the virulence gene *Tup1* in *F. oxysporum* confirmed dsRNA internalization, which correlated with reduced gene expression and fungal pathogenicity (Fan *et al*., 2024). Other studies, such as those on *P. infestans,* utilized confocal microscopy to visualize dsRNA uptake in sporangia, linking RNA uptake to suppression of critical genes involved in infection (Kalyandurg *et al*., 2021). Remarkably, the emerging use of nanosheets or other delivery enhancements has improved dsRNA uptake efficiency and stability under diverse environmental conditions, further broadening the applicability of such approaches (Chen *et al*., 2022). Structural barriers, such as the integrity of spore walls, are likely contributors to this variability, prompting researchers to explore additional delivery strategies, such as enzymatic pre-treatments or improved carriers (Qiao *et al*., 2021). Additionally, confocal microscopy studies have highlighted the importance of distinguishing dsRNA entry from surface adhesion, with methods like Z-stack imaging and colocalization analysis providing evidence for intracellular localization (Qiao *et al*., 2021). By combining fluorescence imaging techniques with functional RNAi assays, these studies have established strong correlations between dsRNA uptake, target gene silencing, and pathogen inhibition. For example, the fluorescence intensity of dsRNA within fungal cells has been linked to reduced expression levels of key target genes, phenotypic outcomes such as growth retardation, and suppression of plant disease symptoms (Marais, 2024; Fan *et al*., 2024; Danielle *et al*., 2021).

In our study, we used Cy-5 labelled SS-dsRNAs and checked their internalisation by *Hpa* and *Pvp* spores. Confocal microscopy clearly indicates that these fluorescently labelled dsRNAs can be detected in spores or germ tubes reinforcing the utility of confocal microscopy in validating RNAi delivery systems and contribute to a growing understanding of dsRNA internalization mechanisms in oomycetes.

The observed discrepancy between the inhibition of pathogen germination on glass slides and the persistence of sporulation on the host plant with some of the SS-dsRNA (Table 2) may arise from fundamental differences in the experimental environments and pathogen biology. *In vitro* germination assays provide a controlled system where dsRNA directly interacts with the pathogen under optimized conditions, potentially achieving higher effective concentrations and uninterrupted activity. In contrast, *in planta* assays introduce dynamic variables: host tissues may absorb or degrade applied dsRNA, reducing its bioavailability to the pathogen (Qiao *et al*., 2023). Additionally, the pathogen’s interaction with the plant cuticle, apoplastic enzymes, or host-derived nutrients could alter its physiological state, enabling partial evasion of dsRNA-mediated silencing (Islam *et al*., 2021). The pathogen might also employ alternative genetic pathways during later infection stages, such as sporulation, that are less reliant on the genes targeted during germination. Furthermore, microenvironmental factors (e.g., humidity, leaf microbiota, or plant immune responses) could stabilize the pathogen or diminish dsRNA efficacy (Ray *et al*., 2022). These findings indicate the necessity of optimizing dsRNA delivery mechanisms to enhance stability and uptake *in planta*, while highlighting the complementary value of both assays in elucidating gene function across distinct phases of the pathogen’s life cycle.

SS-dsRNA-mediated gene silencing in *Hpa* exhibited a sustained reduction in gene expression over an 11-day period, with the strongest suppression observed at 4 dpi (94% reduction) and a gradual decline to 53% by 11 dpi. This longevity aligns with previous studies indicating that RNA silencing duration depends on RNA stabilization strategies and pathogen-specific RNAi efficiency. Unlike naked dsRNA, which is prone to rapid degradation (Whisson *et al*., 2005; Binod *et al*., 2024), our SS-dsRNA exhibited prolonged activity, suggesting enhanced stability and uptake by *Hpa*. While fungal pathogens such as *Sclerotinia sclerotiorum and Botrytis cinerea* generally exhibit more efficient dsRNA uptake and prolonged silencing, the oomycete pathogen *Phytophthora infestans* show weaker RNA uptake, potentially explaining the decline in silencing at later time points (Han *et al.,* 2025; Qiao et al., 2021; Danielle *et al*., 2021). Additionally, gene-specific factors, such as the target sequence and its role in pathogen virulence, likely influenced silencing efficiency. The significant suppression observed here highlights SS-dsRNA as a promising tool for managing downy mildew, though further studies optimizing dsRNA stabilization and delivery could enhance its durability and field applicability.

In conclusion, our study demonstrates that exogenously applied SS-dsRNAs effectively silence targeted genes in DM pathogens, inhibiting spore germination and reducing plant infection. The uptake efficiency of dsRNA varies with length and concentration, highlighting key parameters for optimizing SIGS-based pathogen control. While *CesA3* and *BTUB* genes remain effective targets, pathogen-specific sequence variations influence dsRNA specificity and efficacy. Furthermore, our findings reveal limitations in dsRNA uptake by oomycete spores, emphasizing the need for precise design to balance efficacy and off-target effects. These insights advance the development of RNA-based biopesticides and provide a foundation for future translational applications in sustainable crop protection.

### Experimental Procedures

#### Plant lines

*Arabidopsis* mutant Ws-*eds1* (Parker *et al*., 1996) was used to maintain and test sRNAs with *Hpa* Emoy2 isolate. Marrowfat pea seeds were obtained from Church of Bures, https://churchofbures.co.uk, and used for pea-DM experiments. For lettuce DM experiments, lettuce cv. Gustav’s Salad was used, and seeds were purchased from local garden centre.

#### Downy mildew isolates and propagation

##### *Hpa* propagation

*Hpa* isolate *Emoy2* was maintained on *Arabidopsis* Ws-*eds1* plants. The 7-day-old infected seedlings were collected in cold sterile distilled water (SDW), gently vortexed, and filtered through a layer of Miracloth. The *Hpa* conidiospores were collected by centrifugation, washed twice with cold SDW and the conidiospore concentration released in SDW was adjusted to 5 x 10^4^ spores/mL using a haemocytometer. The 7-day-old *Arabidopsis* plants were inoculated with the obtained *Hpa* conidiospores by using spray atomiser. The inoculated plants were then placed in a plastic tray with a transparent lid, the edges sealed with tape to maintain humidity. The plants were grown at 16°C under a 12-hr photoperiod in growth cabinets. Sporulation was assessed 7 days post-inoculation (dpi) as described (Bilir *et al*., 2019).

##### *BI* propagation

*Bl-*Tid1 isolate was obtained from a natural infection on lettuce cv. Gustav’s Salad, grown at an allotment in Tiddington, Stratford upon Avon. The inoculum was washed off, single spored and subsequently maintained on the same cultivar. The inoculum was prepared in the same way as *Hpa*, adjusted to a concentration of 5 x 10^4^ spores/mL. The 7-day-old lettuce cotyledons were drop inoculated with 5-10 µL of inoculum onto each cotyledon. The inoculated plants were placed in a plastic tray, covered as described in Hpa, and grown under the conditions specified in *Hpa*. Sporulation was assessed 7dpi.

##### *Pvp* propagation

*Pvp* isolate DM3 was from the DM collection of Niab, United Kingdom. The infected pea plants were collected in cold SDW, gently vortexed and filtered through a layer of Miracloth. The spore suspension adjusted to 5 x 10^4^ spores/mL was used to inoculate 4-day-old pre-germinated pea seeds. The seedlings were immersed in the spore suspension for 30 mins with gentle shaking every 5 mins to ensure uniform inoculation and then immediately sown in a compost and grown at 16°C under 12-h photoperiod in growth cabinet. After 10 days, the inoculated plants were covered with a transparent lid for 2 days to aid the pathogen to sporulate.

### Generation of small dsRNAs

Small dsRNAs were designed using InvivoGen’s, https://www.invivogen.com/sirnawizard/design_advanced.php, siRNA wizard software or home developed MYCIsiRNA https://mycosirna.wp.worc.ac.uk/design.html software by adjusting the motif size to 30 nt, and small dsRNA duplexes were either obtained as chemically synthesized ribonucleotides from Merck (Gillingham, Dorset, UK) or GenScript (Oxford, UK). In addition, dsRNAs were also produced by *in vitro* transcription (IVT) in-house by using MEGAscript kit (Thermo Fisher, Waltham, MA, USA) according to the manufacturer instructions or obtained from RNA Greentech LLC (Texas, USA). Sequences of dsRNA duplexes used are given in the Supplemental Table 1. The sequences of the dsRNA duplexes and PCR primers used are provided in Supplemental Table 1.

### Production of dsRNAs in *E. coli*

Production and purification of dsRNA were carried out according to (Ahn *et al*., 2019) with some modification. Synthetic DNA fragments (G-blocks) encoding partial sequences (273 bp, 285 bp, 255 bp) of the *Hpa-CesA3* and *Hpa-BTUB* genes, and *E. coli* GUS gene as control, respectively, were ordered from Integrated DNA Technologies (IDT, Coralville, IA, USA) with added SacI and PstI restriction sites. Amplification of all G-blocks was done using Taq DNA Polymerase (New England Biolabs, Ipswich, MA, USA) along with gene specific primers. The PCR amplified *Hpa-CesA3, Hpa-TUB and GUS* DNA fragments and pL4440 plasmid vector with two inverted T7 promoters were digested with *SacI* and *PstI*. Digested products were then ligated together at 4°C overnight in the presence of T4 DNA ligase. The constructs were then transformed into *E. coli* HT115 (DE3), which lacks RNase III, using a standard transformation protocol. Positive transformants were selected at 100 µg/mL ampicillin and 12.5 µg/mL tetracycline on LB agar plates and verified by colony PCR. Single transformants of HT115 carrying the recombinant pL4440 vectors were inoculated into 4 mL of LB medium containing 100 µg/mL ampicillin and 12.5 µg/mL tetracycline, then incubated overnight at 37°C with shaking (190 rpm). A 500 µL aliquot of the overnight culture was transferred into 100 mL of LB broth containing the same antibiotics and incubated at 37°C with shaking to an OD_600_ of approximately 0.4. RNA transcription was induced by adding 1 mМ isopropyl β-D-1-thiogalactopyranoside (IPTG) (Thermo Fisher Scientific, Waltham, MA, USA) and incubation continued for a further 5 hours under the same conditions. The cultures were harvested for dsRNA extraction.

Total RNA was purified from 100 mL of a bacterial culture using the TRIzol™ Max™ Bacterial RNA Isolation Kit according to the manufacturer’s instructions (Thermo Fisher Scientific). The RNA pellet was dissolved in nuclease-free water and further treated with Turbo^TM^ DNase and RNase A to remove DNA contamination and single-stranded RNA contaminants. The dsRNA was further purified by adding chloroform/isoamyl alcohol 24:1, v/v (Sigma-Aldrich, St. Louis, MO, USA). The upper aqueous phase containing the dsRNA was transferred into a new tube, mixed with 7.5 M ammonium acetate and isopropanol, and centrifuged at 12,000 g for 30 minutes at 4°C. The dsRNA pellet was washed twice with 70% ethanol, air-dried, and resuspended in nuclease-free water. RNA yield and quality were measured using a Synergy H1 hybrid reader (BioTek, Winooski, VT, USA), while the RNA integrity was assessed by 1.2% agarose gel electrophoresis.

### Germination assays

*Hpa*, *Pvp* and *Bl* spores adjusted to 5 x 10^4^ spores/mL in SDW were mixed with small dsRNAs at a final concentration of 1 or 5 µM in an Eppendorf tube, then incubated on ice for 20 minutes. A total of 100 µL spore suspension mixed with or without dsRNA (control) was dropped onto the microscope glass slides with three biological replicates. Microscope glass slides were placed into the Petri dishes, which were covered with cling film to increase the humidity and placed into a plastic box. These boxes were then incubated with a 12 h light/12 h dark regime at 16 °C for 24 h. Spores were examined under a light microscope (Leica DM5500 B and Olympus CK-2) 24 h after incubation, and total and germinated spores were counted. The percent germination was calculated based on the control group representing only pathogen spores without dsRNA. The values were expressed as a percentage of control to eliminate variation encountered in the controls. Each experiment was repeated at least twice.

### Sporulation assays

A total of 100 µL of *Hpa*, *BI*, and *Pvp* spore suspensions, with or without short dsRNA, was prepared following the same procedure as in the germination assay. The only exception was for *Pvp*, where a final dsRNA concentration of 10 µM was used. Seven-day-old *Arabidopsis* seedlings (for *Hpa*) and lettuce seedlings (for *BI*), grown in plastic modules with uniform spacing, were used for inoculation. Three replicates were prepared for each pathogen, with 10-15 seedlings per replicate for Hpa and 3 seedlings per replicate for *BI*. These seedlings were drop-inoculated with the spore suspension, applying 1 µL to each cotyledon for *Hpa* and 10 µL to each cotyledon for *BI*.

For *Pvp*, 4-day-old pre-germinated pea seeds were sown in compost, kept under the same conditions as during propagation. After 5 days, the pea seedlings were inoculated by placing 10 µL of spore suspension inside the newly expanded leaves. All plants (Arabidopsis, lettuce, and pea) were maintained under the same conditions as during propagation. Sporulation was assessed at 7 dpi for *Hpa* and *BI*, and at 8 dpi for *Pvp*.

For conidiospore quantification, infected seedlings from each replicate were collected, with the roots discarded. For *Hpa*, ten seedlings per replicate were placed into an Eppendorf tube containing 250 µL of SDW, while for *BI*, three seedlings per replicate were placed into an Eppendorf tube containing 1 mL of SDW. For *Pvp*, three infected seedlings per replicate were placed into a Falcon tube containing 40 mL of SDW. The samples were gently vortexed to release the spores into suspension, then filtered through Miracloth. The conidiospores were collected by centrifugation and resuspended in 1 mL of SDW for assessment. Spore counts for all pathogens were performed by counting the number of spores present in all four corner grids of a haemocytometer. The sporulation percentage was calculated relative to the control group. Each experiment was repeated twice.

### Multiplexing of SS-dsRNA

Three *Hpa* genes, *HpaT802064*, *HpaT802452*, and *HpaT803108,* were selected for the multiplexing experiments. SS-dsRNAs targeting these genes were combined with the conidial suspension following the same protocol used for germination and sporulation assays. Instead of applying individual treatments, the SS-dsRNAs were tested singly, in pairs, or as a triple combination, with each SS-dsRNA in the mixture maintained at a final concentration of 5 µM. Each experiment was repeated twice.

### Confocal microscopy to detect dsRNA internalisation

To assess dsRNA uptake by *Hpa* spores, a 30-nt SS-dsRNA was designed based on *HpaT802307* (sense: 5’-GUCUGAAAGGAGCCGAGAUUGACCUGAUUA-3’; antisense: 5’-UAAUCAGGUCAAUCUCGGCUCCUUUCAGAC-3’) and synthesized with a Cy-5 fluorophore attached to the 5’ end. *Hpa* or *Pvp* conidiospores were suspended in SDW at a concentration of 5 × 10⁴ spores/mL and mixed with 1 µM labelled SS-dsRNA in an Eppendorf tube. The mixture was incubated on ice for 20 minutes to facilitate interaction between the spores and the dsRNA.

After incubation, 30 µL of the spore-dsRNA mixture was pipetted onto microscope cavity glass slides. A control group containing spores without SS-dsRNA was prepared similarly. The slides were placed in Petri dishes, covered and incubated under the conditions specified in the germination assay. Spores were examined 6 hours post-incubation using a Nikon Ti2-E AX NSPARC laser point-scanning confocal system.

### Isolation of total RNA

Initially, we optimized the timing for RNA isolation for each pathosystem to perform gene expression analysis. For all three pathogens, 5 x 10^4^ spores/mL were used for inoculations. Gene expression was analyzed using a combination of in vitro and in planta assays. For *BTUB* gene expression analysis in *BI* and *Pvp*, spores were mixed with 5 µM siRNA in a 1.5 µL Eppendorf tube and incubated under a 12 h light/12 h dark cycle at 16°C for 24 hours. After incubation, the mixture was centrifuged at 14,000 rpm for 5 minutes, and the resulting pellet, containing conidiospores, was used for total RNA extraction. For *BTUB* and *CesA3* gene expression analysis in *Hpa*, and *CesA3* in *BI* and *Pvp*, 5 µM dsRNAs were used for *Hpa*, while 10 µM were used for *Pvp* and *Bl*. For RNA extraction, four-week-old *Arabidopsis* plants were used. A total of six leaves were marked for each plant and each leaf was drop inoculated with 20 µL *Hpa* spore suspension with or without dsRNA. At 3 dpi, two leaves representing a biological replicate were collected in an Eppendorf tube and snap-frozen in liquid nitrogen before storage at −80 °C until RNA extraction. Similarly, lettuce and pea plants were inoculated with respective pathogen spore suspensions with or without dsRNA as described for sporulation assay and leaves were collected at 4 dpi and kept at −80°C until RNA isolation. Three biological replicates were set up for each treatment. Total RNA was extracted by Qiagene RNeasy Plant Mini Kit (Qiagen, UK), Zymo Quick-RNA Plant Miniprep Kit (R2024) and Bioline Isolate II RNA Plant Kit (BIO-52076) according to the manufacturer’s instructions. RNA concentration and quality were evaluated by NanoDrop 2000c spectrophotometer (Thermo Fisher Scientific, USA).

### Quantitative real-time PCR analysis

To determine whether targeted genes were silenced by the application of dsRNA, qRT-qPCR was performed using Superscript III Platinum SYBR Green One-Step qRT-PCR Kit (Thermofisher Scientific, Paisley, UK). Primer sets used for amplification are listed in Supplemental Table 2. Reactions were performed on Roche Light Cycler 480 II PCR machine using the following touchdown cycling conditions: initial incubation at 45°C for 10 minutes, followed by 40 cycles of denaturation at 95°C for 5 sec, annealing starting at 68°C for 10 seconds (with the annealing temperature decreasing by 0.8°C per cycle from 68°C to 60°C over the first 10 cycles), and extension at 72°C for 5 seconds. Each qPCR reaction contained 5 µL of SYBR Green PCR Master Mix, 0.3 µL of each 10 µM primer, 0.1 µL of reverse transcriptase, 0.2 µL RNA-inhibitor, 1 µL RNA (at 70 ng/ µL) and DEPC water for a final reaction volume of 10 µL. Melt curve analysis was performed to confirm amplification specificity. Data analysis was performed using the 2^−ΔΔCt method, with gene expression normalized to the actin gene of the respective pathogen (*Hpa, Pvp or Bl).* For each sample, the mean of three replicates was used to calculate relative change in mRNA levels.

### Longevity test

Seven-day-old *Arabidopsis* seedlings were drop inoculated using the same procedure as described for the sporulation assay, with the exception that additional seedlings were prepared for each time point. For each of the three biological replicates, samples were collected at 4, 7, 10, and 11 dpi. Seedlings were immediately placed into Eppendorf tubes, flash-frozen in liquid nitrogen, and stored at −80°C until RNA extraction.

### Bioinformatics

Primers were designed either using the Primer3 web tool (https://primer3.ut.ee) or using Geneious (v10.0) (Kearse *et al*., 2012). We used the EnsemblProtist (Kersey *et al*., 2016) and InterPro (Quevillon *et al*., 2005) databases to identify candidate *Hpa*-*BTUB* gene. Reciprocal BLASTN and BLASTX (Altschul et al., 1997) were used to perform similarity-searches of nucleotide and amino acid sequences, respectively, between *Hpa, Pvp and Bl* genes. Multiple sequence alignments were generated using MUSCLE v3.8.31 (Edgar, 2004) and visualised using Jalview 2.11.4.1. (Waterhouse *et al*., 2009).

### Statistical analysis

All the graphs and statistical analyses were performed with GraphPad prism software for Mac (San Diego, CA, USA).

### Accessions

Coding sequences of *Pvp-CesA3* and *Pvp-BTUB* can be accessed under the accession numbers PV324775 and PV324774, respectively.

## Author contributions

MT and JMM conceived the research idea and designed the experiments with DG; DG, EO, GÜ and SMS performed lab-based experiments; AW, TWacker and DS provided bioinformatic support while TW provided plant pathology support. MT, YH, TWacker, DS, JMM and DG analysed the data. All authors contributed to the writing and review of the manuscript and approved the final version for submission.

## Data availability statement

The data that support the findings of this study are available from the corresponding author on reasonable request.

## Acknowledgements

We would like to express our sincere gratitude to The Nikon Imaging Centre at King’s College London, particularly Dr. Michelangelo Colavita, Advanced Imaging Specialist, for his invaluable assistance with confocal microscopy. **We also acknowledge John McDowell’s significant contributions to this field and remember him with gratitude**.

## Supplemental Figures and Tables

**Supplemental Figure 1. Multiple sequence alignment of *CesA3* genes from oomycetes and model organisms.** The black shading indicates the location of the sequence matching 30 bp long *Hpa-CesA3* Common-SS-dsRNA on the *Hyaloperonospora arabidopsidis* sequence. The blue shading indicates CesA3-*E. coli*-produced dsRNA. The black shading indicates *Hpa-CesA3* Specific SS-dsRNA. Sequences were aligned using MUSCLE v3.8.31. The accession numbers and sequence coordinates of the aligned oomycete sequences are:

- *Hyaloperonospora arabidopsidis* Emoy2 (Hy_arabidopsidis) HpaG810051
- *Pernonospora viciae* f. sp. pisi (Pe_v_pisi) this study
- *Phytophthora sojae* (Ph_sojae) RefSeq:XM_009535833.1
- *Phytophthora parasitica* (Ph_parasitica) RefSeq:XM_008915779.1
- *Phytophthora infestans* (Ph_infestans) RefSeq:XM_002897169.1
- *Phytophthora cinnamomi* (Ph_cinnamomi) RefSeq:XM_067930250.1
- *Bremia lactucae* (B_lactucae) RefSeq:XM_067963730.1
- *Plasmopara halstedii* (Pl_halstedii) RefSeq:XM_024717371.1 The alignment has been trimmed at the 3’-end.

**Supplemental Figure 2. Supplementary File 1. Multiple sequence alignment of *tubulin* genes from oomycetes and model organisms.** The red shading indicates the location of the sequence matching *Hpa-BTUB* SS-dsRNA. The blue shading indicates the location of the sequence matching *Hpa-BTUB E. coli*-produced dsRNA. The accession numbers and sequence coordinates of the aligned oomycete sequences are:

- *Bremia lactucae* (Br_lactucae) RefSeq:XM_067962673.1 (1-1290)
- *Hyaloperonospora arabidopsidis* Cala2 (H_arabidopsidis_Cala2) GenBank:LKIA01000455.1
- (10041-11330)
- *Hyaloperonospora arabidopsidis* Emoy2 (H_arabidopsidis_Emoy2) GenBank:ABWE02003170.1 (1637-2926)
- *Hyaloperonospora arabidopsidis* Noks1 (H_arabidopsidis_Noks1) GenBank:LLKM01000460.1 (23019-21730)
- *Hyaloperonospora brassicae* (Hy_brassicae) GenBank:SZZI01000768.1 (16911-15622)
- *Pernonospora viciae* f. sp. Pisi (Pe_v_pisi) This study
- *Peronospora effusa* (Pe_effusa) GenBank:CP090803.1 (3022079-3023368)
- *Phytophthora agathidicida* (Ph_agathacida) GenBank:CP106975.1 (4903626-4904915)
- *Phytophthora cinnamomi* (Ph_cinnamomi) GenBank:U22050.1 (280-1569)
- *Phytophthora infestans* (Ph_infestans) RefSeq:XM_002908737.1 (87-1376)
- *Phytophthora plurivora* (Ph_plurivora) GenBank:CP125268.1 (4846179-4847468)
- *Phytophthora ramorum* (Ph_ramorum) RefSeq:XM_067894435.1 (109-1398)
- *Phytophthora sojae* (Ph_sojae) GenBank:CP155031.1 (1172078-1173367)
- *Plasmopara halstedii* (Pl_halstedii) RefSeq:XM_024718186.1 (202-1490)
- *Pythium aphanidermatum* (Py_aphanidermatum) GenBank:MK752999.1 (1-1290)

The accession numbers and sequence coordinates of the aligned model-organism sequences are:

- *Arabidopsis thaliana* RefSeq:NM_123801.2 (157-1445)
- *Caenorhabditis elegans* (Worm) RefSeq:NM_077184.8 (3-1291)
- Human RefSeq:NM_178014.4 (156-1445)
- *Saccharomyces cerevisiae* (Yeast) GenBank:CP029160.1 (1556403-1557690)
- Zebrafish RefSeq: NM_198809.2 (112-1400). The alignment has been trimmed at the 3’-end.

**Supplemental Figure 3.** dsRNA fragments produced in *E. coli* targeting the *Hpa-CesA*3 (1), *Hpa-BTUB* (2), and *GUS* (3) genes.

**Supplemental Figure 4.** H*p*a spore germination assays using *E. coli*-produced dsRNA. (A) untreated control spores, (B) spores treated with *GUS* dsRNA, (C) spores treated with *CesA3* dsRNA, and (D) spores treated with *BTUB* dsRNA.

**Supplemental Table 1.** Nucleotide sequences of short, IVT- or E. Coli-produced dsRNAs used in this study.

**Supplemental Table 2.** Primers used in this study and their sequences.

**Supplemental Table 3.** Genes targeted in concentration and multiplexing assays and their possible function.

