## Supplemental Figure 1 for "Advancing RNAi-Based Strategies Against Downy Mildews: Insights Into dsRNA Uptake and Gene Silencing": Supplemental Figure 1.html

Launch in Jalview  

{"seqs":[{"start":1,"name":"Hy\_arabidopsidis/1-3370","end":3370,"id":"598887649","seq":"ATGGTGCTGACGGGCGCGGGCGTAATCGCCTCGGTCGTGGGCATCCTCGGCGGCCTGTCGCTTTCATGCGGTGGCTGGTCGTCGCTCTCCCTCGGCGCGCGCTCGCTCTTTGTCACAACGCAGTTCCTCTCGGCCTTTGCCAT-GGGATTTGTCATTGCGTTTACCGCAATCGTCTCGCTCAGCGACACGAACGAATGGGTCGCGCTAGCGGCTGGCGGCGGCGCCGGTTTCTTGATCGCGCTCATTGTCGGCTTCATGACTCTCTGTGGCCCGTACATCCTCATCTTTGCCACGGGCGGCATCGTTGCGTCGTACCTGCTCCTCATTGACGCCTACAACGGCATCAACGTCTTCCCCGCGGACAATCAACTCGCCCGGCAGGAGTTTGTCATTGCCTTTATGATCATCTTTGAGCTCGTGTGCTGCTCGAGCTCCAAGTCGCTCGAGCTCGAGAACCACCGCATCAAGTACGTCGTGTTCTCGGCCATTACGGGCGGCTGGCTCGCTGCCGACGGCGTCTCGCGCCTCATTGACTCGAGCGCTGTGCTCTCGGACGTGGCGTACCAGTCGATTCAAAATGGCGGCACAGCTGCGCTCAAGGGCATGGACGCTGGCTCGCAGACGCTCATGTTTATCATTTGGGCCGCTGTCGTCGTCATTGGCGGCCTCAATCAGCTCTCGATGCGCTGGGGACTCATGTGCTACAACCGCGTGGGCGCCCACGCACAGCTCGGACCCGTGGAGGAGCAGCTACCGGAGCTGCCCACGGGTGCGACGCTCCCGGCGCAGACGCTCACGGAGCGCGTGCGTCTCGTCTGTGAGAACTGCTTTGCGACTGTGCCCAGCGGCACGGCATTCTGCACGGAGTGTGGCGAAGCCATGCCGTCGGAAGACGCGAACCCAGACGTGAGCATCTCACAGGCGCAGATGCCGTCGGTCACGATGAACAGTAAGAGCCAAGCGCCCGAGCGCTGGCAGCAAGTGCCGCATCGCACGTACCTCAGTACGACGTCGTTCGTGGACCCGAAACACGCGAAAGAAGGCGGTGTGAGCATGAAGGACAATGGCCGCAGTATCCGCTTTATGGACGGGGGAGTCCAAGGCCCAGACGGCAAGATGGGGCAGTATAACGACTCGATTGCTGGTGTCCGGAACTACTACGAGCCGTCATTTCGGTCGTTTGCCATGTCGACGTATTCGATTGCGAATCGTGCGGCTGAGCCCGTTGAGACGCCGAATATCCGCAAGTACAAGATGTCGGGCAGTGGCATGTTCCACGTGTTTTACTTTGGCACGGCGGCTACGGGTATCATTTGGCTGTATTACTTGACGACGATGTACCCGCAGCAGTACTTTTGTGACCATGCGCGACCAACGTTACCGTGCGACATGTTGCCGGCCAGTGAAACGGAAGGCTGTTTCAGTTCGACGGTCAACTTTGACGCGGACTCGGGTGATGGATACTGTATCCGTAACGTGCCGTTCATGTCGTGGATCATGTACGCTATGATGATTTTCAGTGAGTTTCTCAATTTCTTTCTGGGACTGCTGTTCAACTTCAGTATGTGGCGTCCGATTCGTCGTGGAGCGCGGTACATGAACGATTTCAAGCCGCCTATTCCAAAGGAACAGTGGCCGACGGTCGATATCTTTTTGTGTCACTACATGGAACCGGTCACGGACTCGATGCAGACACTGAAGAACTGTCTTTCACTCCAGTACCCGCCCGAGTTACTGCACATCTTTGTGCTGGATGATGGTTACACCAAGTCGGTATGGGACGCGAACAACCACTTTAAGGTGACGGTGAACACTAAGGTGATTGAGATTTGTGGTGATTTGCGTGGTGACCTCGCTCGGCTCATGCACGAGCGTGTGGTTGGCCCTGTGCAGGACGACCAAAGCTTGAAGACGTGGCGTCGCCAGCACAGCTCTGTCCGTGAACTCCGCAAGGAGGGTGGCAAGGGCGTGCAGCGTCGTGACTGTGCTGTTGGCTCGCTTTCGGACGACTACGACTACCGTGACCGTGGTATCCCTCGTGTAACGTTCATCGGTCGTATGAAGCCCGAGACGCACCACTCCAAGGCTGGTAACATCAACAATTGCTTGTTCAACGAAGGTGCCGACGGCAAGTATTTGCTGATTCTGGATAACGACATGAAGCCGCATCCGAAGTTTTTGCTTGCGGTGCTGCCGTTTTTCTTCTCGGAAGGCGAGGCTGTTGACGGTGGAGGACGTCAGTACAGTGACGACATTTCGTGGAACCAGGTGTCATACGTGCAGACTCCTCAGTACTTCGAGGACACGCCGCAGTTGACTATCATGGGTGATCCGTGTGGCCACAAGAATACCATTTTCTTCGACGCTGTGCAGTGTGGTCGTGATGGTTTCGACTCTGCAGCTTTTGCCGGTACGAATGCTGTTTTCCGTCGTCAAGCTTTCGATTCGATTGGTGGTATACAGTACGGCACGCAGACAGAAGATGCCTTTACGGGTAACGTGCTGCACACTTCTGGTTGGGACTCGGTGTATTTCCGCAAGGATTTCGAGGGCGATGCCAAGGACCGTATCCGCCTATGCGAAGGTGCCGTTCCCGAGACGGTGGCAGCTGCCATGGGTCAGAAGAAGCGTTGGGCAAAGGGTGCTGTGCAGATTCTGCTGATGAAAAATGAGAGTGAAGTCGACCCGGACTGGCGTCCACCGCGTGTGCCTGCCCCGGACCCGAAGCCGTCTCTTACGTTCCCGCGTAAGATGTTCTTCTACGACTCGGTGCTGTACCCGTTCGGTTCAATTCCTGCCTTGTGTTACGTGGCAATCGCTGTCTACTACTTGTGCACGGGAGACGCTCCGATCTACGCTCGTGGAACCAAGTTCCTGTACTCTTTCTTGCCCGTGACGTTCTGCCGTTGGGTGCTGAATCTGTTGGCGAACCGCGCTGTTGATAACAACGACGTGTGGCGTGCGCAGCAGACATGGTTTTCCTTCTCGTTCATTACGATGATGGCAATTTTTGAGGCCATCCAGGCGCGCTTGACGGGTAAGGACAAGTCATGGGCAAACACGGGTGCTGGTCAAAAGACGTCGTGGACGGAAATCCCAAACGTGCTCTTCTTTTTCACGCTGCTGTTCAGTCAACTCGTGGCGCTCATTCGTTTCTTCGAGTACGAGAACGCAACGAACCCGTGGAACTACGTGTCTGCGATGTTCTTTGGCTTTTTCGTCATGAGCCAGTTCTACCCCATGGTCAAGATGAGTATTACCGAGTACTGTGGTTGGGACCATACTGCTGCCACCTTCACGGCGAACGTGTTTGGTTCGCTGCTGGTGGTCTACATCGTGGTCTTCGTGCAGCTATGGCAAGTGTACTACG","order":1},{"start":1,"name":"Pe\_v\_pisi/1-3370","end":3370,"id":"78344647","seq":"ATGGGGCTCACTGGCGCTGGCATAATCGCCTCCGTGGTTGGCATCCTGGGTGGATTGTCGCTATCTTGTGGCGGTTGGTCGTCACTCTCTCTCGGTGCACGCTCTCTCTTTGTCACGACACAGTTTCTGTCTGCCTTTGCCAT-GGGTTTTGTCGTGGCTTTTTCTGCCATTGTTTCTCTGACCGACACTAACGATTGGGTCGCTGTTGCTGCCGGTGGAGGTGCAGGTTTCGTCGTTGCACTTATCGTAGGCTTTGTGACACTCTTTGGACCGTACGTTTTGATTTTGGTCACAGGTAGCATCATTGCTTCGTACCTTTTGCTCTTTGACGCATTTAATGGTATTAATGTCTTCCCGGAAGACAATCAGCTGGCCCGTCAGGAATTTGTCATTTCTTTTATGATTATTTTTGCGCTCGTGTGTTGCTCCAACTCCAAAACAACGGAGCTGGAGAACCACCGCTTTAAATACGTCATCTTTTCGTCCATTACTGGTGGCTGGATGGCGGCAGACGGTATTTCCCGCTTGATCGACTCAGCAGCAGTGTTGTCTGATGTGGCCTATTTGTCCATCCAGGATGGCGGCAAAGCTGCTGTAGATGGTATGGACGCAGGCTCTCAGACGCTCATGTTTGTGATCTGGGGCGTAGTTGTCGTAATTGGCGGGCTCAACCAGCTCTCGATGCGCTGGGGACTCATGTGCTACAACCGAGTAGGAGCACACGCTCAGCTCGGCCCTGTCGAGGAGCAGTTGCCAGAGTTGCCTACCGGTGCGACGCTTCCTGCCCAGGTTATGACTGAACGCGTTCGCCTCGTATGTGAGAATTGTTTTGCGACTGTCCCTAGTGGCACCGCCTTTTGTACCGAGTGTGGTGAGGCAATGCCCTCGGACGATGCGAACCCTGACGTCAGTATCTCGCAGGCACAAATGCCGTCGGTCACAATGAATAGCAAGAGCCAAGCACCTGATCGCTGGCAGCAAGTTCCGCACCGGACATTTCTTAGCACGACCTCATTTGTTGACCCGAAGCACGCGAAAGAGGGCGGCGTCAGTATGAAGGATAACGGCCGAAGCATCCGCTTTATGGATTCTGGCGTACAAGGCCCGGACGGCAAGATGAGCCAGTACAATGACTCTATTGCTGGTGTCCGCAACTATTACGAGCCTTCGTTCCGATCGTTTGCTATGTCTACATACTCCATCGCCAACCGTGCTGCCGAGCCTGTGGAAACACCCAACATTCGCAAATACAAGATGTCAGGAAGTGGCATGTTTCACGTGTTTTACTTCGGCACGGCTGCCACTGGTATCATCTGGCTGTATTACTTGACAACAATGTACCCGCAGCAGTATTTCTGCGACCATGCTCGTCCTACGCTTCCCTGCAATGGGTTGCCTACCAGCGAGATTGCGGGCTGCTACAGCTCGACGGTAAACTTTGACGCCAATTCCGGTGATGGATACTGTATCAAGAACGTACCGTTCATGTCATGGCTCATGTACGCAATGATGATTTTCAGTGAGTTTCTCAATTTCTTTTTGGGACTGCTCTTCAACTTTAGTATGTGGCGTCCGATCCGGCGTGGTGCTCGTTACATGAATGACTTCAAGCCGCCAATCCCGAAAGAGCAGTGGCCAACGGTCGACATCTTCTTGTGTCACTACATGGAACCTGTTACGGATTCAATGCAGACGCTGAAGAACTGTCTGGCTATGCAATACCCCCCGGAGTTGCTGCACATTTTTGTTCTGGACGACGGTTACACCAAGTCTGTGTGGGACGCAAACAACCACTTCAAAGTTACGGTAAACACGAAGGTGATTGAGATCTGTGGTGATTTGCGTGGTGATCTTGCACGTCTCATGCACGAGCGTGTTGTCGGGCCTGTGCAGGACGACCAGAGCTTGAAAACGTGGCGCCGCCAGCATAGTTCTGTTCGCGAACTCCGCAAGGAGGGTGGCAAGGGCGTGCAGCGTCGTGATTGTGCTGTAGGCTCGCTTTCGGACGACTACGACTACCGTGACCGCGGCATCCCGCGTGTGACTTTCATTGGTCGTATGAAGCCCGAGACACATCATTCTAAGGCTGGCAACATCAACAACGCCTTGTTTAACGAAGGTGCTGATGGCAAGTATCTGCTGATTTTGGATAACGATATGAAGCCGCATCCGAAGTTTTTGCTTGCCGTGCTGCCGTTCTTCTTCTCCGAAGGGGAGGCTGTTGACGGTGGAGGTCGCCAATACAGCGATGACATTTCCTGGAACCAGGTGGCATATGTACAGACTCCCCAGTACTTTGAAGACACGCCACAACTGACGATCATGGGCGATCCATGTGGTCACAAGAATACCATTTTCTTTGATGCCGTACAGTGTGGTCGTGATGGGTTTGACTCTGCAGCTTTTGCCGGTACAAACGCTGTTTTCCGTCGCCAGGCTTTTGACTCGATTGGTGGCATTCAGTATGGTACACAAACGGAAGATGCTTTCACGGGTAATGTGTTGCACACTTCTGGTTGGGACTCGGTGTACTTTCGCAAGGACTTTGAAGGTGATGCCAAGGACCGCATTCGTCTGTGCGAAGGTGCGGTACCTGAAACAGTGGCTGCTGCCATGGGTCAAAAGAAGCGTTGGGCAAAGGGTGCTGTGCAGATTCTGTTGATGAAGAATGAGAGCGAGGTTGACCCGGACTGGCGTCCGCCGCGCGTTCCTGCCCCAGACCCGAAGCCGTCGCTAACGTTCCCACGTAAGATGTTCTTCTACGATTCGGTTCTGTACCCGTTCGGCTCGATCCCTGCCTTGTGTTACGTGTCGATCGCTGTCTACTACTTATGTACGGGTGACGCTCCGATTTATGCTCGTGGTACCAAGTTTTTGTACTCTTTCTTGCCCGTGACGTTCTGTCGTTGGGTGCTGAATTTGTTGGCTAACCGCGCTGTCGACAACAATGACGTGTGGCGTGCGCAGCAGACGTGGTTTTCTTTTTCGTTCATCACGATGATGGCTATCATTGAAGCCATCCAGGCGCGAATGACGGGCAAGGACAAATCTTGGGCAAACACAGGTGCGGGTCAGAAGACGTCGTGGACAGAGATTCCGAACGTGCTTTTCTTCTTTACGCTGATGTTTAGTCAGTTGGTAGCGCTCATCCGCTTCTTCGAGTATGAGAACGCCACGAATCCATGGAATTACGTGTCTTCCATGTTCTTTGGCTTCTTCGTCATGAGCCAGTTCTACCCCATGGTCAAGATGAGTATCACGGAGTACTGTGGCTGGGACCACACTGCCGCTACATTTACGGCGAACGTGTTCGGGTCGCTGCTCGTGGTCTACATCGTGGTGTTTGTGCAGTTGTGGCAGGTCTACTACG","order":2},{"start":1,"name":"B\_lactucae/1-3370","end":3370,"id":"2087501374","seq":"ATGGGGCTTACTGGCGCTGGCGTAATTGCCTCCGTCATAGGCGTCGTGGGCGGCCTGTCGCTGTCATGTGGCGGCTGGTCGTCCCTCTCGCTCGGCGCGCGCTCGCTCTTTGTCACGACGCAGTTCGTGTCCGCCTTTGCAATG-GGCTTTGTGGTAGCATTTACAGCCATTGTGTCACTTTCCGAAACCAATGAATGGGTCGCGATTGCCGCCGGTGGTGGCGCGGGCTTTGCCATTGCGCTCGTCGTCGGCTTTATGACCATTGTGGGTCCGTACGTGTTGATTTTGCTCACGGGTGGCATCATCGCGATGTATCTGCTGCTTATTGACGCGTACAATGGCATTAACGTGTTCCCCGATGACAATCAATTGGCCCGTCAAGAGTTTGTCATCGCGTTCATGATTATCTTCGAGCTAGTTTGCTGCTCGTCGTCTAAAACGTCGGACATGGAGAATCACCGCTTCAAGTATATTCTTTTCTCGGCTATCACCGGTGGATGGATGACTGCCGACGGTATCTCGCGACTCTTAGACTCGTCTGCCGTGCTGTCGACTGTTGGTTTCCACTCGATTCAAGACGGTGGCAAAGCTGCAATGACCGGCATTGACGCAGGCGGGCAAACGGTCATGTTTATGATCTGGGCCGCAATCTTTATCATTGGAGGGCTAAACCAACTATCAATGCGATGGGGACTTGTGTGCTACAATCGTGTCGGAACGCACGCCCAAATGGGTCCCGTAGACGAGCAATTGCCGGAACTCCCAACTGGTGCCACGCTCCCGGCCCAGACAATAACGGAGCGTGTTCGTTTAGTATGTGAGAATTGCTTTGCTACGGTACCTGCAGGCACAGCCTTTTGTACCGAGTGTGGCGAGGCCATGCCTTCGGAAGACGCGAATCCGGACGTGAGCATCTCGCAAGCGCAGATGCCATCAGTTTCAATGAATAACAAGGGTCAAGTCCCTGACCGATGGCAGCAAGTGCCGCACCGGACGTTTATGAGCACGACTTCGTTTGTTGACCCGAAACTCGCCAAGGAAGGAGGCGTGAGCATGAAGGATAACAGTCGTAGCATTCGCTTTATGGACTCGGGCGTTCAAGGGCCGGATGGCAAGATGGGTCAGTATAACGACTCGATTGCGGGCGTTCGGAATTACTATGAACCGTCGTTTCGATCGTTTGCCATGTCGACGTATTCGATTGCAAATCGCGCGGCGGAACCGGTTGAGACGCCTAATATCCGAAAGTACAAGATGTCGGGAAGCGGCATGTTTCACGTGTTTTACTTTAGCACTGCTGCCACGGGAATCTTTTGGCTGTACTACCTCACGACCATGTACCCGCAGCAGTATTTCTGTGACCATGCCCGTCCGACGCTTCCATGCAATGAGCTTCCTACGAGTGAAACGACGGGCTGTTACAGTTCGACGGTCAACTTTGACAGTGGCTCGGGCGATGGTTATTGCATCAAAGACGTGCCATTTATGTCGTGGATGATGTATGCGATGATGATATTCAGCGAGTTTCTCAATTTTTTCCTGGGACTGCTATTTAACTTTAGTATGTGGCGACCAATTCGTCGTGGTGCGCGGTATATGAATGATTTCAAGCCCCCGATACCGAAGGAACAGTGGCCGACGGTCGACATTTTCTTGTGTCATTACATGGAACCGGTGACGGATTCTATGCAGACGCTCAAGAACTGTCTGGCCATGCAGTATCCTCCTGAGCTGCTGCATATCTTTGTACTTGATGACGGGTACACAAAGTCGGTCTGGGATGCTAATAATCACTTTAAAGTGACGGTGAACACGAAGGTGATTGAGATTGCGGGTGACCTGCGTGGTGATCTTGCGCGTCTCATGCACGAGCGTGTGGTAGGACCGGTACAAGACGACCAGAGTCTCAAGTCGTGGCGTCGTCAGCACAGCTCGGTACGAGAGCTTCGAAAAGAAGGTGGCAAGGGTGTGCAGCGTCGTGACTGTGCCGTAGGATCGCTCTCGGACGATTACGATTACCGCGACCGTGGCATTCCTCGCGTGACGTTCATTGGTCGTATGAAGCCTGAGACGCACCACTCGAAGGCTGGTAACATTAACAATGCCTTGTTCAATGAAGGAGCTGACGGCAAGTATTTGCTGATTCTGGATAATGATATGAAACCGCATCCGAAGTTTCTACTCGCTGTGTTGCCGTTTTTCTTCTCCGAGGGTGAAGCGGTGGACGGCGGGGGTCGCCAGTACAGTGACGACATTTCGTGGAACCAAGTCTCGTACGTCCAGACGCCACAGTATTTTGAGGACACACCGCAATTGACGATCATGGGTGACCCGTGTGGACACAAGAATACCATTTTCTTTGACGCTGTGCAGTGTGGGCGTGACGGGTTTGACTCGGCCGCTTTTGCCGGCACGAACGCGGTCTTTCGTCGCCAGGCCTTCGACTCAATTGGGGGTATCTGCTACGGTACTCAAACAGAAGATGCGTACACTGGCAACGTTCTGCACACTTCTGGGTGGGATTCGGTTTACTTTAGAAAAGATTTCGAAGGCGATGCCAAGGACCGGATCCGGTTGTGTGAAGGTGCTGTGCCCGAGACCGTCGCTGCAGCCATGGGTCAGAAGAAACGTTGGGCCAAGGGTGCCGTGCAGATTCTGCTCATGAAAAGTGAAAGCGAGGTCGACCCGGATTGGCGTCCACCTCGCGTTCCTGCCCCGGACCCGAAGCCGTCGCTTGCGTTTCCACGTAAAATGTTTTTCTACGATTCGGTGCTGTACCCTTTCGGCTCAATTCCGGCGCTGTGTTACGTCGCGATTGCTGTGTACTACTTGTGCACGGGTGATGCGCCTATCTACGCGCGTGGTACTAAGTTTATCTACTCTTTCTTACCCGTGACGTTCTGTCGTTGGGTGCTGAACTTGCTCGCGAATCGTGCTGTTGACAACAATGATGTGTGGCGTGCACAGCAGACGTGGTTTTCCTTCTCTTTTATTACGATGATGGCCATTGTGGAGGCTATTCAGGCTCGTGCGACGGGAAAGGATAAGTCGTGGGCGAACACTGGTGCTGGGCAGAAGACATCATGGACGGAAATCCCTAATGTGCTGTTCTTTTTCACGCTTATGTTTAGCCAAGTCGTGGCGCTCATTCGATTCTTTGAGTACGAGAATGCAACGAATCCGTGGAATTACGTGTCAGCAATGTTTTTTGGCTTTTTCGTCATGAGCCAATTCTATCCCATGGTCAAGATGAGTATCACGGAATATTGTGGATGGGACCACACGGCCGCCACGTTCACGGCCAATGTCTTCGGCTCCTTGCTGGTCGTATACGTTGTCGTGTTCGTACAGCTATGGCAAGTGTATTACG","order":3},{"start":1,"name":"Ph\_sojae/1-3370","end":3370,"id":"1410007435","seq":"ATGGGGCTCACCGGCGCGGGCATCATCGCCTCCGTCGTGGGCATCCTGGGCGGCGTGTCGCTCTCGTGCGGCGGCTGGTCGTCGCTGTCCCTCGGCGCGCGCTCCCTCTTCGTCACGACGCAGTTCCTCTCGGCCTTTGCCAT-GGGGTTCGTCGTTGCCTTCACCGCCATCTCGTCGCTGACGGACACTAACGAGTGGATCGCCGTCGCTGCCGGCGGTGGCGCGGGCTTCGTGGTCGCGCTCATCGTGGGCTTCATGACGATCTTCGGACCGTACATCCTTATCCTGGTCACGGGCGGCATCATCGCGTGCTACCTGCTGCTCATCGACGCCTACGACGGTGTGAACCTCTTCCCGGCCGACAACCAGCTGGCGCGCCAGGAGTTCGTGATCGCCTTCATGATCATCTTCGAGCTCGTGTGCTGCTCGTCCTCCAAGACGTCAGAGCTGGAGAACCACCGCTTCAAGTACATCATCTTCTCGGCCATCACTGGCGGCTGGATGGCGTCGGACGGTGTTTCCCGTCTCATTGACTCGACGGCTGTGCTCTCGGACGTGGGCTTCACCTCGATCCAGGATGGCGGCTCGGCCGCGCTCAAGGGCATCGACGGCAGCGCGCAGACGCTCATGTTCCTGATCTGGGGCGCCGTCTTCGTTGTCGGCGGCCTGAACCAGCTCTCGATGCGCTGGGGACTCATGTGCTACAACCGCGTGGGCGCCCACGCCCAGCTCGGCCCCGTCGAGGAGCAGATGCCGGAGCTGCCCACGGGCGCAACGCTGCCCGCGCAGACCATGAACGAGCGTGTCCGCCTTGTGTGCGAGAACTGTTTTGCTACGGTGCCCAGCGGCACGGCCTTCTGTACCGAGTGTGGTGAGGCCATGCCCTCGGACGAAGCCGACCCGAACGTCAGCATCTCGCAGGCCCAGATGCCGTCGGTCACGATGAACAACAAGTCGCAGGTCCCCGACCGCTGGCAGCAGGTGCCGCACCGCACGTACCTCAGCACGACGTCGTTCGTCGACCCCAAGCACGCCAAGGAGGGCGGCGTGAGCATGAAGGACAACGGCCGCAGCATCCGCTTCATGGACTCGGGTGTGCAGGGCCCGGACGGCAAGATGAGCCAGTACAACGACTCCATCGCTGGCGTGCGCAACTACTACGAGCCTTCGTTCCGTTCGTTCGCCATGTCGACCTACTCGATCGCTAACCGTGCAGCTGAGCCCGTGGACACGCCCAACATCCGCAAGTACAAGATGTCTGGCAGTGGCATGTTTCACGTCTTCTACTTCGGCACGGCTGCTACGGGCGTCTTCTGGCTGTACTACTTGACTACGATGTACCCGCAGCAGTACTTCTGCGACCATGCCCGCCCGACGCTTCCCTGCAGTGCGCTGCCCAGCAGTGAAACTTCGGGCTGCTACAGTTCGACGGTCAACTTCGACGCCGACTCCGGCGAGGGTTACTGCATCCAGGACGTGCCGTTCATGTCGTGGCTCATGTACGCGATGATGATCTTCAGTGAGTTCCTCAACTACTTCCTGGGTCTGCTGTTCAACTTCAGTATGTGGCGCCCTATCCGTCGTGGTGCTCGTTACATGAACGACTTCAAGCCGCCTATCCCGAAGGAGCAGTGGCCGACGGTCGACATCTTCCTGTGTCACTACATGGAACCTGTGACGGACTCCATGGCTACGCTAAAGAACTGTCTTGCTATGCAGTACCCTCCGGAGCTGCTCCATATTTTCGTCCTTGATGATGGTTACGCCAAGTCCGTGTGGGACGCCAACAACCACTTCAAGGTTACGGTGAACACGAAGGTGATAGAGATCTGTGGTGACCTGCGTGGCGATGTCGCCCGCATCATGCACGAGCGTGTGGTCGGCCCCGTGCAGGACGACCAGTCCCTGAAGACGTGGCGTCGCCAGCACAGCTCTGTGCGTGAACTCCGCAAGGAGGGCAGCAAGGGTGTGCAGCGTCGTGACTGTGCCGTCGGCTCGCTGTCAGACGACTACGACTACCGTGACCGCGGTATCCCGCGTGTGACTTTCATTGGTCGCATGAAGCCCGAGACCCACCACTCCAAGGCCGGTAACATCAATAACGCCCTGTTCAACGAAGGCGCCGACGGCAAGTACCTGCTGATTCTGGATAACGATATGAAGCCGCACCCGAAGTTCCTGCTCGCCGTGCTGCCGTTCTTCTTCTCGGAGGGCGAGGCTGTGGACGGTGGAGGCCGCCAGTACAGTGACGACATCTCGTGGAACCAGGTGTCGTACGTGCAGACACCGCAGTACTTCGAGGACACGCCGCAGCTGACGATCATGGGTGACCCGTGTGGACACAAGAACACCATTTTCTTCGACGCTGTGCAGTGTGGCCGTGATGGTTTCGACTCGGCAGCTTTCGCCGGTACGAACGCCGTTTTCCGTCGCCAGGCTTTCGACTCGATCGGTGGCATTCAGTACGGTACCCAGACAGAAGATGCCTACACGGGTAACGTGCTGCACACTTCCGGCTGGGACTCCGTGTACTTCCGTAAGGATTTCGAGGGCGATGCCAAGGACCGCATCCGTCTGTGTGAAGGTGCCGTGCCCGAAACGGTCGCTGCCGCCATGGGTCAGAAGAAGCGTTGGGCCAAGGGTGCCGTGCAGATTCTGCTGATGAAAAATGAAAGCGAGGTGGACCCAGACTGGCGTCCGCCGCGTGTGCCTGCCCCGGACCCGAAGCCGTCGCTCGCGTTCCCGCGTAAGATGTTCTTCTACGACTCGGTGCTGTACCCGTTCGGTTCGATCCCCGCTCTGTGTTACGTGGCTATCGCTATCTACTACCTCTGCACGGGTGACGCTCCTATCTACGCTCGCGGTACCAAGTTCCTGTACTCTTTCTTGCCCGTGACGTTCTGCCGTTGGGTGCTCAACTTGCTGGCCAACCGCGCCGTCGACAACAACGATGTGTGGCGTGCCCAGCAGACGTGGTTCTCCTTCTCCTTCATCACGATGATGGCTATCGTGGAGGCCATCCAGGCGCGTGTGACGGGCAAGGACAAGTCGTGGGCCAACACGGGTGCCGGTCAGAAGACGTCGTGGACGGAAATCCCCAACGTGCTGTTCTTCTTCACGCTGCTCTTCAGTCAGTTGGTGGCGCTGATCCGATTCTTCGAGTACGAGAACGCCACGAACCCGTGGAACTACGTGTCTGCCATGTTCTTCGGCTTCTTCGTCATGAGCCAGTTCTACCCTATGGTCAAGATGAGTATCACGGAGTACTGTGGCTGGGACCACACGGCCGCGACGTTCACGGCCAACGTGTTCGGCTCGCTGCTGGTGGTGTACATTGTGGTATTCGTGCAGCTGTGGCAGGTCTACTACG","order":4},{"start":1,"name":"Ph\_cinnamomi/1-3370","end":3370,"id":"1710197423","seq":"ATGGGGCTCACCGGCGCGGGCATCATCGCCTCCGTCGTGGGCATCCTGGGCGGCGTGTCGCTGTCCTGCGGCGGCTGGTCGTCGCTGTCCCTCGGCGCTCGCTCGCTCTTCGTGACGACGCAGTTCCTCTCGGCCTTCGCCAT-GGGATTCGTGGTCGCCTTCACCGCCATCTCGTCGCTGACAAGCACCAACGAGTGGATCGCCGTGGCGGCCGGCGGCGGCGCGGGCTTCGTGATCGCCCTCATCGTGGGCTTCATGACGGTCTTCGGCCCGTACATCCTGATCCTCGTCACAGGCGGCATCATCGCCTGCTACCTGCTGCTCGTGGACGCGTACGACGGCGTGAAACTCTTCCCGTCAGACAACCAGCTGGCGCGCCAGGAGTTCGTCATCGCCTTCATGATCATCTTCGAGCTCGTGTGCTGCTCGTCGTCCAAGACGTCGGAGCTGGAGAACCACCGCTTCAAGTACATCATCTTCTCGGCCATCACGGGCGGCTGGATGGCCTCGGACGGCGTGTCCCGCCTCATCGACTCGACGGCTGTTCTGTCGGACGTGGCCTACACCTCCATCCAGGACGGCGGCTCGGCCGCGCTCAAGGGCATCGACAGCAGCGCGCAGACGCTCATGTTCCTCATCTGGGGCGCCGTCTTCGTGGTCGGCGGCCTCAACCAGCTCTCCATGCGCTGGGGACTCATGTGCTACAACCGCGTTGGTGCGCACGCCCAGCTCGGCCCCGTCGAGGAGCAGATGCCGGAGCTGCCCACGGGCGCCACGCTGCCTGCCCAGACCATGAACGAGCGCGTGCGCCTCGTGTGCGAGAACTGTTTCGCCACGGTGCCCAGCGGCACGGCCTTCTGTACCGAGTGTGGTGAGGCAATGCCCTCGGACGACGCCGACCCGAACGTCAGCATCTCGCAGGCGCAGATGCCGTCGGTCACGATGAACAACAAGTCTCAAGTGCCCGACCGCTGGCAGCAGGTGCCGCACCGCACGTACCTCAGCACGACGTCGTTTGTCGACCCCAAGCACGCCAAGGAGGGCGGCGTGAGCATGAAGGACAACAGCCGCAGCATCCGCTTCATGGACTCGGGTGTGCAGGGCCCCGACGGCAAGATGAGCCAGTACAACGACTCGATCGCCGGCGTGCGCAACTATTACGAGCCTTCATTCCGCTCGTTCGCCATGTCGACCTACTCGATCGCCAACCGCGCCGCTGAGCCCGTTGACACGCCCAACATCCGCAAGTACAAGATGTCGGGTAGTGGCATGTTCCACGTCTTCTACTTCGGTACGGCTGCCACCGGTATCTTCTGGCTGTACTACTTGACTACGATGTACCCGCAGCAGTACTTCTGCGACCACGCCCGCCCCACGCTTCCCTGCAGTGAGCTGCCCAGCAGCGAGATTTCGGGCTGCTACAGTTCAACGGTCAACTTCGACTCCTCTTCCGGCGAGGGTTACTGCATCAAGAACGTGCCGTTCATGTCGTGGCTCATGTACGCGATGATGATCTTTAGCGAGTTCCTCAACTACTTCCTGGGTCTGCTGTTCAACTTCAGTATGTGGCGTCCGATTCGTCGTGGCGCCCGTTACATGAACGACTTCAAACCGCCTATCCCGAAAGAGCAGTGGCCGACCGTCGACATCTTCCTGTGTCACTACATGGAACCTGTGACGGACTCCATGGCTACGCTGAAGAACTGTCTTGCGATGCAGTACCCTCCGGAGCTGCTGCACATTTTCATCCTTGATGATGGTTACGCCAAGTCTGTGTGGGACGCCAACAACCACTTCAAGGTTACGGTCAACACCAAGGTGATTGAGATTTGTGGTGACCTGCGTGGCGACGTCGCTCGCATCATGCACGAGCGCGTGGTCGGCCCTGTGCAGGACGATCAGTCCCTGAAGACGTGGCGTCGCCAGCACAGCTCTGTGCGTGAGCTCCGCAAAGAGGGAAGCAAGGGCGTGCAGCGTCGCGACTGTGCTGTTGGTTCACTGTCGGACGACTACGACTACCGTGACCGCGGTATCCCGCGTGTGACTTTCATCGGTCGCATGAAGCCCGAAACGCACCACTCCAAGGCTGGTAACATCAACAACGCCCTCTTCAACGAAGGTGCCGATGGCAAGTACTTGCTGATTCTGGATAATGATATGAAGCCGCACCCGAAGTTCTTGCTTGCCGTGCTGCCGTTCTTCTTCTCGGAGGGCGAGGCTGTGGACGGCGGAGGCCGCCAGTACAGTGACGACATTTCCTGGAACCAGGTGTCGTACGTGCAGACTCCTCAGTACTTCGAGGACACGCCCCAGCTGACCATCATGGGAGACCCGTGTGGACACAAGAACACCATTTTCTTCGACGCTGTACAGTGTGGTCGTGATGGTTTCGACTCGGCAGCTTTCGCCGGTACCAACGCCGTTTTCCGTCGCCAGGCTTTCGACTCCATCGGTGGCATTCAGTACGGTACCCAGACGGAAGATGCCTACACGGGTAACGTGCTGCACACTTCTGGCTGGGACTCGGTGTACTTCCGCAAGGATTTCGAGGGTGATGCCAAGGACCGCATTCGTCTGTGCGAAGGTGCCGTGCCCGAAACGGTCGCTGCTGCCATGGGTCAGAAGAAGCGTTGGGCCAAGGGTGCCGTGCAGATTCTGCTCATGAAGAATGAGAGCGAGGTCGACCCGGACTGGCGTCCGCCGCGTGTGCCTGCCCCGGACCCGAAGCCGGCGCTTGCGTTCCCGCGCAAGATGTTCTTCTACGACTCGGTGCTCTACCCGTTCGGTTCCATTCCCGCTCTGTGTTACGTGGCGATCGCTATTTACTACCTGTGTACGGGTGACGCTCCCATCTACGCTCGTGGTACCAAGTTCCTGTACTCTTTCTTGCCCGTGACGTTCTGCCGTTGGGTACTCAACCTGCTGGCCAACCGCGCCGTCGACAACAACGATGTGTGGCGTGCCCAGCAGACCTGGTTCTCCTTCTCCTTCATCACGATGATGGCTATTGTGGAGGCTATCCAGGCGCGTGTGACGGGCAAAGACAAGTCGTGGGCCAACACGGGTGCCGGTCAGAAGACGTCGTGGACAGAGATCCCCAACGTGCTCTTCTTCTTCACGCTGCTCTTTAGTCAACTGGTGGCGCTGATTCGGTTCTTTGAGTACGAGAACGCCACGAACCCGTGGAACTACGTGTCTGCTATGTTCTTCGGCTTCTTCGTGATGAGTCAGTTCTACCCCATGGTCAAGATGAGTATCACGGAGTACTGTGGTTGGGACCACACGGCCGCGACATTTACGGCCAACGTGTTCGGCTCGCTGCTGGTGGTGTACATCGTGGTGTTCGTGCAGCTGTGGCAGGTCTACTACG","order":5},{"start":1,"name":"Ph\_parasitica/1-3370","end":3370,"id":"974518940","seq":"ATGGGGCTCACCGGCGCGGGCATCATCGCCTCCGTCGTGGGCATCCTGGGCGGCGTGTCGCTGTCCTGCGGCGGCTGGTCCTCGCTGTCCCTCGGCGCTCGCTCGCTCTTCGTCACCACGCAGTTCCTCTCGGCCTTCGCCAT-GGGGTTTGTGGTAGCTTTTTCCGCCATCGTCTCCCTCTCGGACACGAATGAGTGGGTTGCCGTGGCCGCCGGTGGCGGCGCAGGCTTCGTGATCGCGCTCATCGTGGGTTTCATGACGATTTTCGGCCCGTACATCCTGATCCTGGTCACAGGCGGCATTATCGCCTGCTATTTGCTGCTAGTGGACGCGTACGATGGCGTCAACGTGTTCCCCGAGGACAACCAGCTGGCTCGCCAGGAGTTCGTGATCGCTTTCATGATCATCTTCGAGCTCGTGTGCTGCTCGTCGTCCAAGACGTCGGAGCTGGAGAACCACCGCTTCAAGTACATCATCTTCTCGGCCATCACTGGTGGCTGGATGGCTGCAGACGGCGTCTCTCGTCTCATCGACTCAAGTGCCGTGCTCTCCACTGTGGCCTACACGTCGATGCAGGACGGTGGCAAGGCTGCACTGGACGGCATCGACTCTAGCGCCCAGACGGTCATGTTCTTGGTCTGGGGCGCTGTCTTCGTGGTCGGCGGACTCAACCAGCTGTCGATGCGTTGGGGTCTCATGTGCTACAACCGCGTGGGCGCTCACGCTCAGCTTGGCCCCGTGGAGGAGCAGATGCCTGAGCTCCCCACTGGCGCCACGCTACCGGCTCAGACTATGACGGAACGTGTCCGTCTGGTGTGTGAGAACTGCTTTGCCACTGTCCCCGCTGGCACTGCCTTCTGTACTGAATGTGGTGAGGCTATGCCCTCGGAAGACGCCAACCCGGACGTGAGCATCTCTCAGGCTCAAATGCCGTCAGTCACGATGAACAACAAGAGCCAAGTGCCCGACCGCTGGCAGCAAGTGCCTCACCGTACGTACCTCAGCACCACGTCGTTTGTGGACCCGAAACACGCCAAGGAGGGCGGAGTGAGCATGAAGGACAACAGTCGCAGCATCCGCTTCATGGACTCTGGCGTCCAGGGGCCGGACGGCAAGATGAGCCAGTACAACGACTCGATCGCTGGCGTTCGCAACTATTACGAGCCGTCGTTCCGCTCATTTGCCATGTCCACGTACTCGATCGCTAACCGTGCGGCTGAACCGGTGGAGACGCCCAACATCCGTAAGTACAAGATGTCGGGCAGCGGCATGTTCCACGTCTTCTACTTCGGTACTGCTGCTACCGGTATCTTCTGGCTGTACTACTTGACTACGATGTACCCGCAACAGTATTTCTGCGACCACGCCCGTCCCACGCTTCCTTGCAGTGAACTCCCGTCCAGTGAGACCTCGGGCTGCTACAGTTCGACTGTCAACTTTGACGCTGACTCGGGTGAAGGCTACTGCATCCAGGACGTGCCGTTCATGTCGTGGCTCATGTACGCGATGATGATCTTCAGCGAGTTCCTCAACTACTTCCTGGGTCTGCTTTTCAACTTCAGTATGTGGCGTCCGATCCGTCGTGGTGCTCGCTACATGAACGACTTCAAGCCGCCTATCCCGAAGGAGCAATGGCCCACAGTCGACATCTTCTTGTGTCACTACATGGAACCGGTGACGGACTCGATGCAGACGCTCAAGAACTGTCTGGCCATGCAGTATCCCCCTGAGTTACTGCATATCTTTATTCTGGATGATGGTTACACCAAGTCGGTGTGGGACGCCAACAACCACTTCAAGGTGACGGTTAACACTAAGGTTATTGAGATCGCTGGTGATCTGCGCGGAGATCTCGCTCGTCTCATGCACGAGCGTGTGGTTGGCCCCGTCCAGGACGACCAGAGCTTGAAGGCGTGGCGTCGTCAACACAGTTCGGTCCGTGAGCTCCGTAAGGAAGGCGGCAAGGGAGTTCAGCGTCGTGACTGTGCTGTGGGCTCGCTGTCTGACGACTACGACTACCGTGACCGTGGTATCCCTCGCGTGACGTTCATTGGACGTATGAAGCCTGAGACGCATCACTCGAAGGCTGGTAACATCAACAACGCGCTGTTCAACGAAGGCGCAGACGGCAAGTACCTGTTGATTCTGGATAACGATATGAAGCCGCATCCCAAGTTCCTGCTGGCTGTGCTGCCGTTCTTCTTCTCGGAGGGCGAGGCTGTTGACGGCGGAGGTCGTCAATACAGTGACGATATCTCGTGGAACCAGGTGTCGTATGTGCAGACCCCGCAGTACTTCGAAGACACCCCGCAATTGACAATCATGGGAGACCCCTGTGGACACAAGAACACCATTTTCTTCGACGCTGTGCAGTGTGGACGTGATGGTTTCGACTCGGCCGCTTTCGCCGGTACGAACGCTGTTTTCCGTCGTCAAGCGTTCGACTCTATTGGTGGAATTCAGTACGGTACACAGACAGAAGATGCGTTCACGGGTAACGTGCTGCACACGTCTGGATGGGACTCGGTGTACTTCCGTAAGGACTTTGAAGGTGATGCCAAGGACCGTATCCGTCTGTGTGAAGGTGCCGTGCCCGAGACAGTCGCTGCTGCTATGGGTCAGAAGAAACGTTGGGCCAAGGGTGCCGTCCAGATTCTGTTGATGAAGAATGAAAGCGAAGTCGACCCGGACTGGCGTCCGCCGCGTGTGCCTGCCCCCGACCCGAAGCCGTCGCTCGCGTTCCCGCGTAAGATGTTCTTCTATGACTCGGTGCTGTACCCGTTCGGTTCGATCCCTGCCTTGTGTTACGTGTCGATCGCCGTCTACTACCTGTGCACGGGTGACGCTCCCATTTACGCTCGTGGTACCAAGTTCCTGTACTCTTTCTTGCCCGTGACGTTCTGCCGTTGGGTGCTCAACCTGCTCGCTAACCGCGCTGTCGACAACAACGACGTGTGGCGTGCACAGCAGACGTGGTTCTCATTCTCGTTCATCACAATGATGGCTATTGTGGAGGCTATTCAGGCGCGTGTGACGGGCAAGGACAAGTCGTGGGCTAACACTGGTGCCGGTCAGAAGACGTCATGGACGGAAATCCCCAACGTGCTGTTCTTCTTCACGCTGCTGTTCAGTCAAGTGGTGGCGCTGGTGCGATTCTTCGAGTATGAGAACGCCACGAACCCGTGGAACTACGTGTCTGCCATGTTCTTTGGCTTCTTCGTCATGAGCCAGTTCTACCCCATGGTGAAGATGAGTATCACGGAATACTGTGGTTGGGACCACACGGCTGCCACCTTCACGGCCAACGTGTTCGGCTCGTTGCTGGTGGTGTACGTGGTGGTGTTCGTGCAGCTGTGGCAAGTGTACTACG","order":6},{"start":1,"name":"Ph\_infestans/1-3370","end":3370,"id":"596717230","seq":"ATGGGGCTCACCGGCGCGGGCATTATCGCCTCCGTCGTGGGTATCCTGGGCGGCGTGTCGCTGTCCTGCGGCGGCTGGTCCTCGCTGTCCCTCGGGGCACGCTCACTCTTCGTTACCACGCAGTTCCTCTCCGCCTTCGCGAT-GGGATTTGTGGTAGCTTTTTCCGCCATCGTCTCCCTCTCGGACACAAATGAGTGGGTCGCCGTGGCCGCCGGAGGTGGTGCAGGCTTCGTGATCGCGCTCATCGTGGGGTTCATGACGATCTTTGGCCCGTACATCCTGATCCTGATCACAGGCGGCCTCATCGCCTGCTATTTGCTGCTCGTGGACGCGTACGACGGCATCAACGTGTTCCCTGCGGACAACCAGCTGGCTCGCCAGGAGTTCGTGATCGCCTTCATGATTATCTTCGAGCTCGTGTGCTGCTCTTCGTCCAAGACGTCGGAGCTGGAGAACCACCGCTTCAAGTACATTGTCTTCTCGGCCATCACGGGTGGCTGGATGGCTGCTGATGGTGTCTCGCGTCTCATCGACTCTGGCGCTGTGCTGTCGACTGTGGCCTACACGTCGATCCAGGACGGCGGCAAGGCGGCAATGGACGGCATCGACGCTAGCGCCCAGACGCTCATGTTCGTGATCTGGGGCGCAGTCTTCGTGGTCGGCGGCCTCAACCAGCTGTCGATGCGCTGGGGTCTCATGTGCTACAACCGCGTGGGTGCACACGCTCAACTGGGCCCCGTGGAAGAGCAGATGCCTGAGCTTCCCACTGGCGCCACTCTTCCAGCTCAGACGATGACGGAACGCGTGCGTCTCGTGTGTGAGAACTGCTTTGCTACTGTTCCTGCTGGCACCGCCTTCTGTACCGAGTGTGGTGAGGCTATGCCCTCGGAAGACGGCAACCCGGACGTGAGCATCTCGCAGGCGCAAATGCCGTCAGTCTCGATGAACAACAAGGGCCAGGTGCCCGATCGCTGGCAGCATGTGCCACACCGCACGTACATGAGTACGACGTCGTTCGTGGACCCCAAACACGCCAAGGAGGGCGGTGTGAGCATGAAGGACAACGGCCGCAGTATCCGCTTCATGGACTCTGGCGTCCAGGGCCCCGACGGCAAGATGAGTCAGTACAACGACTCGATTGCTGGCGTGCGCAACTACTACGAGCCTTCGTTCCGCTCGTTCGCCATGTCCACGTACTCGATCGCTAACCGCGCGGCTGAGCCCGTGGAGACGCCCAACATCCGCAAGTACAAGATGTCGGGTAGTGGCATGTTCCACGTCTTCTACTTCGGCACGGCTGCTACCGGTATCTTCTGGCTCTACTACTTGACGACGATGTACCCGCAGCAGTATTTCTGCGACCACGCGCGTCCTACGCTTCCATGCAGTGGGCTGCCGACAAGTGAGACGACGGGATGCTACAGCTCGACCGTCAACTTTGACGCGGACTCTGGCGATGGATACTGTATCAAGGACGTGCCGTTCATGTCGTGGCTCATGTACGCCATGATGATCTTCAGCGAGTTTCTCAACTACTTCCTGGGTCTGCTATTCAACTTCAGTATGTGGCGCCCCATCCGTCGTGGCGCGCGCTACATGAACGACTTCAAGCCGCCCATCCCGAAGGAGCAGTGGCCCACAGTCGACATCTTCTTGTGTCACTACATGGAACCTGTGACGGACTCTATGCAGACGCTGAAGAACTGTCTGGCCATGCAGTATCCCCCGGAGCTGCTGCACATCTTCATTCTGGATGATGGCTACACCAAGTCGGTGTGGGACGCCAACAACCACTTCAAGGTGACGGTGAACACCAAGGTGATTGAGGTCGCTGGTGACTTGCGTGGCGATCTGGCGCGACTCATGCACGAGCGCGTGGTGGGTCCCGTCCAGGACGACCAGAGCCTGAAGTCGTGGCGTCGTCAGCACAGCTCGGTGCGCGAGCTGCGTAAGGAGGGCGGCAAGGGCGTGCAGCGTCGTGACTGTGCCGTGGGTTCGCTCTCGGATGACTACGACTACCGTGACCGCGGCATCCCTCGTGTGACGTTCATTGGACGTATGAAGCCCGAGACGCACCACTCCAAGGCCGGTAACATCAACAACGCGCTGTTTAACGAAGGCGCAGACGGCAAGTACCTGCTGATTCTGGATAACGATATGAAGCCGCACCCCAAGTTCCTATTGGCGGTGCTGCCCTTCTTCTTCTCGGAGGGCGAGGCGGTGGACGGTGGAGGTCGTCAGTACTCTGACGATATCTCGTGGAACCAGGTCTCGTACGTGCAGACCCCGCAGTACTTCGAAGACACCCCGCAACTGACCATCATGGGCGACCCGTGTGGACACAAGAATACCATTTTCTTCGACGCCGTGCAGTGCGGACGTGACGGCTTCGACTCGGCCGCTTTCGCCGGTACCAACGCCGTTTTCCGTCGTCAAGCCTTCGACTCGATTGGTGGAATCTGCTACGGTACCCAGACAGAAGATGCCTACACGGGTAACGTGCTGCACACGTCTGGCTGGGACTCGGTGTACTTCCGCAAGGACTTTGAAGGAGACGCAAAGGACCGTATCCGTCTGTGCGAAGGAGCTGTGCCCGACACGGTCGCTGCTGCCATGGGACAGAAGAAACGTTGGGCCAAGGGTGCCGTGCAGATTCTGCTGATGAAGAATGAGAGCGAAGTCGACCCGGACTGGCGTCCCCCGCGTGTGCCTGCCCCTGACCCGAAGCCGTCGCTCACGTTCCCGCGTAAGATGTTCTTCTACGACTCGGTGCTGTATCCGTTCGGCTCCATCCCGGCCCTGTGTTACGTGTCGATCGCCGTCTACTACCTGTGCACGGGCGACGCCCCCATTTACGCGCGTGGAACCAAGTTCCTGTACTCTTTCTTGCCCGTGACGTTCTGTCGCTGGGTGCTGAACTTGCTCGCTAACCGCGCAGTGGACAACAACGACGTGTGGCGCGCCCAGCAGACGTGGTTCTCGTTCTCGTTCATTACGATGATGGCTATTGTGGAGGCCATTCAGGCGCGTGTGACAGGCAAGGACAAGTCGTGGGCCAACACGGGTGCCGGTCAGAAGACGTCGTGGACAGAGATTCCCAACGTTCTCTTCTTCTTCACGCTGCTCTTCAGTCAACTGGTGGCGCTGGTGCGTTTCTTTGAGTACGAGAACGCCACGAACCCGTGGAACTACGTGTCTGCCATGTTCTTTGGCTTCTTCGTCATGAGCCAGTTCTACCCCATGGTCAAGATGAGTATCACAGAATACTGTGGCTGGGACCACACGGCTGCCACGTTCACGGCGAACGTGTTCGGCTCGCTGCTGGTGGTGTACATCGTGGTGTTTGTGCAGCTGTGGCAGGTGTACTACG","order":7},{"start":1,"name":"Pl\_halstedii/1-3370","end":3370,"id":"790262798","seq":"ATGGGGCTCACTGGCGCGGGTGTTATCGCCTCCGTCGTGGGTATCCTAGGCGGTCTGTCGCTATCCTGCGGTGGATGGTCGTCGCTGTCACTTGGCGCTCGCTCACTTTTTGTGACTACGCAATTTCTTTCCGCCTTTGCCAT-GGGATTTGTGGTTGCTTTTACTGCCATCGTCTCATTATCGGACACAAATGAATGGGTTGCTTTAGCAGCAGGAGGAGGCGCAGGCTTTGTGATTGCTCTTATTGTTGGATTCATGACGTTCTTTGGACCGTACATACTAATTTTGATCACGGGTGGCGTCATCGCGTGCTACTTGCTGCTTATTGACGCATACAATGGTATCAACGTGTTTCCAGCGGATAATCAATTGGCTCGTCAAGAATTTGTCATTGCTTTTATGATTATTTTCGAGCTTGTATGTAGCTCATCTTCTAAAACAACGGAATTAGAGAATCATCGATTTAAGTATATCATCTTTTCTGCCATAACTGGTGGATGGATGGCTGGTGACGGCGTTTCTCGTCTCATTGACTCGACAGCTGTGCTTTCGACTGTGGCTTACACGTCGTTTCAAGATGGAGGCAAGGCTGCACTACGAGGCATTGACGCGAGCGGGCAATCACTTATGTTTCTTATTTGGGCCGCCGTTGTGGTGATTGGAGGTCTTAACCAGCTGGCCATGCGCTGGGGCCTTGTGTGCTACAATCGTGTGGGTGCCCACGCTCAATTGGGACCTGTTGAAGAGCAGCTGCCAGAGCTTCCTACTGGAGCAACGCTACCAGCTCAAACAATGACCGAACGTGTACGTCTCGTGTGTGAGAATTGCTTTGCCACTGTTCCAGCTGGCACTGCTTTCTGTACAGAATGTGGTGAAGCGATGCCTTCGGAAGACGCAAATCCTGATGTGAGCATTTCACAGGCTCAAATGCCATCTGTGGCTATGAACAATAAAAGTCAAGTGCCAGACCGCTGGCAGCAAGTGCCACATCGTACGTACATGAGTACAACATCATTTGTTGACCCGAAACATGCCAAGGAAGGTGGCGTGAGTATGAAGGACAACAGTCGCAGTATCCGCTTTATGGACTCGGGTGTTCAAGGTCCAGATGGTAAAATGAGTCAGTATAATGACTCAATAGCCGGTGTTCGCAACTACTATGAACCATCATTTCGATCATTTGCCATGTCCACATACTCTATTGCCAATCGCGCTGCGGAACCGGTGGAAACACCTAACATCCGCAAGTACAAAATGTCGGGTAGTGGCATGTTTCACGTCTTTTACTTCGGCACGGCGGCTACCGGTATTTTCTGGTTGTATTATCTCACAACTATGTACCCTCAGCAGTATTTCTGTGACCATGCACGTCCGACGCTTCCATGTAGTGAACTACCGACAAGTGAAATTGTAGGATGCTACAGTTCAACTGTCAACTTTGATGCTGGATCTGGAGAAGGGTATTGTATTAAGGATGTGCCCTTCATGTCGTGGGTCATGTACGCTATGATGATTTTCAGCGAGTTTCTCAATTTTTTCCTGGGTTTGCTGTTTAACTTCAGTATGTGGCGCCCCATTCGTCGTGGAGCTCGTTTCATGAATGACTTTAAGCCGCCTATTCCAAAGGAACAGTGGCCGACTGTTGATATTTTCTTGTGTCACTACATGGAACCAGTGACGGATTCTATGCAGACACTAAAAAATTGTCTGGCCATGCAATATCCTCCTGAACTGCTTCATATCTTTGTTTTGGATGATGGTTACACCAAGTCAGTTTGGGATGCTAATAATCATTTCAAAGTGACAGTCAATACGAAGGTGATTGAGATTGCTGGTGACTTACGTGGCGATCTGGCCCGATTAATGCACGAGCGCGTTGTTGGACCTGTACAGGACGATCAGAGTCTAAAGTCGTGGCGACGTCAGCATAGTTCTGTCCGAGAGCTTCGTAAAGAGGGAGGAAAAGGTGTTCAGCGTCGTGATTGTGCTGTCGGCTCACTATCGGATGATTATGATTACCGTGATCGAGGTATCCCGCGTGTGACCTTCATTGGTCGCATGAAACCTGAGACCCATCACTCCAAGGCCGGTAACATCAACAACGCGCTGTTTAATGAAGGAGCTGACGGAAAGTATCTGCTGATTCTTGATAACGATATGAAGCCGCACCCGAAGTTTCTCCTTGCCGTGCTTCCGTTCTTCTTCTCGGAAGGCGAAGCCGTGGATGGCGGAGGCCGCCAGTACAGTGATGACATTTCGTGGAATCAAGTGTCCTACGTACAGACGCCACAATATTTCGAAGACACGCCGCAGCTGACCATCATGGGTGACCCGTGTGGACACAAGAATACCATTTTCTTCGATGCCGTGCAATGTGGTCGTGATGGTTTTGACTCTGCTGCCTTTGCCGGCACGAACGCTGTTTTCCGCCGACAGGCCTTTGACTCGATTGGTGGCATTTGCTATGGTACGCAGACGGAAGATGCGTATACTGGTAACGTGCTTCACACTTCCGGCTGGGACTCGGTTTACTTTCGAAAGGACTTTGAAGGCGATGCTAAGGATCGTATTCGTCTGTGTGAAGGTGCGGTGCCCGAAACTGTGGCTGCAGCCATGGGCCAGAAGAAACGTTGGGCTAAGGGTGCTGTCCAAATTCTACTTATGAAAAATGAGAGTGAGGTCGACCCGGACTGGCGTCCACCGCGCGTGCCGGCTCCGGACCCAAAGCCGTCGCTAGCTTTTCCGCGTAAGATGTTCTTCTACGATTCAGTGCTTTATCCGTTCGGTTCGATCCCTGCTCTATGCTACGTGGCAATCGCCGTCTACTACCTGTGCACAGGTGATGCACCCATCTACGCACGCGGTACAAAGTTTCTCTATTCTTTCTTGCCTGTGACTTTCTGTCGTTGGGTACTCAACCTGCTGGCCAATCGTGCTGTCGATAACAATGACGTGTGGCGTGCTCAGCAGACGTGGTTCTCCTTCTCATTTATCACGATGATGGCTATCGTTGAGGCTATACAAGCGCGTGTGACGGGCAAAGACAAATCGTGGGCTAATACTGGTGCAGGACAGAAAACGTCCTGGACGGAAATTCCCAACGTTTTGTTTTTCTTCACACTACTTTTCAGTCAGGTGGTGGCCTTGATTCGATTCTTCGAGTATGAAAATGCAACAAATCCCTGGAACTACGTATCTGCCATGTTTTTTGGCTTTTTTGTCATGAGCCAATTCTACCCCATGGTCAAGATGAGTATCACTGAATATTGTGGATGGGACCACACTGCTGCTACCTTTACCGCAAATGTGTTCGGCTCGTTATTGGTCGTTTACGTTGTCGTGTTTGTGCAGCTATGGCAAGTCTACTACG","order":8}],"appSettings":{"globalColorScheme":"Nucleotide","webStartUrl":"https://www.jalview.org/services/launchApp","application":"Jalview","showSeqFeatures":"true","version":"2.11.4.1"},"seqGroups":[],"alignAnnotation":[],"svid":"1.0","seqFeatures":[{"fillColor":"#0000ff","score":0,"sequenceRef":"598887649","featureGroup":"Jalview","description":"","xStart":838,"xEnd":850,"type":"CESA3-E. coli-produced dsRNA"},{"fillColor":"#0000ff","score":0,"sequenceRef":"598887649","featureGroup":"Jalview","description":"","xStart":850,"xEnd":1020,"type":"CESA3-E. coli-produced dsRNA"},{"fillColor":"#0000ff","score":0,"sequenceRef":"598887649","featureGroup":"Jalview","description":"","xStart":1020,"xEnd":1123,"type":"CESA3-E. coli-produced dsRNA"},{"fillColor":"#ff0000","score":0,"sequenceRef":"598887649","featureGroup":"Jalview","description":"","xStart":981,"xEnd":1011,"type":"Hpa-CESA3- Specific SS-dsRNA"},{"fillColor":"#ff0000","score":0,"sequenceRef":"598887649","featureGroup":"Jalview","description":"","xStart":668,"xEnd":676,"type":"Hpa-CESA3-Common-SS-dsRNA"},{"fillColor":"#ff0000","score":0,"sequenceRef":"598887649","featureGroup":"Jalview","description":"","xStart":676,"xEnd":718,"type":"Hpa-CESA3-Common-SS-dsRNA"}]}

  

xml version="1.0"?

Hy\_arabidopsidisPe\_v\_pisiB\_lactucaePh\_sojaePh\_cinnamomiPh\_parasiticaPh\_infestansPl\_halstediiConsensusHy\_arabidopsidisPe\_v\_pisiB\_lactucaePh\_sojaePh\_cinnamomiPh\_parasiticaPh\_infestansPl\_halstediiConsensusHy\_arabidopsidisPe\_v\_pisiB\_lactucaePh\_sojaePh\_cinnamomiPh\_parasiticaPh\_infestansPl\_halstediiConsensusHy\_arabidopsidisPe\_v\_pisiB\_lactucaePh\_sojaePh\_cinnamomiPh\_parasiticaPh\_infestansPl\_halstediiConsensusHy\_arabidopsidisPe\_v\_pisiB\_lactucaePh\_sojaePh\_cinnamomiPh\_parasiticaPh\_infestansPl\_halstediiConsensusHy\_arabidopsidisPe\_v\_pisiB\_lactucaePh\_sojaePh\_cinnamomiPh\_parasiticaPh\_infestansPl\_halstediiConsensusHy\_arabidopsidisPe\_v\_pisiB\_lactucaePh\_sojaePh\_cinnamomiPh\_parasiticaPh\_infestansPl\_halstediiConsensusHy\_arabidopsidisPe\_v\_pisiB\_lactucaePh\_sojaePh\_cinnamomiPh\_parasiticaPh\_infestansPl\_halstediiConsensusHy\_arabidopsidisPe\_v\_pisiB\_lactucaePh\_sojaePh\_cinnamomiPh\_parasiticaPh\_infestansPl\_halstediiConsensusHy\_arabidopsidisPe\_v\_pisiB\_lactucaePh\_sojaePh\_cinnamomiPh\_parasiticaPh\_infestansPl\_halstediiConsensusHy\_arabidopsidisPe\_v\_pisiB\_lactucaePh\_sojaePh\_cinnamomiPh\_parasiticaPh\_infestansPl\_halstediiConsensusHy\_arabidopsidisPe\_v\_pisiB\_lactucaePh\_sojaePh\_cinnamomiPh\_parasiticaPh\_infestansPl\_halstediiConsensusHy\_arabidopsidisPe\_v\_pisiB\_lactucaePh\_sojaePh\_cinnamomiPh\_parasiticaPh\_infestansPl\_halstediiConsensusHy\_arabidopsidisPe\_v\_pisiB\_lactucaePh\_sojaePh\_cinnamomiPh\_parasiticaPh\_infestansPl\_halstediiConsensusHy\_arabidopsidisPe\_v\_pisiB\_lactucaePh\_sojaePh\_cinnamomiPh\_parasiticaPh\_infestansPl\_halstediiConsensusHy\_arabidopsidisPe\_v\_pisiB\_lactucaePh\_sojaePh\_cinnamomiPh\_parasiticaPh\_infestansPl\_halstediiConsensusHy\_arabidopsidisPe\_v\_pisiB\_lactucaePh\_sojaePh\_cinnamomiPh\_parasiticaPh\_infestansPl\_halstediiConsensusHy\_arabidopsidisPe\_v\_pisiB\_lactucaePh\_sojaePh\_cinnamomiPh\_parasiticaPh\_infestansPl\_halstediiConsensusHy\_arabidopsidisPe\_v\_pisiB\_lactucaePh\_sojaePh\_cinnamomiPh\_parasiticaPh\_infestansPl\_halstediiConsensusATGGTGCTGACGGGCGCGGGCGTAATCGCCTCGGTCGTGGGCATCCTCGGCGGCCTGTCGCTTTCATGCGGTGGCTGGTCGTCGCTCTCCCTCGGCGCGCGCTCGCTCTTTGTCACAACGCAGTTCCTCTCGGCCTTTGCCAT-GGGATTTGTCATTGCGTTTACCGCAATCGTCTCGCTCAGCATGGGGCTCACTGGCGCTGGCATAATCGCCTCCGTGGTTGGCATCCTGGGTGGATTGTCGCTATCTTGTGGCGGTTGGTCGTCACTCTCTCTCGGTGCACGCTCTCTCTTTGTCACGACACAGTTTCTGTCTGCCTTTGCCAT-GGGTTTTGTCGTGGCTTTTTCTGCCATTGTTTCTCTGACCATGGGGCTTACTGGCGCTGGCGTAATTGCCTCCGTCATAGGCGTCGTGGGCGGCCTGTCGCTGTCATGTGGCGGCTGGTCGTCCCTCTCGCTCGGCGCGCGCTCGCTCTTTGTCACGACGCAGTTCGTGTCCGCCTTTGCAATG-GGCTTTGTGGTAGCATTTACAGCCATTGTGTCACTTTCCATGGGGCTCACCGGCGCGGGCATCATCGCCTCCGTCGTGGGCATCCTGGGCGGCGTGTCGCTCTCGTGCGGCGGCTGGTCGTCGCTGTCCCTCGGCGCGCGCTCCCTCTTCGTCACGACGCAGTTCCTCTCGGCCTTTGCCAT-GGGGTTCGTCGTTGCCTTCACCGCCATCTCGTCGCTGACGATGGGGCTCACCGGCGCGGGCATCATCGCCTCCGTCGTGGGCATCCTGGGCGGCGTGTCGCTGTCCTGCGGCGGCTGGTCGTCGCTGTCCCTCGGCGCTCGCTCGCTCTTCGTGACGACGCAGTTCCTCTCGGCCTTCGCCAT-GGGATTCGTGGTCGCCTTCACCGCCATCTCGTCGCTGACAATGGGGCTCACCGGCGCGGGCATCATCGCCTCCGTCGTGGGCATCCTGGGCGGCGTGTCGCTGTCCTGCGGCGGCTGGTCCTCGCTGTCCCTCGGCGCTCGCTCGCTCTTCGTCACCACGCAGTTCCTCTCGGCCTTCGCCAT-GGGGTTTGTGGTAGCTTTTTCCGCCATCGTCTCCCTCTCGATGGGGCTCACCGGCGCGGGCATTATCGCCTCCGTCGTGGGTATCCTGGGCGGCGTGTCGCTGTCCTGCGGCGGCTGGTCCTCGCTGTCCCTCGGGGCACGCTCACTCTTCGTTACCACGCAGTTCCTCTCCGCCTTCGCGAT-GGGATTTGTGGTAGCTTTTTCCGCCATCGTCTCCCTCTCGATGGGGCTCACTGGCGCGGGTGTTATCGCCTCCGTCGTGGGTATCCTAGGCGGTCTGTCGCTATCCTGCGGTGGATGGTCGTCGCTGTCACTTGGCGCTCGCTCACTTTTTGTGACTACGCAATTTCTTTCCGCCTTTGCCAT-GGGATTTGTGGTTGCTTTTACTGCCATCGTCTCATTATCGATGGGGCTCACCGGCGCGGGCAT+ATCGCCTCCGTCGTGGGCATCCTGGGCGGCGTGTCGCTGTCCTGCGGCGGCTGGTCGTCGCTGTCCCTCGGCGC+CGCTCGCTCTT+GTCACGACGCAGTTCCTCTCGGCCTTTGCCATGGGGATTTGTGGT+GCTTTTACCGCCATCGTCTCGCT++CGGACACGAACGAATGGGTCGCGCTAGCGGCTGGCGGCGGCGCCGGTTTCTTGATCGCGCTCATTGTCGGCTTCATGACTCTCTGTGGCCCGTACATCCTCATCTTTGCCACGGGCGGCATCGTTGCGTCGTACCTGCTCCTCATTGACGCCTACAACGGCATCAACGTCTTCCCCGCGGACAATCGACACTAACGATTGGGTCGCTGTTGCTGCCGGTGGAGGTGCAGGTTTCGTCGTTGCACTTATCGTAGGCTTTGTGACACTCTTTGGACCGTACGTTTTGATTTTGGTCACAGGTAGCATCATTGCTTCGTACCTTTTGCTCTTTGACGCATTTAATGGTATTAATGTCTTCCCGGAAGACAATCGAAACCAATGAATGGGTCGCGATTGCCGCCGGTGGTGGCGCGGGCTTTGCCATTGCGCTCGTCGTCGGCTTTATGACCATTGTGGGTCCGTACGTGTTGATTTTGCTCACGGGTGGCATCATCGCGATGTATCTGCTGCTTATTGACGCGTACAATGGCATTAACGTGTTCCCCGATGACAATCGACACTAACGAGTGGATCGCCGTCGCTGCCGGCGGTGGCGCGGGCTTCGTGGTCGCGCTCATCGTGGGCTTCATGACGATCTTCGGACCGTACATCCTTATCCTGGTCACGGGCGGCATCATCGCGTGCTACCTGCTGCTCATCGACGCCTACGACGGTGTGAACCTCTTCCCGGCCGACAACCAGCACCAACGAGTGGATCGCCGTGGCGGCCGGCGGCGGCGCGGGCTTCGTGATCGCCCTCATCGTGGGCTTCATGACGGTCTTCGGCCCGTACATCCTGATCCTCGTCACAGGCGGCATCATCGCCTGCTACCTGCTGCTCGTGGACGCGTACGACGGCGTGAAACTCTTCCCGTCAGACAACCGACACGAATGAGTGGGTTGCCGTGGCCGCCGGTGGCGGCGCAGGCTTCGTGATCGCGCTCATCGTGGGTTTCATGACGATTTTCGGCCCGTACATCCTGATCCTGGTCACAGGCGGCATTATCGCCTGCTATTTGCTGCTAGTGGACGCGTACGATGGCGTCAACGTGTTCCCCGAGGACAACCGACACAAATGAGTGGGTCGCCGTGGCCGCCGGAGGTGGTGCAGGCTTCGTGATCGCGCTCATCGTGGGGTTCATGACGATCTTTGGCCCGTACATCCTGATCCTGATCACAGGCGGCCTCATCGCCTGCTATTTGCTGCTCGTGGACGCGTACGACGGCATCAACGTGTTCCCTGCGGACAACCGACACAAATGAATGGGTTGCTTTAGCAGCAGGAGGAGGCGCAGGCTTTGTGATTGCTCTTATTGTTGGATTCATGACGTTCTTTGGACCGTACATACTAATTTTGATCACGGGTGGCGTCATCGCGTGCTACTTGCTGCTTATTGACGCATACAATGGTATCAACGTGTTTCCAGCGGATAATCGACAC+AA+GAGTGGGTCGCCGTGGCCGCCGG+GG+GGCGCAGGCTTCGTGATCGCGCTCATCGTGGGCTTCATGACGATCTTTGGCCCGTACATCCTGATC+TGGTCAC+GGCGGCATCATCGCGTGCTACCTGCTGCTCATTGACGCGTAC+A+GGCATCAACGT+TTCCC+GCGGACAA+CAACTCGCCCGGCAGGAGTTTGTCATTGCCTTTATGATCATCTTTGAGCTCGTGTGCTGCTCGAGCTCCAAGTCGCTCGAGCTCGAGAACCACCGCATCAAGTACGTCGTGTTCTCGGCCATTACGGGCGGCTGGCTCGCTGCCGACGGCGTCTCGCGCCTCATTGACTCGAGCGCTGTGCTCTCAGCTGGCCCGTCAGGAATTTGTCATTTCTTTTATGATTATTTTTGCGCTCGTGTGTTGCTCCAACTCCAAAACAACGGAGCTGGAGAACCACCGCTTTAAATACGTCATCTTTTCGTCCATTACTGGTGGCTGGATGGCGGCAGACGGTATTTCCCGCTTGATCGACTCAGCAGCAGTGTTGTCAATTGGCCCGTCAAGAGTTTGTCATCGCGTTCATGATTATCTTCGAGCTAGTTTGCTGCTCGTCGTCTAAAACGTCGGACATGGAGAATCACCGCTTCAAGTATATTCTTTTCTCGGCTATCACCGGTGGATGGATGACTGCCGACGGTATCTCGCGACTCTTAGACTCGTCTGCCGTGCTGTCAGCTGGCGCGCCAGGAGTTCGTGATCGCCTTCATGATCATCTTCGAGCTCGTGTGCTGCTCGTCCTCCAAGACGTCAGAGCTGGAGAACCACCGCTTCAAGTACATCATCTTCTCGGCCATCACTGGCGGCTGGATGGCGTCGGACGGTGTTTCCCGTCTCATTGACTCGACGGCTGTGCTCTCAGCTGGCGCGCCAGGAGTTCGTCATCGCCTTCATGATCATCTTCGAGCTCGTGTGCTGCTCGTCGTCCAAGACGTCGGAGCTGGAGAACCACCGCTTCAAGTACATCATCTTCTCGGCCATCACGGGCGGCTGGATGGCCTCGGACGGCGTGTCCCGCCTCATCGACTCGACGGCTGTTCTGTCAGCTGGCTCGCCAGGAGTTCGTGATCGCTTTCATGATCATCTTCGAGCTCGTGTGCTGCTCGTCGTCCAAGACGTCGGAGCTGGAGAACCACCGCTTCAAGTACATCATCTTCTCGGCCATCACTGGTGGCTGGATGGCTGCAGACGGCGTCTCTCGTCTCATCGACTCAAGTGCCGTGCTCTCAGCTGGCTCGCCAGGAGTTCGTGATCGCCTTCATGATTATCTTCGAGCTCGTGTGCTGCTCTTCGTCCAAGACGTCGGAGCTGGAGAACCACCGCTTCAAGTACATTGTCTTCTCGGCCATCACGGGTGGCTGGATGGCTGCTGATGGTGTCTCGCGTCTCATCGACTCTGGCGCTGTGCTGTCAATTGGCTCGTCAAGAATTTGTCATTGCTTTTATGATTATTTTCGAGCTTGTATGTAGCTCATCTTCTAAAACAACGGAATTAGAGAATCATCGATTTAAGTATATCATCTTTTCTGCCATAACTGGTGGATGGATGGCTGGTGACGGCGTTTCTCGTCTCATTGACTCGACAGCTGTGCTTTCAGCTGGC+CGCCAGGAGTT+GTCATCGCCTTCATGAT+ATCTTCGAGCTCGTGTGCTGCTCGTCGTCCAAGACGTCGGAGCTGGAGAACCACCGCTTCAAGTACATCATCTTCTCGGCCATCACTGGTGGCTGGATGGCTGC+GACGG+GTCTC+CGTCTCATCGACTCGAC+GCTGTGCTGTCGGACGTGGCGTACCAGTCGATTCAAAATGGCGGCACAGCTGCGCTCAAGGGCATGGACGCTGGCTCGCAGACGCTCATGTTTATCATTTGGGCCGCTGTCGTCGTCATTGGCGGCCTCAATCAGCTCTCGATGCGCTGGGGACTCATGTGCTACAACCGCGTGGGCGCCCACGCACAGCTCGGATCAATCAGCTCTCGATGCGCTGGGGACTCATGTGCTACAACCGCGTGGGCTGATGTGGCCTATTTGTCCATCCAGGATGGCGGCAAAGCTGCTGTAGATGGTATGGACGCAGGCTCTCAGACGCTCATGTTTGTGATCTGGGGCGTAGTTGTCGTAATTGGCGGGCTCAACCAGCTCTCGATGCGCTGGGGACTCATGTGCTACAACCGAGTAGGAGCACACGCTCAGCTCGGCGACTGTTGGTTTCCACTCGATTCAAGACGGTGGCAAAGCTGCAATGACCGGCATTGACGCAGGCGGGCAAACGGTCATGTTTATGATCTGGGCCGCAATCTTTATCATTGGAGGGCTAAACCAACTATCAATGCGATGGGGACTTGTGTGCTACAATCGTGTCGGAACGCACGCCCAAATGGGTGGACGTGGGCTTCACCTCGATCCAGGATGGCGGCTCGGCCGCGCTCAAGGGCATCGACGGCAGCGCGCAGACGCTCATGTTCCTGATCTGGGGCGCCGTCTTCGTTGTCGGCGGCCTGAACCAGCTCTCGATGCGCTGGGGACTCATGTGCTACAACCGCGTGGGCGCCCACGCCCAGCTCGGCGGACGTGGCCTACACCTCCATCCAGGACGGCGGCTCGGCCGCGCTCAAGGGCATCGACAGCAGCGCGCAGACGCTCATGTTCCTCATCTGGGGCGCCGTCTTCGTGGTCGGCGGCCTCAACCAGCTCTCCATGCGCTGGGGACTCATGTGCTACAACCGCGTTGGTGCGCACGCCCAGCTCGGCCACTGTGGCCTACACGTCGATGCAGGACGGTGGCAAGGCTGCACTGGACGGCATCGACTCTAGCGCCCAGACGGTCATGTTCTTGGTCTGGGGCGCTGTCTTCGTGGTCGGCGGACTCAACCAGCTGTCGATGCGTTGGGGTCTCATGTGCTACAACCGCGTGGGCGCTCACGCTCAGCTTGGCGACTGTGGCCTACACGTCGATCCAGGACGGCGGCAAGGCGGCAATGGACGGCATCGACGCTAGCGCCCAGACGCTCATGTTCGTGATCTGGGGCGCAGTCTTCGTGGTCGGCGGCCTCAACCAGCTGTCGATGCGCTGGGGTCTCATGTGCTACAACCGCGTGGGTGCACACGCTCAACTGGGCGACTGTGGCTTACACGTCGTTTCAAGATGGAGGCAAGGCTGCACTACGAGGCATTGACGCGAGCGGGCAATCACTTATGTTTCTTATTTGGGCCGCCGTTGTGGTGATTGGAGGTCTTAACCAGCTGGCCATGCGCTGGGGCCTTGTGTGCTACAATCGTGTGGGTGCCCACGCTCAATTGGGAG++TGTGGCCTACACGTCGATCCAGGA+GGCGGCAAGGCTGCACT+AA+GGCATCGACGCTAGCGCGCAGACGCTCATGTT+CTGATCTGGGGCGC+GTCTTCGTG+T+GGCGGCCTCAACCAGCTCTCGATGCGCTGGGGACTCATGTGCTACAACCGCGTGGG+GCCCACGCTCAGCTCGGCCCCGTGGAGGAGCAGCTACCGGAGCTGCCCACGGGTGCGACGCTCCCGGCGCAGACGCTCACGGAGCGCGTGCGTCTCGTCTGTGAGAACTGCTTTGCGACTGTGCCCAGCGGCACGGCATTCTGCACGGAGTGTGGCGAAGCCATGCCGTCGGAAGACGCGAACCCAGACGTGAGCATCTCACGTGCCCAGCGGCACGGCATTCTGCACGGAGTGTGGCGAAGCCATGCCGTCGGAAGACGCGAACCCAGACGTGAGCATCTCACCCTGTCGAGGAGCAGTTGCCAGAGTTGCCTACCGGTGCGACGCTTCCTGCCCAGGTTATGACTGAACGCGTTCGCCTCGTATGTGAGAATTGTTTTGCGACTGTCCCTAGTGGCACCGCCTTTTGTACCGAGTGTGGTGAGGCAATGCCCTCGGACGATGCGAACCCTGACGTCAGTATCTCGCCCCGTAGACGAGCAATTGCCGGAACTCCCAACTGGTGCCACGCTCCCGGCCCAGACAATAACGGAGCGTGTTCGTTTAGTATGTGAGAATTGCTTTGCTACGGTACCTGCAGGCACAGCCTTTTGTACCGAGTGTGGCGAGGCCATGCCTTCGGAAGACGCGAATCCGGACGTGAGCATCTCGCCCCGTCGAGGAGCAGATGCCGGAGCTGCCCACGGGCGCAACGCTGCCCGCGCAGACCATGAACGAGCGTGTCCGCCTTGTGTGCGAGAACTGTTTTGCTACGGTGCCCAGCGGCACGGCCTTCTGTACCGAGTGTGGTGAGGCCATGCCCTCGGACGAAGCCGACCCGAACGTCAGCATCTCGCCCCGTCGAGGAGCAGATGCCGGAGCTGCCCACGGGCGCCACGCTGCCTGCCCAGACCATGAACGAGCGCGTGCGCCTCGTGTGCGAGAACTGTTTCGCCACGGTGCCCAGCGGCACGGCCTTCTGTACCGAGTGTGGTGAGGCAATGCCCTCGGACGACGCCGACCCGAACGTCAGCATCTCGCCCCGTGGAGGAGCAGATGCCTGAGCTCCCCACTGGCGCCACGCTACCGGCTCAGACTATGACGGAACGTGTCCGTCTGGTGTGTGAGAACTGCTTTGCCACTGTCCCCGCTGGCACTGCCTTCTGTACTGAATGTGGTGAGGCTATGCCCTCGGAAGACGCCAACCCGGACGTGAGCATCTCTCCCCGTGGAAGAGCAGATGCCTGAGCTTCCCACTGGCGCCACTCTTCCAGCTCAGACGATGACGGAACGCGTGCGTCTCGTGTGTGAGAACTGCTTTGCTACTGTTCCTGCTGGCACCGCCTTCTGTACCGAGTGTGGTGAGGCTATGCCCTCGGAAGACGGCAACCCGGACGTGAGCATCTCGCCCTGTTGAAGAGCAGCTGCCAGAGCTTCCTACTGGAGCAACGCTACCAGCTCAAACAATGACCGAACGTGTACGTCTCGTGTGTGAGAATTGCTTTGCCACTGTTCCAGCTGGCACTGCTTTCTGTACAGAATGTGGTGAAGCGATGCCTTCGGAAGACGCAAATCCTGATGTGAGCATTTCACCCCGT+GAGGAGCAGATGCCGGAGCTGCCCACTGGCGCCACGCT+CCGGC+CAGAC+ATGACGGA+CG+GTGCGTCTCGTGTGTGAGAACTGCTTTGC+ACTGTGCCC++TGGCACGGCCTTCTGTACCGAGTGTGGTGAGGCCATGCCCTCGGAAGACGCCAACCCGGACGTGAGCATCTCGCAGGCGCAGATGCCGTCGGTCACGATGAACAGTAAGAGCCAAGCGCCCGAGCGCTGGCAGCAAGTGCCGCATCGCACGTACCTCAGTACGACGTCGTTCGTGGACCCGAAACACGCGAAAGAAGGCGGTGTGAGCATGAAGGACAATGGCCGCAGTATCCGCTTTATGGACGGGGGAGTCCAAGGAGGCGCAGATGCCGTCGGTCACGATGAACAGTAAGAGCCAAGCGCCCGAGCGCTGGCAGCAAGTGCCGCATCGCACGTACCTCAGTACGACGTCGTTCGTGGACCCGAAACACGCGAAAGAAGGCGGTGTGAGCATGAAGGACAATGGCCGCAGTATCCGCTTTATGGACGGGGGAGTCCAAGGAGTGCCGCATCGCACGTACCTCAGTACGACAGGCACAAATGCCGTCGGTCACAATGAATAGCAAGAGCCAAGCACCTGATCGCTGGCAGCAAGTTCCGCACCGGACATTTCTTAGCACGACCTCATTTGTTGACCCGAAGCACGCGAAAGAGGGCGGCGTCAGTATGAAGGATAACGGCCGAAGCATCCGCTTTATGGATTCTGGCGTACAAGGAAGCGCAGATGCCATCAGTTTCAATGAATAACAAGGGTCAAGTCCCTGACCGATGGCAGCAAGTGCCGCACCGGACGTTTATGAGCACGACTTCGTTTGTTGACCCGAAACTCGCCAAGGAAGGAGGCGTGAGCATGAAGGATAACAGTCGTAGCATTCGCTTTATGGACTCGGGCGTTCAAGGAGGCCCAGATGCCGTCGGTCACGATGAACAACAAGTCGCAGGTCCCCGACCGCTGGCAGCAGGTGCCGCACCGCACGTACCTCAGCACGACGTCGTTCGTCGACCCCAAGCACGCCAAGGAGGGCGGCGTGAGCATGAAGGACAACGGCCGCAGCATCCGCTTCATGGACTCGGGTGTGCAGGGAGGCGCAGATGCCGTCGGTCACGATGAACAACAAGTCTCAAGTGCCCGACCGCTGGCAGCAGGTGCCGCACCGCACGTACCTCAGCACGACGTCGTTTGTCGACCCCAAGCACGCCAAGGAGGGCGGCGTGAGCATGAAGGACAACAGCCGCAGCATCCGCTTCATGGACTCGGGTGTGCAGGGAGGCTCAAATGCCGTCAGTCACGATGAACAACAAGAGCCAAGTGCCCGACCGCTGGCAGCAAGTGCCTCACCGTACGTACCTCAGCACCACGTCGTTTGTGGACCCGAAACACGCCAAGGAGGGCGGAGTGAGCATGAAGGACAACAGTCGCAGCATCCGCTTCATGGACTCTGGCGTCCAGGGAGGCGCAAATGCCGTCAGTCTCGATGAACAACAAGGGCCAGGTGCCCGATCGCTGGCAGCATGTGCCACACCGCACGTACATGAGTACGACGTCGTTCGTGGACCCCAAACACGCCAAGGAGGGCGGTGTGAGCATGAAGGACAACGGCCGCAGTATCCGCTTCATGGACTCTGGCGTCCAGGGAGGCTCAAATGCCATCTGTGGCTATGAACAATAAAAGTCAAGTGCCAGACCGCTGGCAGCAAGTGCCACATCGTACGTACATGAGTACAACATCATTTGTTGACCCGAAACATGCCAAGGAAGGTGGCGTGAGTATGAAGGACAACAGTCGCAGTATCCGCTTTATGGACTCGGGTGTTCAAGGAGGCGCA+ATGCCGTCGGTCACGATGAACAACAAGAGCCAAGTGCCCGACCGCTGGCAGCAAGTGCCGCACCGCACGTACCTCAGCACGACGTCGTTTGT+GACCCGAAACACGCCAAGGAGGGCGGCGTGAGCATGAAGGACAAC+GCCGCAGCATCCGCTT+ATGGACTCGGGCGTCCA+GGCCCAGACGGCAAGATGGGGCAGTATAACGACTCGATTGCTGGTGTCCGGAACTACTACGAGCCGTCATTTCGGTCGTTTGCCATGTCGACGTATTCGATTGCGAATCGTGCGGCTGAGCCCGTTGAGACGCCGAATATCCGCAAGTACAAGATGTCGGGCAGTGGCATGTTCCACGTGTTTTACCCCAGACGGCAAGATGGGGCCCGGACGGCAAGATGAGCCAGTACAATGACTCTATTGCTGGTGTCCGCAACTATTACGAGCCTTCGTTCCGATCGTTTGCTATGTCTACATACTCCATCGCCAACCGTGCTGCCGAGCCTGTGGAAACACCCAACATTCGCAAATACAAGATGTCAGGAAGTGGCATGTTTCACGTGTTTTACGCCGGATGGCAAGATGGGTCAGTATAACGACTCGATTGCGGGCGTTCGGAATTACTATGAACCGTCGTTTCGATCGTTTGCCATGTCGACGTATTCGATTGCAAATCGCGCGGCGGAACCGGTTGAGACGCCTAATATCCGAAAGTACAAGATGTCGGGAAGCGGCATGTTTCACGTGTTTTACCCCGGACGGCAAGATGAGCCAGTACAACGACTCCATCGCTGGCGTGCGCAACTACTACGAGCCTTCGTTCCGTTCGTTCGCCATGTCGACCTACTCGATCGCTAACCGTGCAGCTGAGCCCGTGGACACGCCCAACATCCGCAAGTACAAGATGTCTGGCAGTGGCATGTTTCACGTCTTCTACCCCCGACGGCAAGATGAGCCAGTACAACGACTCGATCGCCGGCGTGCGCAACTATTACGAGCCTTCATTCCGCTCGTTCGCCATGTCGACCTACTCGATCGCCAACCGCGCCGCTGAGCCCGTTGACACGCCCAACATCCGCAAGTACAAGATGTCGGGTAGTGGCATGTTCCACGTCTTCTACGCCGGACGGCAAGATGAGCCAGTACAACGACTCGATCGCTGGCGTTCGCAACTATTACGAGCCGTCGTTCCGCTCATTTGCCATGTCCACGTACTCGATCGCTAACCGTGCGGCTGAACCGGTGGAGACGCCCAACATCCGTAAGTACAAGATGTCGGGCAGCGGCATGTTCCACGTCTTCTACCCCCGACGGCAAGATGAGTCAGTACAACGACTCGATTGCTGGCGTGCGCAACTACTACGAGCCTTCGTTCCGCTCGTTCGCCATGTCCACGTACTCGATCGCTAACCGCGCGGCTGAGCCCGTGGAGACGCCCAACATCCGCAAGTACAAGATGTCGGGTAGTGGCATGTTCCACGTCTTCTACTCCAGATGGTAAAATGAGTCAGTATAATGACTCAATAGCCGGTGTTCGCAACTACTATGAACCATCATTTCGATCATTTGCCATGTCCACATACTCTATTGCCAATCGCGCTGCGGAACCGGTGGAAACACCTAACATCCGCAAGTACAAAATGTCGGGTAGTGGCATGTTTCACGTCTTTTACCCCGGACGGCAAGATGAGCCAGTACAACGACTCGATTGCTGGCGT+CGCAACTACTACGAGCCTTCGTTCCG+TCGTTTGCCATGTCGACGTACTCGATCGC+AACCG+GCGGCTGAGCCCGTGGAGACGCCCAACATCCGCAAGTACAAGATGTCGGG+AGTGGCATGTT+CACGTCTT+TACTTTGGCACGGCGGCTACGGGTATCATTTGGCTGTATTACTTGACGACGATGTACCCGCAGCAGTACTTTTGTGACCATGCGCGACCAACGTTACCGTGCGACATGTTGCCGGCCAGTGAAACGGAAGGCTGTTTCAGTTCGACGGTCAACTTTGACGCGGACTCGGGTGATGGATACTGTATCCTTCGGCACGGCTGCCACTGGTATCATCTGGCTGTATTACTTGACAACAATGTACCCGCAGCAGTATTTCTGCGACCATGCTCGTCCTACGCTTCCCTGCAATGGGTTGCCTACCAGCGAGATTGCGGGCTGCTACAGCTCGACGGTAAACTTTGACGCCAATTCCGGTGATGGATACTGTATCATTTAGCACTGCTGCCACGGGAATCTTTTGGCTGTACTACCTCACGACCATGTACCCGCAGCAGTATTTCTGTGACCATGCCCGTCCGACGCTTCCATGCAATGAGCTTCCTACGAGTGAAACGACGGGCTGTTACAGTTCGACGGTCAACTTTGACAGTGGCTCGGGCGATGGTTATTGCATCATTCGGCACGGCTGCTACGGGCGTCTTCTGGCTGTACTACTTGACTACGATGTACCCGCAGCAGTACTTCTGCGACCATGCCCGCCCGACGCTTCCCTGCAGTGCGCTGCCCAGCAGTGAAACTTCGGGCTGCTACAGTTCGACGGTCAACTTCGACGCCGACTCCGGCGAGGGTTACTGCATCCTTCGGTACGGCTGCCACCGGTATCTTCTGGCTGTACTACTTGACTACGATGTACCCGCAGCAGTACTTCTGCGACCACGCCCGCCCCACGCTTCCCTGCAGTGAGCTGCCCAGCAGCGAGATTTCGGGCTGCTACAGTTCAACGGTCAACTTCGACTCCTCTTCCGGCGAGGGTTACTGCATCATTCGGTACTGCTGCTACCGGTATCTTCTGGCTGTACTACTTGACTACGATGTACCCGCAACAGTATTTCTGCGACCACGCCCGTCCCACGCTTCCTTGCAGTGAACTCCCGTCCAGTGAGACCTCGGGCTGCTACAGTTCGACTGTCAACTTTGACGCTGACTCGGGTGAAGGCTACTGCATCCTTCGGCACGGCTGCTACCGGTATCTTCTGGCTCTACTACTTGACGACGATGTACCCGCAGCAGTATTTCTGCGACCACGCGCGTCCTACGCTTCCATGCAGTGGGCTGCCGACAAGTGAGACGACGGGATGCTACAGCTCGACCGTCAACTTTGACGCGGACTCTGGCGATGGATACTGTATCATTCGGCACGGCGGCTACCGGTATTTTCTGGTTGTATTATCTCACAACTATGTACCCTCAGCAGTATTTCTGTGACCATGCACGTCCGACGCTTCCATGTAGTGAACTACCGACAAGTGAAATTGTAGGATGCTACAGTTCAACTGTCAACTTTGATGCTGGATCTGGAGAAGGGTATTGTATTATTCGGCACGGCTGCTACCGGTATCTTCTGGCTGTACTACTTGAC+ACGATGTACCCGCAGCAGTATTTCTGCGACCATGCCCGTCCGACGCTTCC+TGCAGTGAGCTGCCGACCAGTGA+ACT+CGGGCTGCTACAGTTCGACGGTCAACTTTGACGC+GACTC+GGCGATGG+TACTG+ATCAGTAACGTGCCGTTCATGTCGTGGATCATGTACGCTATGATGATTTTCAGTGAGTTTCTCAATTTCTTTCTGGGACTGCTGTTCAACTTCAGTATGTGGCGTCCGATTCGTCGTGGAGCGCGGTACATGAACGATTTCAAGCCGCCTATTCCAAAGGAACAGTGGCCGACGGTCGATATCTTTTTAGAACGTACCGTTCATGTCATGGCTCATGTACGCAATGATGATTTTCAGTGAGTTTCTCAATTTCTTTTTGGGACTGCTCTTCAACTTTAGTATGTGGCGTCCGATCCGGCGTGGTGCTCGTTACATGAATGACTTCAAGCCGCCAATCCCGAAAGAGCAGTGGCCAACGGTCGACATCTTCTTAAGACGTGCCATTTATGTCGTGGATGATGTATGCGATGATGATATTCAGCGAGTTTCTCAATTTTTTCCTGGGACTGCTATTTAACTTTAGTATGTGGCGACCAATTCGTCGTGGTGCGCGGTATATGAATGATTTCAAGCCCCCGATACCGAAGGAACAGTGGCCGACGGTCGACATTTTCTTAGGACGTGCCGTTCATGTCGTGGCTCATGTACGCGATGATGATCTTCAGTGAGTTCCTCAACTACTTCCTGGGTCTGCTGTTCAACTTCAGTATGTGGCGCCCTATCCGTCGTGGTGCTCGTTACATGAACGACTTCAAGCCGCCTATCCCGAAGGAGCAGTGGCCGACGGTCGACATCTTCCTAGAACGTGCCGTTCATGTCGTGGCTCATGTACGCGATGATGATCTTTAGCGAGTTCCTCAACTACTTCCTGGGTCTGCTGTTCAACTTCAGTATGTGGCGTCCGATTCGTCGTGGCGCCCGTTACATGAACGACTTCAAACCGCCTATCCCGAAAGAGCAGTGGCCGACCGTCGACATCTTCCTAGGACGTGCCGTTCATGTCGTGGCTCATGTACGCGATGATGATCTTCAGCGAGTTCCTCAACTACTTCCTGGGTCTGCTTTTCAACTTCAGTATGTGGCGTCCGATCCGTCGTGGTGCTCGCTACATGAACGACTTCAAGCCGCCTATCCCGAAGGAGCAATGGCCCACAGTCGACATCTTCTTAGGACGTGCCGTTCATGTCGTGGCTCATGTACGCCATGATGATCTTCAGCGAGTTTCTCAACTACTTCCTGGGTCTGCTATTCAACTTCAGTATGTGGCGCCCCATCCGTCGTGGCGCGCGCTACATGAACGACTTCAAGCCGCCCATCCCGAAGGAGCAGTGGCCCACAGTCGACATCTTCTTAGGATGTGCCCTTCATGTCGTGGGTCATGTACGCTATGATGATTTTCAGCGAGTTTCTCAATTTTTTCCTGGGTTTGCTGTTTAACTTCAGTATGTGGCGCCCCATTCGTCGTGGAGCTCGTTTCATGAATGACTTTAAGCCGCCTATTCCAAAGGAACAGTGGCCGACTGTTGATATTTTCTTAGGACGTGCCGTTCATGTCGTGGCTCATGTACGCGATGATGATCTTCAGCGAGTTTCTCAA+T+CTTCCTGGGTCTGCTGTTCAACTTCAGTATGTGGCGTCCGAT+CGTCGTGGTGCTCGTTACATGAACGACTTCAAGCCGCCTATCCCGAAGGAGCAGTGGCCGACGGTCGACATCTTCTTGTGTCACTACATGGAACCGGTCACGGACTCGATGCAGACACTGAAGAACTGTCTTTCACTCCAGTACCCGCCCGAGTTACTGCACATCTTTGTGCTGGATGATGGTTACACCAAGTCGGTATGGGACGCGAACAACCACTTTAAGGTGACGGTGAACACTAAGGTGATTGAGATTTGTGGTGATGTGTCACTACATGGAACCTGTTACGGATTCAATGCAGACGCTGAAGAACTGTCTGGCTATGCAATACCCCCCGGAGTTGCTGCACATTTTTGTTCTGGACGACGGTTACACCAAGTCTGTGTGGGACGCAAACAACCACTTCAAAGTTACGGTAAACACGAAGGTGATTGAGATCTGTGGTGATGTGTCATTACATGGAACCGGTGACGGATTCTATGCAGACGCTCAAGAACTGTCTGGCCATGCAGTATCCTCCTGAGCTGCTGCATATCTTTGTACTTGATGACGGGTACACAAAGTCGGTCTGGGATGCTAATAATCACTTTAAAGTGACGGTGAACACGAAGGTGATTGAGATTGCGGGTGACGTGTCACTACATGGAACCTGTGACGGACTCCATGGCTACGCTAAAGAACTGTCTTGCTATGCAGTACCCTCCGGAGCTGCTCCATATTTTCGTCCTTGATGATGGTTACGCCAAGTCCGTGTGGGACGCCAACAACCACTTCAAGGTTACGGTGAACACGAAGGTGATAGAGATCTGTGGTGACGTGTCACTACATGGAACCTGTGACGGACTCCATGGCTACGCTGAAGAACTGTCTTGCGATGCAGTACCCTCCGGAGCTGCTGCACATTTTCATCCTTGATGATGGTTACGCCAAGTCTGTGTGGGACGCCAACAACCACTTCAAGGTTACGGTCAACACCAAGGTGATTGAGATTTGTGGTGACGTGTCACTACATGGAACCGGTGACGGACTCGATGCAGACGCTCAAGAACTGTCTGGCCATGCAGTATCCCCCTGAGTTACTGCATATCTTTATTCTGGATGATGGTTACACCAAGTCGGTGTGGGACGCCAACAACCACTTCAAGGTGACGGTTAACACTAAGGTTATTGAGATCGCTGGTGATGTGTCACTACATGGAACCTGTGACGGACTCTATGCAGACGCTGAAGAACTGTCTGGCCATGCAGTATCCCCCGGAGCTGCTGCACATCTTCATTCTGGATGATGGCTACACCAAGTCGGTGTGGGACGCCAACAACCACTTCAAGGTGACGGTGAACACCAAGGTGATTGAGGTCGCTGGTGACGTGTCACTACATGGAACCAGTGACGGATTCTATGCAGACACTAAAAAATTGTCTGGCCATGCAATATCCTCCTGAACTGCTTCATATCTTTGTTTTGGATGATGGTTACACCAAGTCAGTTTGGGATGCTAATAATCATTTCAAAGTGACAGTCAATACGAAGGTGATTGAGATTGCTGGTGACGTGTCACTACATGGAACCTGTGACGGACTCTATGCAGACGCTGAAGAACTGTCTGGCCATGCAGTA+CCTCCGGAGCTGCTGCA+ATCTTTGTTCTGGATGATGGTTACACCAAGTCGGTGTGGGACGCCAACAACCACTTCAAGGTGACGGTGAACACGAAGGTGATTGAGAT+++TGGTGACTTGCGTGGTGACCTCGCTCGGCTCATGCACGAGCGTGTGGTTGGCCCTGTGCAGGACGACCAAAGCTTGAAGACGTGGCGTCGCCAGCACAGCTCTGTCCGTGAACTCCGCAAGGAGGGTGGCAAGGGCGTGCAGCGTCGTGACTGTGCTGTTGGCTCGCTTTCGGACGACTACGACTACCGTGTTGCGTGGTGATCTTGCACGTCTCATGCACGAGCGTGTTGTCGGGCCTGTGCAGGACGACCAGAGCTTGAAAACGTGGCGCCGCCAGCATAGTTCTGTTCGCGAACTCCGCAAGGAGGGTGGCAAGGGCGTGCAGCGTCGTGATTGTGCTGTAGGCTCGCTTTCGGACGACTACGACTACCGTGCTGCGTGGTGATCTTGCGCGTCTCATGCACGAGCGTGTGGTAGGACCGGTACAAGACGACCAGAGTCTCAAGTCGTGGCGTCGTCAGCACAGCTCGGTACGAGAGCTTCGAAAAGAAGGTGGCAAGGGTGTGCAGCGTCGTGACTGTGCCGTAGGATCGCTCTCGGACGATTACGATTACCGCGCTGCGTGGCGATGTCGCCCGCATCATGCACGAGCGTGTGGTCGGCCCCGTGCAGGACGACCAGTCCCTGAAGACGTGGCGTCGCCAGCACAGCTCTGTGCGTGAACTCCGCAAGGAGGGCAGCAAGGGTGTGCAGCGTCGTGACTGTGCCGTCGGCTCGCTGTCAGACGACTACGACTACCGTGCTGCGTGGCGACGTCGCTCGCATCATGCACGAGCGCGTGGTCGGCCCTGTGCAGGACGATCAGTCCCTGAAGACGTGGCGTCGCCAGCACAGCTCTGTGCGTGAGCTCCGCAAAGAGGGAAGCAAGGGCGTGCAGCGTCGCGACTGTGCTGTTGGTTCACTGTCGGACGACTACGACTACCGTGCTGCGCGGAGATCTCGCTCGTCTCATGCACGAGCGTGTGGTTGGCCCCGTCCAGGACGACCAGAGCTTGAAGGCGTGGCGTCGTCAACACAGTTCGGTCCGTGAGCTCCGTAAGGAAGGCGGCAAGGGAGTTCAGCGTCGTGACTGTGCTGTGGGCTCGCTGTCTGACGACTACGACTACCGTGTTGCGTGGCGATCTGGCGCGACTCATGCACGAGCGCGTGGTGGGTCCCGTCCAGGACGACCAGAGCCTGAAGTCGTGGCGTCGTCAGCACAGCTCGGTGCGCGAGCTGCGTAAGGAGGGCGGCAAGGGCGTGCAGCGTCGTGACTGTGCCGTGGGTTCGCTCTCGGATGACTACGACTACCGTGTTACGTGGCGATCTGGCCCGATTAATGCACGAGCGCGTTGTTGGACCTGTACAGGACGATCAGAGTCTAAAGTCGTGGCGACGTCAGCATAGTTCTGTCCGAGAGCTTCGTAAAGAGGGAGGAAAAGGTGTTCAGCGTCGTGATTGTGCTGTCGGCTCACTATCGGATGATTATGATTACCGTG+TGCGTGGCGATCTCGCTCGTCTCATGCACGAGCGTGTGGT+GGCCCTGTGCAGGACGACCAGAGCCTGAAGACGTGGCGTCG+CAGCACAGCTCTGT+CGTGAGCTCCGCAAGGAGGG+GGCAAGGGCGTGCAGCGTCGTGACTGTGCTGT+GGCTCGCTGTCGGACGACTACGACTACCGTGACCGTGGTATCCCTCGTGTAACGTTCATCGGTCGTATGAAGCCCGAGACGCACCACTCCAAGGCTGGTAACATCAACAATTGCTTGTTCAACGAAGGTGCCGACGGCAAGTATTTGCTGATTCTGGATAACGACATGAAGCCGCATCCGAAGTTTTTGCTTGCGGTGCTGCCGTTTTTCTTCTCACCGCGGCATCCCGCGTGTGACTTTCATTGGTCGTATGAAGCCCGAGACACATCATTCTAAGGCTGGCAACATCAACAACGCCTTGTTTAACGAAGGTGCTGATGGCAAGTATCTGCTGATTTTGGATAACGATATGAAGCCGCATCCGAAGTTTTTGCTTGCCGTGCTGCCGTTCTTCTTCTCACCGTGGCATTCCTCGCGTGACGTTCATTGGTCGTATGAAGCCTGAGACGCACCACTCGAAGGCTGGTAACATTAACAATGCCTTGTTCAATGAAGGAGCTGACGGCAAGTATTTGCTGATTCTGGATAATGATATGAAACCGCATCCGAAGTTTCTACTCGCTGTGTTGCCGTTTTTCTTCTCACCGCGGTATCCCGCGTGTGACTTTCATTGGTCGCATGAAGCCCGAGACCCACCACTCCAAGGCCGGTAACATCAATAACGCCCTGTTCAACGAAGGCGCCGACGGCAAGTACCTGCTGATTCTGGATAACGATATGAAGCCGCACCCGAAGTTCCTGCTCGCCGTGCTGCCGTTCTTCTTCTCACCGCGGTATCCCGCGTGTGACTTTCATCGGTCGCATGAAGCCCGAAACGCACCACTCCAAGGCTGGTAACATCAACAACGCCCTCTTCAACGAAGGTGCCGATGGCAAGTACTTGCTGATTCTGGATAATGATATGAAGCCGCACCCGAAGTTCTTGCTTGCCGTGCTGCCGTTCTTCTTCTCACCGTGGTATCCCTCGCGTGACGTTCATTGGACGTATGAAGCCTGAGACGCATCACTCGAAGGCTGGTAACATCAACAACGCGCTGTTCAACGAAGGCGCAGACGGCAAGTACCTGTTGATTCTGGATAACGATATGAAGCCGCATCCCAAGTTCCTGCTGGCTGTGCTGCCGTTCTTCTTCTCACCGCGGCATCCCTCGTGTGACGTTCATTGGACGTATGAAGCCCGAGACGCACCACTCCAAGGCCGGTAACATCAACAACGCGCTGTTTAACGAAGGCGCAGACGGCAAGTACCTGCTGATTCTGGATAACGATATGAAGCCGCACCCCAAGTTCCTATTGGCGGTGCTGCCCTTCTTCTTCTCATCGAGGTATCCCGCGTGTGACCTTCATTGGTCGCATGAAACCTGAGACCCATCACTCCAAGGCCGGTAACATCAACAACGCGCTGTTTAATGAAGGAGCTGACGGAAAGTATCTGCTGATTCTTGATAACGATATGAAGCCGCACCCGAAGTTTCTCCTTGCCGTGCTTCCGTTCTTCTTCTCACCGCGGTATCCC+CGTGTGACGTTCATTGGTCGTATGAAGCCCGAGACGCACCACTCCAAGGCTGGTAACATCAACAACGCCCTGTTCAACGAAGG+GC+GACGGCAAGTA+CTGCTGATTCTGGATAACGATATGAAGCCGCA+CCGAAGTT+CTGCTTGCCGTGCTGCCGTTCTTCTTCTCGGAAGGCGAGGCTGTTGACGGTGGAGGACGTCAGTACAGTGACGACATTTCGTGGAACCAGGTGTCATACGTGCAGACTCCTCAGTACTTCGAGGACACGCCGCAGTTGACTATCATGGGTGATCCGTGTGGCCACAAGAATACCATTTTCTTCGACGCTGTGCAGTGTGGTCGTGATGGTTTCCGAAGGGGAGGCTGTTGACGGTGGAGGTCGCCAATACAGCGATGACATTTCCTGGAACCAGGTGGCATATGTACAGACTCCCCAGTACTTTGAAGACACGCCACAACTGACGATCATGGGCGATCCATGTGGTCACAAGAATACCATTTTCTTTGATGCCGTACAGTGTGGTCGTGATGGGTTTCGAGGGTGAAGCGGTGGACGGCGGGGGTCGCCAGTACAGTGACGACATTTCGTGGAACCAAGTCTCGTACGTCCAGACGCCACAGTATTTTGAGGACACACCGCAATTGACGATCATGGGTGACCCGTGTGGACACAAGAATACCATTTTCTTTGACGCTGTGCAGTGTGGGCGTGACGGGTTTGGAGGGCGAGGCTGTGGACGGTGGAGGCCGCCAGTACAGTGACGACATCTCGTGGAACCAGGTGTCGTACGTGCAGACACCGCAGTACTTCGAGGACACGCCGCAGCTGACGATCATGGGTGACCCGTGTGGACACAAGAACACCATTTTCTTCGACGCTGTGCAGTGTGGCCGTGATGGTTTCGGAGGGCGAGGCTGTGGACGGCGGAGGCCGCCAGTACAGTGACGACATTTCCTGGAACCAGGTGTCGTACGTGCAGACTCCTCAGTACTTCGAGGACACGCCCCAGCTGACCATCATGGGAGACCCGTGTGGACACAAGAACACCATTTTCTTCGACGCTGTACAGTGTGGTCGTGATGGTTTCGGAGGGCGAGGCTGTTGACGGCGGAGGTCGTCAATACAGTGACGATATCTCGTGGAACCAGGTGTCGTATGTGCAGACCCCGCAGTACTTCGAAGACACCCCGCAATTGACAATCATGGGAGACCCCTGTGGACACAAGAACACCATTTTCTTCGACGCTGTGCAGTGTGGACGTGATGGTTTCGGAGGGCGAGGCGGTGGACGGTGGAGGTCGTCAGTACTCTGACGATATCTCGTGGAACCAGGTCTCGTACGTGCAGACCCCGCAGTACTTCGAAGACACCCCGCAACTGACCATCATGGGCGACCCGTGTGGACACAAGAATACCATTTTCTTCGACGCCGTGCAGTGCGGACGTGACGGCTTCGGAAGGCGAAGCCGTGGATGGCGGAGGCCGCCAGTACAGTGATGACATTTCGTGGAATCAAGTGTCCTACGTACAGACGCCACAATATTTCGAAGACACGCCGCAGCTGACCATCATGGGTGACCCGTGTGGACACAAGAATACCATTTTCTTCGATGCCGTGCAATGTGGTCGTGATGGTTTTGGAGGGCGAGGCTGTGGACGG+GGAGGTCGCCAGTACAGTGACGACATTTCGTGGAACCAGGTGTCGTACGTGCAGACTCCGCAGTACTTCGA+GACACGCCGCA+CTGAC+ATCATGGGTGACCCGTGTGGACACAAGAATACCATTTTCTTCGACGCTGTGCAGTGTGGTCGTGATGGTTTCGACTCTGCAGCTTTTGCCGGTACGAATGCTGTTTTCCGTCGTCAAGCTTTCGATTCGATTGGTGGTATACAGTACGGCACGCAGACAGAAGATGCCTTTACGGGTAACGTGCTGCACACTTCTGGTTGGGACTCGGTGTATTTCCGCAAGGATTTCGAGGGCGATGCCAAGGACCGTATCCGCCGACTCTGCAGCTTTTGCCGGTACAAACGCTGTTTTCCGTCGCCAGGCTTTTGACTCGATTGGTGGCATTCAGTATGGTACACAAACGGAAGATGCTTTCACGGGTAATGTGTTGCACACTTCTGGTTGGGACTCGGTGTACTTTCGCAAGGACTTTGAAGGTGATGCCAAGGACCGCATTCGTCGACTCGGCCGCTTTTGCCGGCACGAACGCGGTCTTTCGTCGCCAGGCCTTCGACTCAATTGGGGGTATCTGCTACGGTACTCAAACAGAAGATGCGTACACTGGCAACGTTCTGCACACTTCTGGGTGGGATTCGGTTTACTTTAGAAAAGATTTCGAAGGCGATGCCAAGGACCGGATCCGGTGACTCGGCAGCTTTCGCCGGTACGAACGCCGTTTTCCGTCGCCAGGCTTTCGACTCGATCGGTGGCATTCAGTACGGTACCCAGACAGAAGATGCCTACACGGGTAACGTGCTGCACACTTCCGGCTGGGACTCCGTGTACTTCCGTAAGGATTTCGAGGGCGATGCCAAGGACCGCATCCGTCGACTCGGCAGCTTTCGCCGGTACCAACGCCGTTTTCCGTCGCCAGGCTTTCGACTCCATCGGTGGCATTCAGTACGGTACCCAGACGGAAGATGCCTACACGGGTAACGTGCTGCACACTTCTGGCTGGGACTCGGTGTACTTCCGCAAGGATTTCGAGGGTGATGCCAAGGACCGCATTCGTCGACTCGGCCGCTTTCGCCGGTACGAACGCTGTTTTCCGTCGTCAAGCGTTCGACTCTATTGGTGGAATTCAGTACGGTACACAGACAGAAGATGCGTTCACGGGTAACGTGCTGCACACGTCTGGATGGGACTCGGTGTACTTCCGTAAGGACTTTGAAGGTGATGCCAAGGACCGTATCCGTCGACTCGGCCGCTTTCGCCGGTACCAACGCCGTTTTCCGTCGTCAAGCCTTCGACTCGATTGGTGGAATCTGCTACGGTACCCAGACAGAAGATGCCTACACGGGTAACGTGCTGCACACGTCTGGCTGGGACTCGGTGTACTTCCGCAAGGACTTTGAAGGAGACGCAAAGGACCGTATCCGTCGACTCTGCTGCCTTTGCCGGCACGAACGCTGTTTTCCGCCGACAGGCCTTTGACTCGATTGGTGGCATTTGCTATGGTACGCAGACGGAAGATGCGTATACTGGTAACGTGCTTCACACTTCCGGCTGGGACTCGGTTTACTTTCGAAAGGACTTTGAAGGCGATGCTAAGGATCGTATTCGTCGACTCGGCAGCTTT+GCCGGTACGAACGCTGTTTTCCGTCGCCAGGCTTTCGACTCGATTGGTGGCATTCAGTACGGTACCCAGACAGAAGATGCCTACACGGGTAACGTGCTGCACACTTCTGGCTGGGACTCGGTGTACTTCCGCAAGGA+TT+GAAGGCGATGCCAAGGACCGTATCCGTCTATGCGAAGGTGCCGTTCCCGAGACGGTGGCAGCTGCCATGGGTCAGAAGAAGCGTTGGGCAAAGGGTGCTGTGCAGATTCTGCTGATGAAAAATGAGAGTGAAGTCGACCCGGACTGGCGTCCACCGCGTGTGCCTGCCCCGGACCCGAAGCCGTCTCTTACGTTCCCGCGTAAGATGTTCTTTGTGCGAAGGTGCGGTACCTGAAACAGTGGCTGCTGCCATGGGTCAAAAGAAGCGTTGGGCAAAGGGTGCTGTGCAGATTCTGTTGATGAAGAATGAGAGCGAGGTTGACCCGGACTGGCGTCCGCCGCGCGTTCCTGCCCCAGACCCGAAGCCGTCGCTAACGTTCCCACGTAAGATGTTCTTTGTGTGAAGGTGCTGTGCCCGAGACCGTCGCTGCAGCCATGGGTCAGAAGAAACGTTGGGCCAAGGGTGCCGTGCAGATTCTGCTCATGAAAAGTGAAAGCGAGGTCGACCCGGATTGGCGTCCACCTCGCGTTCCTGCCCCGGACCCGAAGCCGTCGCTTGCGTTTCCACGTAAAATGTTTTTTGTGTGAAGGTGCCGTGCCCGAAACGGTCGCTGCCGCCATGGGTCAGAAGAAGCGTTGGGCCAAGGGTGCCGTGCAGATTCTGCTGATGAAAAATGAAAGCGAGGTGGACCCAGACTGGCGTCCGCCGCGTGTGCCTGCCCCGGACCCGAAGCCGTCGCTCGCGTTCCCGCGTAAGATGTTCTTTGTGCGAAGGTGCCGTGCCCGAAACGGTCGCTGCTGCCATGGGTCAGAAGAAGCGTTGGGCCAAGGGTGCCGTGCAGATTCTGCTCATGAAGAATGAGAGCGAGGTCGACCCGGACTGGCGTCCGCCGCGTGTGCCTGCCCCGGACCCGAAGCCGGCGCTTGCGTTCCCGCGCAAGATGTTCTTTGTGTGAAGGTGCCGTGCCCGAGACAGTCGCTGCTGCTATGGGTCAGAAGAAACGTTGGGCCAAGGGTGCCGTCCAGATTCTGTTGATGAAGAATGAAAGCGAAGTCGACCCGGACTGGCGTCCGCCGCGTGTGCCTGCCCCCGACCCGAAGCCGTCGCTCGCGTTCCCGCGTAAGATGTTCTTTGTGCGAAGGAGCTGTGCCCGACACGGTCGCTGCTGCCATGGGACAGAAGAAACGTTGGGCCAAGGGTGCCGTGCAGATTCTGCTGATGAAGAATGAGAGCGAAGTCGACCCGGACTGGCGTCCCCCGCGTGTGCCTGCCCCTGACCCGAAGCCGTCGCTCACGTTCCCGCGTAAGATGTTCTTTGTGTGAAGGTGCGGTGCCCGAAACTGTGGCTGCAGCCATGGGCCAGAAGAAACGTTGGGCTAAGGGTGCTGTCCAAATTCTACTTATGAAAAATGAGAGTGAGGTCGACCCGGACTGGCGTCCACCGCGCGTGCCGGCTCCGGACCCAAAGCCGTCGCTAGCTTTTCCGCGTAAGATGTTCTTTGTG+GAAGGTGCCGTGCCCGAAACGGTCGCTGCTGCCATGGGTCAGAAGAA+CGTTGGGCCAAGGGTGCCGTGCAGATTCTGCTGATGAA+AATGAGAGCGAGGTCGACCCGGACTGGCGTCCGCCGCGTGTGCCTGCCCCGGACCCGAAGCCGTCGCT+GCGTTCCCGCGTAAGATGTTCTTCTACGACTCGGTGCTGTACCCGTTCGGTTCAATTCCTGCCTTGTGTTACGTGGCAATCGCTGTCTACTACTTGTGCACGGGAGACGCTCCGATCTACGCTCGTGGAACCAAGTTCCTGTACTCTTTCTTGCCCGTGACGTTCTGCCGTTGGGTGCTGAATCTGTTGGCGAACCGCGCTGTTGATCTACGATTCGGTTCTGTACCCGTTCGGCTCGATCCCTGCCTTGTGTTACGTGTCGATCGCTGTCTACTACTTATGTACGGGTGACGCTCCGATTTATGCTCGTGGTACCAAGTTTTTGTACTCTTTCTTGCCCGTGACGTTCTGTCGTTGGGTGCTGAATTTGTTGGCTAACCGCGCTGTCGACCTACGATTCGGTGCTGTACCCTTTCGGCTCAATTCCGGCGCTGTGTTACGTCGCGATTGCTGTGTACTACTTGTGCACGGGTGATGCGCCTATCTACGCGCGTGGTACTAAGTTTATCTACTCTTTCTTACCCGTGACGTTCTGTCGTTGGGTGCTGAACTTGCTCGCGAATCGTGCTGTTGACCTACGACTCGGTGCTGTACCCGTTCGGTTCGATCCCCGCTCTGTGTTACGTGGCTATCGCTATCTACTACCTCTGCACGGGTGACGCTCCTATCTACGCTCGCGGTACCAAGTTCCTGTACTCTTTCTTGCCCGTGACGTTCTGCCGTTGGGTGCTCAACTTGCTGGCCAACCGCGCCGTCGACCTACGACTCGGTGCTCTACCCGTTCGGTTCCATTCCCGCTCTGTGTTACGTGGCGATCGCTATTTACTACCTGTGTACGGGTGACGCTCCCATCTACGCTCGTGGTACCAAGTTCCTGTACTCTTTCTTGCCCGTGACGTTCTGCCGTTGGGTACTCAACCTGCTGGCCAACCGCGCCGTCGACCTATGACTCGGTGCTGTACCCGTTCGGTTCGATCCCTGCCTTGTGTTACGTGTCGATCGCCGTCTACTACCTGTGCACGGGTGACGCTCCCATTTACGCTCGTGGTACCAAGTTCCTGTACTCTTTCTTGCCCGTGACGTTCTGCCGTTGGGTGCTCAACCTGCTCGCTAACCGCGCTGTCGACCTACGACTCGGTGCTGTATCCGTTCGGCTCCATCCCGGCCCTGTGTTACGTGTCGATCGCCGTCTACTACCTGTGCACGGGCGACGCCCCCATTTACGCGCGTGGAACCAAGTTCCTGTACTCTTTCTTGCCCGTGACGTTCTGTCGCTGGGTGCTGAACTTGCTCGCTAACCGCGCAGTGGACCTACGATTCAGTGCTTTATCCGTTCGGTTCGATCCCTGCTCTATGCTACGTGGCAATCGCCGTCTACTACCTGTGCACAGGTGATGCACCCATCTACGCACGCGGTACAAAGTTTCTCTATTCTTTCTTGCCTGTGACTTTCTGTCGTTGGGTACTCAACCTGCTGGCCAATCGTGCTGTCGATCTACGACTCGGTGCTGTACCCGTTCGGTTCGATCCCTGCCCTGTGTTACGTGGCGATCGCTGTCTACTACCTGTGCACGGGTGACGCTCCCATCTACGCTCGTGGTACCAAGTTCCTGTACTCTTTCTTGCCCGTGACGTTCTG+CGTTGGGTGCT+AAC+TGCTGGC+AACCGCGCTGTCGACAACAACGACGTGTGGCGTGCGCAGCAGACATGGTTTTCCTTCTCGTTCATTACGATGATGGCAATTTTTGAGGCCATCCAGGCGCGCTTGACGGGTAAGGACAAGTCATGGGCAAACACGGGTGCTGGTCAAAAGACGTCGTGGACGGAAATCCCAAACGTGCTCTTCTTTTTCACGCTGCTGTAACAATGACGTGTGGCGTGCGCAGCAGACGTGGTTTTCTTTTTCGTTCATCACGATGATGGCTATCATTGAAGCCATCCAGGCGCGAATGACGGGCAAGGACAAATCTTGGGCAAACACAGGTGCGGGTCAGAAGACGTCGTGGACAGAGATTCCGAACGTGCTTTTCTTCTTTACGCTGATGTAACAATGATGTGTGGCGTGCACAGCAGACGTGGTTTTCCTTCTCTTTTATTACGATGATGGCCATTGTGGAGGCTATTCAGGCTCGTGCGACGGGAAAGGATAAGTCGTGGGCGAACACTGGTGCTGGGCAGAAGACATCATGGACGGAAATCCCTAATGTGCTGTTCTTTTTCACGCTTATGTAACAACGATGTGTGGCGTGCCCAGCAGACGTGGTTCTCCTTCTCCTTCATCACGATGATGGCTATCGTGGAGGCCATCCAGGCGCGTGTGACGGGCAAGGACAAGTCGTGGGCCAACACGGGTGCCGGTCAGAAGACGTCGTGGACGGAAATCCCCAACGTGCTGTTCTTCTTCACGCTGCTCTAACAACGATGTGTGGCGTGCCCAGCAGACCTGGTTCTCCTTCTCCTTCATCACGATGATGGCTATTGTGGAGGCTATCCAGGCGCGTGTGACGGGCAAAGACAAGTCGTGGGCCAACACGGGTGCCGGTCAGAAGACGTCGTGGACAGAGATCCCCAACGTGCTCTTCTTCTTCACGCTGCTCTAACAACGACGTGTGGCGTGCACAGCAGACGTGGTTCTCATTCTCGTTCATCACAATGATGGCTATTGTGGAGGCTATTCAGGCGCGTGTGACGGGCAAGGACAAGTCGTGGGCTAACACTGGTGCCGGTCAGAAGACGTCATGGACGGAAATCCCCAACGTGCTGTTCTTCTTCACGCTGCTGTAACAACGACGTGTGGCGCGCCCAGCAGACGTGGTTCTCGTTCTCGTTCATTACGATGATGGCTATTGTGGAGGCCATTCAGGCGCGTGTGACAGGCAAGGACAAGTCGTGGGCCAACACGGGTGCCGGTCAGAAGACGTCGTGGACAGAGATTCCCAACGTTCTCTTCTTCTTCACGCTGCTCTAACAATGACGTGTGGCGTGCTCAGCAGACGTGGTTCTCCTTCTCATTTATCACGATGATGGCTATCGTTGAGGCTATACAAGCGCGTGTGACGGGCAAAGACAAATCGTGGGCTAATACTGGTGCAGGACAGAAAACGTCCTGGACGGAAATTCCCAACGTTTTGTTTTTCTTCACACTACTTTAACAACGACGTGTGGCGTGCCCAGCAGACGTGGTTCTCCTTCTCGTTCATCACGATGATGGCTATTGTGGAGGC+ATCCAGGCGCGTGTGACGGGCAAGGACAAGTCGTGGGCCAACACGGGTGCCGGTCAGAAGACGTCGTGGACGGAAATCCCCAACGTGCTGTTCTTCTTCACGCTGCTGTTCAGTCAACTCGTGGCGCTCATTCGTTTCTTCGAGTACGAGAACGCAACGAACCCGTGGAACTACGTGTCTGCGATGTTCTTTGGCTTTTTCGTCATGAGCCAGTTCTACCCCATGGTCAAGATGAGTATTACCGAGTACTGTGGTTGGGACCATACTGCTGCCACCTTCACGGCGAACGTGTTTTAGTCAGTTGGTAGCGCTCATCCGCTTCTTCGAGTATGAGAACGCCACGAATCCATGGAATTACGTGTCTTCCATGTTCTTTGGCTTCTTCGTCATGAGCCAGTTCTACCCCATGGTCAAGATGAGTATCACGGAGTACTGTGGCTGGGACCACACTGCCGCTACATTTACGGCGAACGTGTTTTAGCCAAGTCGTGGCGCTCATTCGATTCTTTGAGTACGAGAATGCAACGAATCCGTGGAATTACGTGTCAGCAATGTTTTTTGGCTTTTTCGTCATGAGCCAATTCTATCCCATGGTCAAGATGAGTATCACGGAATATTGTGGATGGGACCACACGGCCGCCACGTTCACGGCCAATGTCTTTCAGTCAGTTGGTGGCGCTGATCCGATTCTTCGAGTACGAGAACGCCACGAACCCGTGGAACTACGTGTCTGCCATGTTCTTCGGCTTCTTCGTCATGAGCCAGTTCTACCCTATGGTCAAGATGAGTATCACGGAGTACTGTGGCTGGGACCACACGGCCGCGACGTTCACGGCCAACGTGTTTTAGTCAACTGGTGGCGCTGATTCGGTTCTTTGAGTACGAGAACGCCACGAACCCGTGGAACTACGTGTCTGCTATGTTCTTCGGCTTCTTCGTGATGAGTCAGTTCTACCCCATGGTCAAGATGAGTATCACGGAGTACTGTGGTTGGGACCACACGGCCGCGACATTTACGGCCAACGTGTTTCAGTCAAGTGGTGGCGCTGGTGCGATTCTTCGAGTATGAGAACGCCACGAACCCGTGGAACTACGTGTCTGCCATGTTCTTTGGCTTCTTCGTCATGAGCCAGTTCTACCCCATGGTGAAGATGAGTATCACGGAATACTGTGGTTGGGACCACACGGCTGCCACCTTCACGGCCAACGTGTTTCAGTCAACTGGTGGCGCTGGTGCGTTTCTTTGAGTACGAGAACGCCACGAACCCGTGGAACTACGTGTCTGCCATGTTCTTTGGCTTCTTCGTCATGAGCCAGTTCTACCCCATGGTCAAGATGAGTATCACAGAATACTGTGGCTGGGACCACACGGCTGCCACGTTCACGGCGAACGTGTTTCAGTCAGGTGGTGGCCTTGATTCGATTCTTCGAGTATGAAAATGCAACAAATCCCTGGAACTACGTATCTGCCATGTTTTTTGGCTTTTTTGTCATGAGCCAATTCTACCCCATGGTCAAGATGAGTATCACTGAATATTGTGGATGGGACCACACTGCTGCTACCTTTACCGCAAATGTGTTTCAGTCAA+TGGTGGCGCTGATTCGATTCTTCGAGTACGAGAACGCCACGAACCCGTGGAACTACGTGTCTGCCATGTTCTTTGGCTTCTTCGTCATGAGCCAGTTCTACCCCATGGTCAAGATGAGTATCACGGA+TACTGTGG+TGGGACCACACGGC+GCCAC+TTCACGGCCAACGTGTTTGGTTCGCTGCTGGTGGTCTACATCGTGGTCTTCGTGCAGCTATGGCAAGTGTACTACGCGGGTCGCTGCTCGTGGTCTACATCGTGGTGTTTGTGCAGTTGTGGCAGGTCTACTACGCGGCTCCTTGCTGGTCGTATACGTTGTCGTGTTCGTACAGCTATGGCAAGTGTATTACGCGGCTCGCTGCTGGTGGTGTACATTGTGGTATTCGTGCAGCTGTGGCAGGTCTACTACGCGGCTCGCTGCTGGTGGTGTACATCGTGGTGTTCGTGCAGCTGTGGCAGGTCTACTACGCGGCTCGTTGCTGGTGGTGTACGTGGTGGTGTTCGTGCAGCTGTGGCAAGTGTACTACGCGGCTCGCTGCTGGTGGTGTACATCGTGGTGTTTGTGCAGCTGTGGCAGGTGTACTACGCGGCTCGTTATTGGTCGTTTACGTTGTCGTGTTTGTGCAGCTATGGCAAGTCTACTACGCGGCTCGCTGCTGGTGGTGTACATCGTGGTGTTCGTGCAGCTGTGGCA+GT+TACTACG1111111118318318318318318318318310203040506070809010011012013014015016017018018418418418418418418418436736736736736736736736719020021022023024025026027028029030031032033034035036036836836836836836836836855155155155155155155155137038039040041042043044045046047048049050051052053054055055255255255255255255255273573573573573573573573556057058059060061062063064065066067068069070071072073073673673673673673673673691991991991991991991991974075076077078079080081082083084085086087088089090091092092092092092092092092011031103110311031103110311031103930940950960970980990100010101020103010401050106010701080109011001104110411041104110411041104110412871287128712871287128712871287111011201130114011501160117011801190120012101220123012401250126012701280128812881288128812881288128812881471147114711471147114711471147112901300131013201330134013501360137013801390140014101420143014401450146014701472147214721472147214721472147216551655165516551655165516551655148014901500151015201530154015501560157015801590160016101620163016401650165616561656165616561656165616561839183918391839183918391839183916601670168016901700171017201730174017501760177017801790180018101820183018401840184018401840184018401840202320232023202320232023202320231850186018701880189019001910192019301940195019601970198019902000201020202024202420242024202420242024202422072207220722072207220722072207203020402050206020702080209021002110212021302140215021602170218021902200220822082208220822082208220822082391239123912391239123912391239122102220223022402250226022702280229023002310232023302340235023602370238023902392239223922392239223922392239225752575257525752575257525752575240024102420243024402450246024702480249025002510252025302540255025602570257625762576257625762576257625762759275927592759275927592759275925802590260026102620263026402650266026702680269027002710272027302740275027602760276027602760276027602760294329432943294329432943294329432770278027902800281028202830284028502860287028802890290029102920293029402944294429442944294429442944294431273127312731273127312731273127295029602970298029903000301030203030304030503060307030803090310031103120312831283128312831283128312831283311331133113311331133113311331131303140315031603170318031903200321032203230324032503260327032803290330033103312331233123312331233123312331233703370337033703370337033703370332033303340335033603370
