## Supplemental Figure 2 for "Advancing RNAi-Based Strategies Against Downy Mildews: Insights Into dsRNA Uptake and Gene Silencing": Supplemental Figure 2.html

Launch in Jalview  

{"seqs":[{"start":1,"name":"Zebrafish/1-1288","end":1288,"id":"1877257810","seq":"ATGAGGGAGATCGTGCATTTACAGGCTGGACAGTGCGGCAACCAGATTGGTGCCAAGTTCTGGGAAGTCATTAGTGACGAGCATGGAATTGACCCAACAGGCAGTTACCATGGCGACAGTGACCTTCAGCTGGACCGAATTAATGTTTATTATAATGAAGCCACAGGTGGAAAGTATGTTCCACGTGCCGTGCTGGTGGATTTGGAGCCTGGTACAATGGACTCCGTGAGGTCTGGTCCATTTGGTCAGATCTTCAGACCAGACAACTTTGTGTTCGGCCAGAGTGGTGCTGGAAACAACTGGGCCAAGGGCCACTACACTGAAGGAGCTGAGCTGGTTGATTCCGTTCTGGATGTGGTCCGAAAAGAGGCTGAGAGCTGCGACTGTCTGCAGGGCTTCCAACTCACTCACTCACTGGGTGGAGGTACAGGGTCTGGTATGGGCACCCTCCTCATTAGCAAAATCCGCGAGGAGTATCCCGACCGCATCATGAACACCTTCAGCGTGGTGCCCTCTCCTAAAGTCTCGGACACTGTGGTCGAGCCCTACAACGCCACACTGTCCGTCCATCAGCTAGTAGAGAACACAGACGAGACCTATTGTATTGATAACGAGGCCCTGTACGATATCTGCTTCCGCACACTCAAACTCACAACCCCCACATACGGAGACCTCAACCATCTCGTCTCCGCCACAATGAGCGGTGTGACCACTTGCTTGAGGTTTCCAGGCCAGTTGAACGCTGATCTCCGTAAATTGGCGGTCAACATGGTGCCCTTCCCCCGACTGCACTTCTTCATGCCTGGCTTCGCGCCTCTGACTAGCAGGGGAAGCCAGCAGTATCGTGCACTTACCGTTCCCGAACTCACCCAGCAGATGTTCGATGCCAAAAACATGATGGCTGCCTGCGACCCACGTCACGGCCGTTATCTGACGGTCGCCGCCGTCTTCCGTGGTCGCATGTCCATGAAGGAGGTGGACGAGCAGATGCTCAACGTCCAGAACAAGAACAGCAGCTACTTCGTTGAATGGATCCCAAACAACGTCAAGACCGCCGTCTGCGACATTCCACCACGAGGCCTCAAGATGGCCGCCACCTTCATCGGCAACAGCACCGCCATCCAGGAGCTCTTCAAGCGCATCTCAGAGCAGTTCACGGCTATGTTCAGGCGCAAGGCTTTCCTGCATTGGTACACCGGAGAGGGCATGGATGAGATGGAGTTCACAGAGGCCGAGAGCAACATGAACGACCTGGTGTCCGAGTACCAGCAGTACCAGGACGCCACCG","order":1},{"start":1,"name":"Arabidopsis/1-1288","end":1288,"id":"440441233","seq":"ATGAGAGAGATCCTTCATATCCAAGGCGGTCAATGTGGAAACCAGATCGGAGCAAAGTTCTGGGAAGTGATCTGCGACGAACACGGCATTGATCACACCGGTCAATACGTCGGCGATTCTCCGTTACAGCTTGAACGTATCGATGTCTATTTCAACGAAGCTAGCGGTGGAAAGTACGTTCCTCGCGCTGTTCTTATGGATCTGGAGCCTGGTACCATGGATTCTCTCAGATCTGGTCCGTTCGGTCAGATTTTCCGTCCTGATAACTTCGTCTTTGGTCAATCTGGTGCCGGAAATAACTGGGCGAAAGGTCATTACACCGAAGGTGCTGAGTTGATTGATTCTGTTCTCGATGTTGTGAGGAAGGAAGCTGAGAACAGCGATTGTCTTCAAGGTTTCCAAGTGTGTCATTCATTGGGAGGAGGAACTGGATCTGGAATGGGAACTCTATTGATTTCTAAGATAAGAGAAGAGTATCCAGATCGTATGATGATGACTTTCTCAGTGTTTCCTTCTCCTAAGGTCTCTGACACTGTTGTTGAGCCATACAATGCAACTCTCTCTGTGCATCAGCTTGTCGAAAACGCTGACGAGTGTATGGTTTTGGACAATGAGGCTCTCTACGATATCTGTTTCCGTACCCTCAAGCTCGCTAATCCTACCTTTGGTGATCTTAACCATCTCATCTCTGCTACAATGAGTGGTGTCACTTGCTGTCTTCGTTTCCCTGGCCAGCTTAACTCTGACCTTAGGAAACTCGCTGTGAACCTTATCCCATTCCCAAGGCTTCACTTCTTCATGGTTGGTTTCGCACCATTGACATCGAGAGGATCACAGCAATACAGTGCCTTGAGTGTTCCTGAACTGACCCAGCAGATGTGGGATGCAAAGAACATGATGTGTGCTGCTGACCCTCGTCATGGACGTTACTTGACTGCATCCGCTGTGTTCCGTGGAAAGCTGAGCACCAAAGAGGTTGACGAGCAGATGATGAACATTCAGAACAAGAACTCATCCTACTTTGTGGAATGGATCCCAAACAACGTCAAGTCCAGTGTCTGTGATATTGCACCAAAGGGTTTGAAAATGGCGTCTACTTTCATTGGTAACTCAACCTCAATCCAGGAGATGTTTAGGCGTGTGAGCGAACAGTTCACAGCTATGTTCAGGAGAAAGGCTTTCCTTCATTGGTACACAGGAGAAGGCATGGACGAGATGGAGTTCACTGAAGCAGAGAGTAACATGAATGATCTTGTCGCAGAGTACCAGCAGTACCAAGATGCTACAG","order":2},{"start":1,"name":"Human/1-1288","end":1288,"id":"265885290","seq":"ATGAGGGAAATCGTGCACATCCAGGCTGGTCAGTGTGGCAACCAGATCGGTGCCAAGTTCTGGGAGGTGATCAGTGATGAACATGGCATCGACCCCACCGGCACCTACCACGGGGACAGCGACCTGCAGCTGGACCGCATCTCTGTGTACTACAATGAAGCCACAGGTGGCAAATATGTTCCTCGTGCCATCCTGGTGGATCTAGAACCTGGGACCATGGACTCTGTTCGCTCAGGTCCTTTTGGCCAGATCTTTAGACCAGACAACTTTGTATTTGGTCAGTCTGGGGCAGGTAACAACTGGGCCAAAGGCCACTACACAGAGGGCGCCGAGCTGGTTGATTCTGTCCTGGATGTGGTACGGAAGGAGGCAGAGAGCTGTGACTGCCTGCAGGGCTTCCAGCTGACCCACTCACTGGGCGGGGGCACAGGCTCTGGAATGGGCACTCTCCTTATCAGCAAGATCCGAGAAGAATACCCTGATCGCATCATGAATACCTTCAGTGTGGTGCCTTCACCCAAAGTGTCTGACACCGTGGTCGAGCCCTACAATGCCACCCTCTCCGTCCATCAGTTGGTAGAGAATACTGATGAGACCTATTGCATTGACAACGAGGCCCTCTATGATATCTGCTTCCGCACTCTGAAGCTGACCACACCAACCTACGGGGATCTGAACCACCTTGTCTCAGCCACCATGAGTGGTGTCACCACCTGCCTCCGTTTCCCTGGCCAGCTCAATGCTGACCTCCGCAAGTTGGCAGTCAACATGGTCCCCTTCCCACGTCTCCATTTCTTTATGCCTGGCTTTGCCCCTCTCACCAGCCGTGGAAGCCAGCAGTATCGAGCTCTCACAGTGCCGGAACTCACCCAGCAGGTCTTCGATGCCAAGAACATGATGGCTGCCTGTGACCCCCGCCACGGCCGATACCTCACCGTGGCTGCTGTCTTCCGTGGTCGGATGTCCATGAAGGAGGTCGATGAGCAGATGCTTAACGTGCAGAACAAGAACAGCAGCTACTTTGTGGAATGGATCCCCAACAATGTCAAGACAGCCGTCTGTGACATCCCACCTCGTGGCCTCAAGATGGCAGTCACCTTCATTGGCAATAGCACAGCCATCCAGGAGCTCTTCAAGCGCATCTCGGAGCAGTTCACTGCCATGTTCCGCCGGAAGGCCTTCCTCCACTGGTACACAGGCGAGGGCATGGACGAGATGGAGTTCACCGAGGCTGAGAGCAACATGAACGACCTCGTCTCTGAGTATCAGCAGTACCAGGATGCCACCG","order":3},{"start":1,"name":"Worm/1-1288","end":1288,"id":"609353849","seq":"ATGCGTGAAATTGTTCATATCCAGGCAGGTCAGTGTGGTAACCAAATTGGGGCAAAGTTCTGGGAAGTTATTTCCGACGAGCACGGGATCGATCCCACCGGAGCATACAATGGAGACTCCGATTTGCAGTTGGAGAGAATCAATGTCTACTACAACGAAGCTAGCGGAGGAAAGTATGTCCCACGTGCTTGTCTTGTTGATTTGGAGCCAGGAACCATGGACTCCGTTAGGGCAGGTCCCTTTGGACAGCTCTTTCGACCAGACAACTTTGTTTTTGGTCAAAGCGGTGCCGGAAACAACTGGGCCAAAGGTCACTACACCGAGGGCGCCGAACTTGTAGACAATGTTCTCGACGTTGTCCGTAAAGAAGCTGAAAGCTGTGATTGTCTTCAGGGATTCCAAATGACTCACTCTTTGGGAGGAGGAACTGGTTCTGGAATGGGAACTCTTTTGATTTCCAAAATTCGTGAAGAATATCCAGATCGTATCATGATGACTTTCTCCGTAGTGCCAAGTCCAAAAGTGTCCGACACGGTCGTCGAGCCGTACAACGCAACTCTTTCTGTTCATCAACTTGTTGAAAACACCGACGAAACATTCTGTATTGACAACGAAGCCTTGTATGACATCTGTTTCCGCACGCTCAAGCTCACCACACCGACCTACGGAGATTTGAATCATCTCGTTTCGATGACGATGAGTGGTGTCACCACCTGTCTTCGTTTCCCGGGACAGCTGAATGCAGATTTGCGCAAATTAGCTGTTAACATGGTTCCGTTCCCACGTCTTCATTTCTTCATGCCCGGATTTGCTCCGCTCACATCCAGAGGAAGCCAGCAATACAGATCGCTCACCGTTCCAGAGCTCACACAACAAATGTTCGACGCCAAGAATATGATGGCAGCCTGCGATCCAAGACACGGTCGCTACCTGACAGTTGCTGCAATGTTCCGCGGAAGAATGAGCATGAAAGAAGTCGACGAGCAAATGCTCAATGTGCAAAATAAAAACTCCTCGTACTTTGTTGAATGGATTCCAAACAACGTCAAAACTGCTGTTTGTGATATTCCGCCAAGAGGAGTAAAGATGGCGGCAACATTCGTCGGAAATTCCACTGCAATTCAAGAGCTTTTCAAGAGAATCAGTGAGCAATTCACAGCTATGTTCCGTAGAAAAGCTTTCCTTCATTGGTACACTGGAGAAGGTATGGACGAAATGGAGTTTACTGAAGCTGAGAGTAACATGAATGACTTGGTCTCAGAGTATCAACAGTATCAAGAAGCAACCG","order":4},{"start":1,"name":"Yeast/1-1288","end":1288,"id":"1854848159","seq":"ATGAGAGAAATCATTCATATCTCGACAGGTCAGTGTGGTAACCAAATTGGTGCTGCATTCTGGGAAACTATCTGTGGTGAGCACGGTTTGGATTTCAATGGGACATATCACGGCCATGACGATATCCAGAAGGAGAGACTGAACGTGTACTTCAACGAGGCATCTTCTGGGAAGTGGGTTCCAAGATCTATTAACGTCGATCTAGAACCTGGGACGATTGACGCAGTACGCAATTCTGCCATCGGGAATTTGTTTAGACCTGACAATTATATCTTTGGGCAAAGTTCTGCGGGCAACGTGTGGGCCAAGGGTCACTACACAGAAGGTGCTGAGCTTGTAGACAGCGTCATGGATGTTATTAGACGAGAGGCCGAAGGATGCGACTCCCTTCAAGGTTTCCAGATCACACATTCTCTTGGTGGTGGTACCGGTTCCGGTATGGGTACGCTTTTGATCTCGAAGATTAGGGAAGAGTTTCCTGATCGTATGATGGCCACCTTCTCCGTCTTGCCCTCTCCGAAGACTTCTGACACCGTTGTCGAACCATACAATGCCACGTTGTCTGTGCACCAATTGGTAGAACACTCTGATGAAACATTCTGTATCGATAACGAAGCACTTTATGACATCTGTCAAAGGACCTTAAAGTTGAATCAACCTTCTTATGGAGATTTGAACAACTTGGTCTCGAGCGTCATGTCTGGTGTGACAACTTCATTGCGTTATCCCGGCCAATTGAACTCTGATTTGAGAAAGTTGGCTGTTAATCTTGTCCCATTCCCACGTTTACATTTCTTCATGGTCGGCTACGCTCCATTGACGGCAATTGGCTCTCAATCATTTAGATCTTTGACTGTCCCTGAATTAACACAGCAAATGTTTGATGCCAAGAACATGATGGCTGCTGCCGATCCAAGAAACGGTAGATACCTTACCGTTGCAGCCTTCTTTAGAGGTAAAGTTTCCGTTAAGGAGGTGGAAGATGAAATGCATAAAGTGCAATCTAAAAACTCAGACTATTTCGTGGAATGGATCCCCAACAATGTGCAAACTGCTGTGTGTTCTGTCGCTCCTCAAGGTTTGGACATGGCTGCTACTTTCATTGCTAACTCCACATCTATTCAAGAGCTATTCAAGAGAGTTGGTGACCAATTTTCCGCTATGTTCAAAAGAAAAGCTTTCTTGCACTGGTATACTAGTGAAGGTATGGACGAATTGGAATTCTCTGAGGCTGAATCTAATATGAATGATCTGGTTAGCGAATACCAACAATACCAAGAGGCTACTG","order":5},{"start":1,"name":"Hy\_brassicae/1-1288","end":1288,"id":"1889287374","seq":"ATGCGCGAGCTCGTCCACATTCAGGGCGGCCAGTGCGGCAACCAGATCGGCGCCAAGTTCTGGGAAGTCATTTCCGACGAGCACGGCGTGGACCCCACGGGCTCGTACCGCGGCGACTCGGACCTGCAGCTCGAGCGCATCAACGTGTACTACAACGAAGCGACGGGCGGACGCTATGTGCCCCGCGCGATTCTGATGGACCTGGAGCCCGGCACGATGGACTCTGTCCGCGCAGGTCCGTACGGCCAGCTCTTCCGTCCGGACAACTTTGTGTTTGGACAGACAGGCGCGGGTAACAACTGGGCCAAGGGACACTATACGGAGGGTGCGGAGCTCATTGATTCGGTGCTGGACGTTGTCCGTAAAGAGGCCGAGAGCTGTGACTGCCTACAGGGTTTCCAGTTCACGCACTCGCTTGGCGGTGGTACTGGCTCGGGTATGGGCACGCTTCTTATCTCGAAGATCCGCGAGGAATACCCGGATCGCATCATGTGCACGTACTCCGTGTGCCCGTCCCCGAAGGTGTCGGACACGGTGGTGGAGCCCTACAATGCGACGCTGTCGGTGCACCAGCTCGTTGAAAACGCTGACGAAGTCATGTGCCTGGACAACGAGGCCCTGTACGACATTTGCTTCCGTACGCTGAAACTCACCACCCCTACGTATGGTGACCTGAACCACTTGGTGTGCGCTGCTATGTCTGGCATCACGACGTGTCTACGTTTTCCCGGCCAGCTGAACTCCGACCTGCGTAAGCTAGCCGTGAACCTGATTCCGTTTCCACGTCTCCACTTCTTCATGATCGGATTCGCTCCGCTGACGTCGCGAAGCTCGCAGCAGTACCGTGCGCTGACGGTGCCCGAGCTCACGCAGCAGCAGTTTGATGCGAAGAACATGATGTGCGCTGCAGACCCTCGCCATGGCCGTTATTTGACCGCCGCGTGTATGTTCCGCGGTCGCATGAGTACGAAGGAGGTGGACGAGCAAATGCTGAACGTGCAGAACAAGAATTCGTCGTACTTCGTCGAGTGGATTCCGAATAACATTAAGGCTAGCGTGTGCGACATCCCGCCCAAGGGCCTGAAGATGAGTACGACGTTTATCGGCAACTCGACCGCGATCCAGGAGATGTTCAAGCGTGTGTCTGAACAGTTTACGGCTATGTTTCGTCGTAAGGCTTTCTTGCACTGGTACACTGGCGAGGGTATGGATGAGATGGAGTTCACGGAGGCCGAGTCGAACATGAACGATCTCGTGTCTGAGTACCAACAGTACCAGGATGCCACCG","order":6},{"start":1,"name":"Ph\_ramorum/1-1288","end":1288,"id":"934354673","seq":"ATGAGAGAGCTCGTTCACATCCAGGGTGGCCAGTGCGGTAACCAGATCGGCGCCAAGTTCTGGGAGGTTATCTCCGACGAGCACGGCGTGGACCCCACGGGCTCGTACCACGGCGACTCGGACCTGCAGCTGGAGCGCATCAATGTGTACTACAACGAGGCCACGGGCGGCCGCTACGTGCCCCGCGCCATCCTCATGGACCTGGAGCCCGGCACCATGGACTCGGTCCGCGCCGGCCCCTACGGCCAGCTCTTCCGCCCGGACAACTTCGTGTTCGGTCAGACCGGCGCCGGTAACAACTGGGCTAAGGGACACTACACGGAGGGTGCCGAGCTTATCGACTCGGTGCTCGACGTCGTCCGCAAGGAGGCCGAGAGCTGTGACTGCCTGCAGGGGTTCCAGATCACGCACTCTCTTGGTGGCGGTACCGGTTCTGGTATGGGCACGCTTCTGATCTCCAAGATCCGTGAGGAGTACCCGGACCGTATCATGTGCACGTACTCGGTGTGCCCGTCGCCCAAGGTGTCGGACACGGTCGTGGAGCCCTACAACGCCACGCTGTCGGTGCACCAGCTTGTCGAGAACGCCGACGAGGTCATGTGCCTGGATAACGAGGCGCTGTACGACATTTGCTTCCGCACGCTCAAGCTCACCACCCCCACCTACGGTGACCTGAACCACCTGGTGTGCGCCGCTATGTCCGGCATCACCACGTGCCTGCGTTTCCCGGGTCAGCTGAACTCGGACCTGCGGAAGCTGGCGGTGAACTTGATTCCGTTCCCGCGTCTTCACTTCTTCATGATTGGTTTCGCCCCGCTGACCTCGCGTGGCTCGCAGCAGTACCGTGCCCTGACGGTGCCCGAGCTGACCCAGCAGCAGTTCGACGCAAAGAACATGATGTGCGCCGCCGACCCTCGTCACGGCCGCTATTTAACTGCCGCGTGTATGTTCCGCGGACGTATGAGCACGAAGGAGGTTGATGAGCAGATGCTGAACGTGCAGAACAAGAACTCGTCGTACTTCGTCGAGTGGATCCCTAACAACATCAAGGCTAGCGTGTGTGACATCCCGCCCAAGGGCCTGAAGATGAGCACTACGTTCATCGGTAACTCGACCGCTATCCAGGAGATGTTCAAGCGTGTGTCTGAGCAGTTTACGGCTATGTTCCGTCGTAAGGCTTTCTTGCACTGGTACACGGGCGAGGGTATGGACGAGATGGAGTTCACGGAGGCCGAGTCCAACATGAACGATCTTGTGTCTGAGTACCAGCAGTACCAGGACGCCACCG","order":7},{"start":1,"name":"Ph\_cinnamomi/1-1288","end":1288,"id":"64258486","seq":"ATGAGAGAGCTCGTTCACATCCAGGGTGGCCAGTGCGGTAACCAGATCGGCGCCAAGTTCTGGGAGGTCATCTCCGACGAGCACGGCGTGGACCCGACGGGATCCTACCACGGCGACTCGGACCTGCAGCTGGAGCGCATCAACGTGTACTACAACGAGGCCACGGGCGGCCGCTACGTGCCCCGCGCCATCCTCATGGACCTGGAGCCCGGCACCATGGACTCGGTGCGCGCCGGCCCCTACGGCCAGCTCTTCCGCCCGGACAACTTCGTGTTCGGCCAGACGGGCGCCGGTAACAACTGGGCCAAGGGACACTACACGGAGGGCGCCGAGCTCATCGACTCGGTGCTCGATGTCGTCCGCAAGGAGGCGGAGAGCTGCGACTGCCTGCAGGGATTCCAGATCACGCACTCGCTCGGTGGCGGTACCGGTTCCGGTATGGGCACGCTTCTTATCTCCAAGATCCGTGAGGAGTACCCGGACCGTATCATGTGCACGTACTCGGTGTGCCCGTCGCCCAAGGTGTCGGACACGGTCGTGGAGCCCTACAACGCGACGCTGTCCGTGCACCAGCTTGTCGAGAACGCCGATGAGGTCATGTGCCTGGATAACGAGGCCCTGTACGACATTTGCTTCCGCACCTTGAAATTGACGAACCCGACGTACGGTGATCTGAACCACCTGGTGTGCGCCGCCATGTCCGGCATCACCACGTGCCTCCGTTTCCCCGGCCAACTGAACTCGGTCCTGAAGCTGTTTGCCGTTAACCTGATCCCGTTCCCCCGTCTCCATTTCTTCATGATCGGTTTCGCTCCGCTGACGTCGCGTGGATCTCAGCAGTACCGTGCCCTGACGGTGCCGGAGCTGACGCAGCAGCAGTTCGATGCTAAGAACATGATGTGCGCCGCGGACCCTCGCCACGGCCGCTATTTAACTGCCGCGTGTATGTTCCGCGGACGTATGAGCACGAAGGAGGTCGATGAGCAGATGCTCAACGTGCAGAACAAGAACTCGTCGTACTTCGTCGAGTGGATCCCCAACAACATCAAGGCCAGCGTGTGTGACATCCCGCCTCAGGGTCTCAAGATGAGCACCACGTTCATCGGCAACTCCACTGCCATCCAGGAGATGTTCAAGCGTGTTTCCGAACAGTTCACGGCGATGTTCCGTCGTAAGGCTTTCTTGCACTGGTACACGGGCGAGGGTATGGATGAGATGGAGTTCACGGAGGCTGAGTCCAACATGAACGATCTTGTGTCTGAGTACCAGCAGTACCAGGACGGCACCG","order":8},{"start":1,"name":"Ph\_agathacida/1-1288","end":1288,"id":"405122839","seq":"ATGAGAGAGCTCGTTCACATCCAGGGCGGCCAGTGCGGTAACCAGATCGGCGCCAAGTTCTGGGAAGTCATCTCTGACGAGCACGGCGTGGACCCCACGGGCTCGTACCACGGCGACTCGGACCTGCAGCTCGAGCGCATCAACGTGTACTACAACGAGGCCACGGGCGGCCGCTACGTGCCCCGCGCCATCCTCATGGATCTCGAGCCCGGCACCATGGACTCGGTCCGCGCCGGCCCCTACGGCCAGCTCTTCCGTCCGGACAACTTTGTGTTCGGCCAGACGGGCGCCGGTAACAACTGGGCCAAGGGACACTACACGGAGGGTGCCGAGCTTATCGACTCGGTGCTTGACGTCGTCCGTAAGGAGGCTGAGAGCTGTGATTGCCTGCAGGGTTTCCAGATCACCCACTCGCTTGGTGGCGGTACCGGTTCCGGTATGGGCACGCTTCTTATTTCCAAGATCCGCGAAGAGTACCCCGACCGTATCATGTGCACGTACTCGGTGTGCCCGTCGCCCAAGGTGTCGGACACGGTCGTGGAGCCTTACAACGCGACGCTGTCGGTGCACCAGCTTGTCGAGAACGCCGATGAGGTCATGTGCCTGGATAACGAGGCCCTGTACGACATTTGCTTCCGTACGCTCAAGCTCACCACCCCCACGTACGGTGACCTGAACCACTTGGTGTGCGCCGCCATGTCCGGTATCACGACGTGCCTGCGTTTCCCCGGTCAGCTGAACTCGGACCTGCGTAAGCTGGCCGTGAATCTGATTCCGTTCCCGCGTCTCCACTTCTTCATGATTGGTTTCGCCCCGCTGACGTCGCGTGGCTCGCAGCAGTACCGTGCCCTTACGGTGCCCGAGCTGACCCAGCAGCAATTCGATGCTAAGAACATGATGTGCGCCGCCGACCCTCGCCACGGCCGCTATTTAACTGCCGCGTGTATGTTCCGCGGACGTATGAGTACGAAGGAGGTTGATGAGCAGATGCTGAACGTGCAGAACAAGAACTCGTCGTACTTCGTTGAGTGGATCCCCAACAACATCAAGGCTAGCGTGTGTGACATCCCGCCCAAGGGTCTCAAGATGAGCACCACGTTCATTGGTAACTCGACTGCCATCCAGGAGATGTTCAAGCGTGTGTCCGAACAGTTCACTGCTATGTTCCGTCGTAAGGCTTTCTTGCACTGGTACACTGGTGAGGGTATGGATGAGATGGAGTTCACGGAGGCTGAGTCCAACATGAACGACCTGGTGTCTGAGTACCAGCAGTACCAGGACGCGACCG","order":9},{"start":1,"name":"Ph\_sojae/1-1288","end":1288,"id":"164530833","seq":"ATGAGAGAGCTCGTTCACATCCAGGGTGGCCAGTGCGGTAACCAGATCGGCGCCAAGTTCTGGGAGGTCATCTCCGACGAGCACGGCGTGGACCCCACGGGATCCTACCACGGCGACTCGGACCTGCAGCTGGAGCGCATCAACGTGTACTACAACGAGGCCACGGGCGGCCGCTACGTGCCGCGCGCCATCCTCATGGACCTGGAGCCCGGCACCATGGACTCGGTGCGCGCCGGCCCCTACGGCCAGCTCTTCCGCCCGGACAACTTCGTGTTCGGCCAGACGGGCGCCGGTAACAACTGGGCCAAGGGACACTACACGGAGGGTGCCGAGCTTATCGACTCGGTTCTCGACGTCGTCCGCAAGGAGGCTGAGAGCTGTGACTGCCTTCAGGGTTTCCAGATCACGCACTCGCTGGGTGGCGGTACCGGTTCCGGTATGGGTACGCTTCTTATCTCCAAGATTCGTGAGGAGTACCCGGACCGTATCATGTGCACGTACTCGGTCTGCCCGTCGCCTAAGGTGTCGGACACGGTCGTCGAGCCCTACAACGCTACGCTGTCCGTCCACCAGCTCGTTGAGAACGCCGATGAGGTCATGTGCCTGGATAACGAGGCCCTGTACGACATTTGCTTCCGTACCCTGAAGCTCACGACCCCCACCTACGGTGACCTGAACCACCTGGTGTGCGCCGCCATGTCCGGCATTACCACGTGCCTGCGTTTCCCCGGTCAGCTGAACTCGGACCTGCGTAAGCTTGCCGTGAACCTGATCCCGTTCCCGCGTCTCCACTTCTTCATGATCGGTTTCGCCCCGCTGACGTCGCGCGGCTCGCAGCAGTACCGTGCCCTGACGGTGCCCGAGCTGACCCAGCAGCAGTTCGATGCTAAGAACATGATGTGTGCCGCCGACCCTCGCCACGGCCGCTATTTAACTGCCGCGTGTATGTTCCGCGGACGTATGAGCACGAAGGAGGTTGACGAGCAGATGCTCAACGTGCAGAACAAGAACTCGTCGTACTTCGTCGAGTGGATCCCCAACAACATCAAGGCTAGCGTGTGTGACATCCCGCCCAAGGGTCTCAAGATGAGCACCACGTTCATCGGTAACTCGACCGCTATCCAGGAGATGTTCAAGCGCGTGTCCGAACAGTTCACGGCTATGTTCCGTCGTAAGGCTTTCTTGCACTGGTACACGGGTGAGGGTATGGACGAGATGGAGTTCACGGAGGCCGAGTCCAACATGAACGATCTTGTGTCTGAGTACCAGCAGTACCAGGACGCTACCG","order":10},{"start":1,"name":"Pe\_v\_pisi/1-1288","end":1288,"id":"909597647","seq":"ATGAGAGAGCTTGTTCACATTCAGGGTGGCCAGTGCGGAAACCAGATCGGAGCCAAGTTTTGGGAGGTTATCTCCGACGAGCATGGTGTGGATCCCACTGGCTCGTACCACGGCGACTCGGACCTGCAGCTTGAGCGTATCAATGTGTACTATAACGAAGCCACAGGTGGACGTTACGTACCCCGCGCTATCCTGATGGATCTGGAGCCCGGCACCATGGACTCGGTTCGCGCTGGCCCCTACGGTCAGCTTTTCCGCCCCGACAATTTTGTGTTTGGCCAGACAGGTGCCGGTAATAACTGGGCCAAAGGACACTACACTGAGGGTGCTGAGCTCATTGACTCAGTGCTGGACGTCGTCCGTAAAGAGGCGGAAAGCTGTGACTGTTTGCAGGGTTTCCAGATCACGCACTCGCTTGGTGGCGGGACCGGCTCCGGTATGGGAACGCTTCTTATCTCGAAGATCCGAGAAGAATACCCGGACCGTATCATGTGCACGTACTCGGTGTGTCCATCCCCCAAAGTGTCGGACACGGTTGTGGAGCCTTACAACGCTACGCTGTCTGTGCACCAGCTTGTCGAGAATGCCGATGAGGTCATGTGCCTCGATAACGAGGCCTTATACGACATTTGCTTTCGCACGCTGAAGCTCACCACCCCCACTTACGGTGACCTGAACCACTTGGTATGTGCCGCTATGTCTGGCATCACCACTTCCCTGCGATTCCCTGGACAGCTAAACTCGGACCTGCGGAAGCTGGCCGTGAACCTAATCCCTTTCCCCCGTCTCCATTTCTTCATGATTGGTTTTGCCCCGCTGACGTCACGTGGATCGCAGCAATACCGTGCCCTGACGGTGCCTGAGCTGACGCAGCAGCAGTTCGATGCTAAGAACATGATGTGCGCCGCTGACCCTCGTCACGGCCGCTATTTAACTGCCGCCTGTATGTTCCGCGGACGTATGAGCACGAAGGAGGTTGATGAGCAAATGCTGAACGTGCAGAACAAGAACTCGTCGTACTTTGTCGAGTGGATCCCCAACAATATCAAGGCTAGCGTGTGTGACATCCCGCCCAAGGGCCTGAAGATGAGCACTACCTTTATCGGTAATTCGACAGCTATCCAAGAGATGTTCAAGCGTGTGTCTGAACAGTTCACGGCTATGTTTCGTCGCAAGGCTTTCTTGCACTGGTACACTGGTGAGGGTATGGATGAGATGGAGTTCACGGAGGCTGAGTCCAACATGAACGATCTTGTGTCTGAGTACCAGCAATATCAGGATGCGACCG","order":11},{"start":1,"name":"Py\_aphanidermatum/1-1288","end":1288,"id":"290039182","seq":"ATGCGTGAGCTCGTTCACATTCAGGGTGGTCAGTGTGGTAACCAGATCGGTGCCAAGTTCTGGGAGGTGATCTCCGATGAGCACGGCGTGGACCCGACCGGTTCGTACCACGGTGACTCGGACCTTCAGCTTGAGCGCATCAACGTGTACTACAACGAAGCCACGGGTGGTCGCTATGTGCCTCGTGCCATTCTGATGGACTTGGAGCCTGGTACCATGGACTCGGTCCGCGCTGGTCCATACGGCCAGCTTTTCCGTCCGGACAACTTCGTGTTCGGCCAGACTGGTGCCGGTAACAACTGGGCCAAGGGTCACTACACGGAGGGTGCTGAGCTCATCGACTCGGTGCTTGACGTCGTCCGCAAGGAGGCTGAGAGCTGTGACTGCCTTCAGGGTTTCCAGATCACCCACTCGCTTGGTGGTGGTACCGGTTCTGGTATGGGTACCCTCCTCATCTCGAAGATTCGTGAAGAGTACCCAGACCGTATCATGTGCACGTACTCGGTGTGCCCATCGCCAAAGGTCTCGGATACCGTCGTTGAGCCATACAACGCCACGCTTTCGGTCCACCAGCTTGTCGAGAACGCCGATGAGGTGATGTGTCTTGACAACGAAGCCCTTTACGATATCTGTTTCCGTACCCTGAAGCTCACGACGCCAACCTACGGTGACCTGAACCACCTTGTGTGTGCTGCCATGTCGGGTATCACGACCAGTCTGCGTTTCCCAGGTCAGCTGAACTCGGATCTTCGTAAGCTTGCCGTCAACCTTATCCCGTTCCCACGTCTCCACTTCTTCATGGTCGGTTTCGCTCCGCTCACCTCGCGCGGCTCGCAGCAGTACCGTGCCCTTACGGTGCCAGAGCTGACCCAGCAGCAGTTCGACGCCAAGAACATGATGTGTGCCGCTGATCCTCGTCACGGTCGTTACCTGACCGCCGCTTGTATGTTCCGTGGTCGTATGAGCACCAAGGAGGTCGACGAGCAGATGCTCAACGTCCAGAACAAGAACTCGTCGTACTTCGTTGAGTGGATCCCGAACAACATCAAGGCCAGCGTTTGTGACATCCCGCCGAAGGGTCTCAAGATGAGCGCCACGTTCATCGGTAACTCGACTGCCATCCAGGAGATGTTCAAGCGTGTCAGCGAGCAGTTCACGGCCATGTTCCGTCGTAAGGCCTTCTTGCACTGGTACACTGGTGAGGGTATGGACGAGATGGAGTTCACTGAAGCCGAGTCGAACATGAACGATCTCGTGTCGGAGTACCAGCAGTACCAGGATGCTACGG","order":12},{"start":1,"name":"Pe\_effusa/1-1288","end":1288,"id":"614591911","seq":"ATGAGAGAGCTCGTTCATATTCAGGGTGGTCAGTGCGGAAACCAAATTGGAGCCAAGTTCTGGGAGGTTATTTCCGACGAGCATGGTGTAGATCCCACCGGCTCGTACCACGGTGACTCGGACCTGCAGCTCGAGCGCATTAATGTGTATTATAACGAGGCCACAGGCGGACGCTACGTGCCCCGTGCTATTCTTATGGACCTGGAGCCCGGCACCATGGACTCGGTTCGTGCTGGTCCCTACGGCCAGCTCTTCCGCCCCGACAATTTTGTGTTCGGCCAGACAGGTGCCGGTAACAATTGGGCTAAGGGACACTACACTGAGGGCGCCGAGCTCATTGACTCCGTTCTGGACGTCGTCCGTAAGGAGGCGGAAAGCTGTGACTGTTTGCAGGGTTTCCAGATCACGCACTCGCTTGGTGGCGGTACCGGCTCCGGTATGGGAACGCTTCTTATCTCAAAGATCCGAGAAGAGTACCCGGACCGTATCATGTGCACGTACTCGGTATGTCCATCTCCCAAGGTGTCGGACACTGTTGTAGAGCCCTACAATGCTACGCTGTCTGTACATCAGCTTGTCGAAAACGCCGATGAGGTCATGTGCCTGGATAATGAGGCCTTGTACGACATTTGCTTCCGTACGCTGAAGCTCACCACCCCCACCTATGGTGACCTGAACCACTTGGTGTGTGCCGCTATGTCGGGCATCACCACTTCCCTACGATTTCCGGGACAGCTGAACTCGGACCTGCGGAAGCTGGCCGTGAACCTGATCCCGTTCCCTCGTCTTCACTTCTTCATGATTGGTTTCGCGCCGCTGACGTCACGTGGTTCGCAGCAGTACCGTGCCTTGACGGTGCCTGAGCTGACACAGCAGCAGTTTGATGCTAAAAACATGATGTGCGCCGCCGACCCTCGCCACGGTCGCTATTTAACTGCCGCTTGTATGTTCCGCGGACGCATGAGCACGAAGGAGGTTGATGAGCAAATGCTGAACGTGCAGAACAAAAACTCGTCGTACTTTGTCGAGTGGATTCCTAACAACATCAAGGCTAGTGTGTGTGACATCCCACCCAAGGGCCTGAAGATGAGCACCACCTTTATTGGTAATTCGACAGCCATCCAAGAGATGTTCAAGCGTGTGTCTGAACAGTTCACGGCTATGTTCCGTCGTAAGGCTTTCTTGCATTGGTACACTGGTGAGGGTATGGACGAGATGGAGTTCACGGAGGCTGAGTCTAACATGAACGATCTCGTGTCTGAGTACCAGCAATACCAGGATGCGACCG","order":13},{"start":1,"name":"H\_arabidopsidis\_Emoy2/1-1288","end":1288,"id":"1497784736","seq":"ATGAGAGAACTCGTCCACATTCAGGGTGGACAGTGCGGAAACCAGATCGGTGCCAAGTTTTGGGAAGTCATTTCCGACGAGCACGGCGTGGACCCCACGGGCTCGTACCACGGCGACTCGGACCTGCAGCTGGAACGCATTAACGTGTACTACAACGAAGCAACGGGTGGACGCTATGTGCCTCGTGCTATTCTAATGGATCTCGAGCCCGGCACGATGGACTCGGTCCGCGCTGGTCCGTACGGCCAGCTCTTCCGTCCGGATAACTTTGTCTTTGGACAGACAGGAGCTGGTAACAATTGGGCCAAGGGACACTACACGGAGGGTGCAGAGCTTATTGACTCGGTGCTGGACGTTGTCCGCAAAGAGGCTGAGAGCTGTGACTGCCTACAAGGTTTCCAGTTCACGCACTCGCTCGGCGGTGGTACCGGCTCTGGTATGGGCACGCTTCTTATCTCAAAGATCCGAGAAGAGTATCCGGACCGTATTATGTGCACATACTCCGTGTGTCCGTCCCCCAAGGTGTCGGACACTGTCGTCGAGCCCTACAACGCTACGTTGTCGGTGCACCAGCTCGTGGAGAACGCTGATGAAGTCATGTGCCTTGACAATGAGGCCCTGTATGACATTTGCTTCCGCACGCTGAAGCTCACGACTCCTACTTACGGTGATCTGAATCACTTGGTGTGTGCTGCTATGTCTGGTATCACGACGTGCCTACGTTTTCCCGGTCAGCTGAACTCAGACCTCCGTAAGCTGGCTGTGAACTTGATTCCGTTTCCACGTCTTCACTTCTTCATGATTGGTTTCGCTCCGCTGACGTCGCGAAGCTCGCAGCAGTACCGTGCGTTGACGGTGCCCGAGCTGACGCAGCAGCAGTTCGATGCTAAAAACATGATGTGTGCCGCCGACCCCCGCCATGGCCGCTATTTAACTGCCGCGTGTATGTTCCGCGGACGTATGAGTACGAAGGAGGTGGATGAGCAAATGCTGAACGTGCAAAACAAGAACTCGTCGTACTTCGTCGAGTGGATTCCCAACAACATCAAGGCTAGCGTGTGTGACATCCCGCCCAAGGGGTTGAAGATGAGTACCACCTTCATCGGTAACTCGACCGCAATCCAGGAGATGTTCAAGCGCGTGTCTGAGCAATTTACGGCCATGTTTCGTCGTAAGGCTTTCTTGCACTGGTACACTGGTGAGGGCATGGATGAGATGGAGTTCACGGAGGCCGAGTCCAACATGAACGACCTCGTGTCCGAGTACCAGCAGTACCAGGATGCGACCG","order":14},{"start":1,"name":"H\_arabidopsidis\_Noks1/1-1288","end":1288,"id":"1436790884","seq":"ATGAGAGAACTCGTCCACATTCAGGGTGGACAGTGCGGAAACCAGATCGGTGCCAAGTTTTGGGAAGTCATTTCCGACGAGCACGGCGTGGACCCCACGGGCTCGTACCACGGCGACTCGGACCTGCAGCTGGAACGCATTAACGTGTACTACAACGAAGCAACGGGTGGACGCTATGTGCCTCGTGCTATTCTAATGGATCTCGAGCCCGGCACGATGGACTCGGTCCGCGCTGGTCCGTACGGCCAGCTCTTCCGTCCGGATAACTTTGTCTTTGGACAGACAGGAGCTGGTAACAATTGGGCCAAGGGACACTACACGGAGGGTGCAGAGCTTATTGACTCGGTGCTGGACGTTGTCCGCAAAGAGGCTGAGAGCTGTGACTGCCTACAAGGTTTCCAGTTCACGCACTCGCTCGGCGGTGGTACCGGCTCTGGTATGGGCACGCTTCTTATCTCAAAGATCCGAGAAGAGTATCCGGACCGTATTATGTGCACATACTCCGTGTGTCCGTCCCCCAAGGTGTCGGACACTGTCGTCGAGCCCTACAACGCTACGTTGTCGGTGCACCAGCTCGTGGAGAACGCTGATGAAGTCATGTGCCTTGACAATGAGGCCCTGTATGACATTTGCTTCCGCACGCTGAAGCTCACGACTCCTACTTACGGTGATCTGAATCACTTGGTGTGTGCTGCTATGTCTGGTATCACGACGTGCCTACGTTTTCCCGGTCAGCTGAACTCAGACCTCCGTAAGCTGGCTGTGAACTTGATTCCGTTTCCACGTCTTCACTTCTTCATGATTGGTTTCGCTCCGTTGACGTCGCGAAGCTCGCAGCAGTACCGTGCGTTGACGGTGCCCGAGCTGACGCAGCAGCAGTTCGATGCTAAAAACATGATGTGTGCCGCCGACCCCCGCCATGGCCGCTATTTAACTGCCGCGTGTATGTTCCGCGGACGTATGAGTACGAAGGAGGTGGATGAGCAAATGCTGAACGTGCAAAACAAGAACTCGTCGTACTTCGTCGAGTGGATTCCCAACAACATCAAGGCTAGCGTGTGTGACATCCCGCCCAAGGGGTTGAAGATGAGTACCACCTTCATCGGTAACTCGACCGCAATCCAGGAGATGTTCAAGCGCGTGTCTGAGCAATTTACGGCCATGTTTCGTCGTAAGGCTTTCTTGCACTGGTACACTGGTGAGGGCATGGATGAGATGGAGTTCACGGAGGCCGAGTCCAACATGAACGACCTCGTGTCCGAGTACCAGCAGTACCAGGATGCGACCG","order":15},{"start":1,"name":"H\_arabidopsidis\_Cala2/1-1288","end":1288,"id":"161094283","seq":"ATGAGAGAACTCGTCCACATTCAGGGTGGACAGTGCGGAAACCAGATCGGTGCCAAGTTTTGGGAAGTCATTTCCGACGAGCACGGCGTGGACCCCACGGGCTCGTACCACGGCGACTCGGACCTGCAGCTGGAACGCATTAACGTGTACTACAACGAAGCAACGGGTGGACGCTATGTGCCTCGTGCTATTCTAATGGATCTCGAGCCCGGCACGATGGACTCGGTCCGCGCTGGTCCGTACGGCCAGCTCTTCCGTCCGGATAACTTTGTCTTTGGACAGACAGGAGCTGGTAACAATTGGGCCAAGGGACACTACACGGAGGGTGCAGAGCTTATTGACTCGGTGCTGGACGTTGTCCGCAAAGAGGCTGAGAGCTGTGACTGCCTACAAGGTTTCCAGTTCACGCACTCGCTCGGCGGTGGTACCGGCTCTGGTATGGGCACGCTTCTTATCTCAAAGATCCGAGAAGAGTATCCGGACCGTATTATGTGCACATACTCCGTGTGTCCGTCCCCCAAGGTGTCGGACACTGTCGTCGAGCCCTACAACGCTACGTTGTCGGTGCACCAGCTCGTGGAGAACGCTGATGAAGTCATGTGCCTTGACAATGAGGCCCTGTATGACATTTGCTTCCGCACGCTGAAGCTCACGACTCCTACTTACGGTGATCTGAATCACTTGGTGTGTGCTGCTATGTCTGGTATCACGACGTGCCTACGTTTTCCCGGTCAGCTGAACTCAGACCTCCGTAAGCTGGCTGTGAACTTGATTCCGTTTCCACGTCTTCACTTCTTCATGATTGGTTTCGCTCCGTTGACGTCGCGAAGCTCGCAGCAGTACCGTGCGTTGACGGTGCCCGAGCTGACGCAGCAGCAGTTCGATGCTAAAAACATGATGTGTGCCGCCGACCCCCGCCATGGCCGCTATTTAACTGCCGCGTGTATGTTCCGCGGACGTATGAGTACGAAGGAGGTGGATGAGCAAATGCTGAACGTGCAAAACAAGAACTCGTCGTACTTCGTCGAGTGGATTCCCAACAACATCAAGGCTAGCGTGTGTGACATCCCGCCCAAGGGGTTGAAGATGAGTACCACCTTCATCGGTAACTCGACCGCAATCCAGGAGATGTTCAAGCGCGTGTCTGAGCAATTTACGGCCATGTTTCGTCGTAAGGCTTTCTTGCACTGGTACACTGGTGAGGGCATGGATGAGATGGAGTTCACGGAGGCCGAGTCCAACATGAACGACCTCGTGTCCGAGTACCAGCAGTACCAGGATGCGACCG","order":16},{"start":1,"name":"Ph\_infestans/1-1288","end":1288,"id":"587010762","seq":"ATGAGAGAGCTCGTTCACATTCAGGGTGGTCAGTGTGGTAACCAGATCGGTGCCAAGTTCTGGGAGGTCATCTCCGACGAGCACGGCGTGGACCCCACGGGCTCGTACCACGGCGACTCAGACCTGCAACTGGAGCGCATTAATGTGTACTACAACGAGGCCACGGGCGGCCGTTACGTGCCCCGCGCCATCCTCATGGATCTTGAGCCCGGTACCATGGACTCCGTTCGCGCTGGCCCCTACGGTCAGCTTTTCCGCCCAGACAATTTCGTGTTCGGACAGACTGGCGCTGGTAACAACTGGGCCAAGGGACACTACACTGAGGGCGCTGAGCTGATTGACTCGGTGCTTGACGTCGTTCGCAAGGAGGCAGAGAGCTGTGATTGCCTTCAGGGTTTCCAGATCACGCACTCGCTTGGTGGCGGTACCGGTTCCGGTATGGGTACGCTTCTTATCTCGAAGATTCGTGAGGAGTACCCCGATCGCATCATGTGCACATACTCGGTCTGCCCGTCGCCCAAGGTATCGGACACGGTCGTGGAGCCCTATAACGCTACGCTATCGGTACACCAGCTTGTCGAGAACGCCGATGAGGTCATGTGCCTGGACAATGAGGCCCTGTACGACATTTGCTTCCGCACATTGAAGCTCACCACCCCCACTTATGGTGACCTGAACCACTTGGTGTGTGCCGCCATGTCCGGTATTACCACGTGCCTTCGTTTCCCCGGTCAGCTGAACTCGGACCTGCGTAAGCTGGCCGTGAACCTGATCCCGTTCCCGCGTCTCCACTTCTTTATGATTGGTTTCGCTCCTCTGACATCGCGCGGCTCGCAGCAGTACCGCGCCCTGACGGTGCCCGAGCTGACCCAGCAGCAGTTCGATGCTAAGAACATGATGTGTGCCGCCGACCCTCGCCATGGCCGCTATTTAACTGCAGCTTGTATGTTCCGCGGACGCATGAGCACGAAGGAGGTTGATGAGCAGATGCTGAACGTGCAGAACAAGAACTCGTCATACTTCGTCGAGTGGATCCCCAACAACATCAAGGCTAGCGTGTGTGACATCCCGCCCAAGGGTCTGAAGATGAGCACTACGTTCATTGGTAACTCTACTGCTATCCAAGAGATGTTCAAGCGTGTGTCCGAACAGTTTACGGCTATGTTCCGTCGTAAGGCTTTCTTGCACTGGTACACCGGTGAGGGTATGGACGAGATGGAGTTCACTGAGGCTGAGTCCAACATGAACGATCTGGTGTCTGAGTACCAGCAGTACCAGGACGCCACTG","order":17},{"start":1,"name":"Ph\_plurivora/1-1288","end":1288,"id":"1214004308","seq":"ATGAGAGAGCTTGTTCACATCCAGGGTGGTCAGTGCGGTAACCAGATCGGTGCCAAGTTCTGGGAGGTCATCTCCGACGAGCACGGCGTGGACCCCACGGGATCGTACCACGGCGACTCGGACCTACAGCTGGAGCGCATCAATGTGTACTACAACGAAGCTACGGGCGGCCGTTACGTGCCCCGCGCCATCCTCATGGACCTGGAGCCCGGCACCATGGACTCGGTCCGTGCTGGCCCTTACGGCCAGCTCTTCCGTCCGGACAACTTCGTGTTCGGCCAAACAGGCGCTGGTAACAACTGGGCCAAGGGACACTACACAGAGGGTGCCGAGCTTATCGACTCGGTGCTTGATGTCGTCCGCAAGGAGGCTGAGAGCTGTGACTGCCTGCAGGGTTTCCAGATCACCCACTCGCTTGGTGGCGGTACCGGTTCCGGTATGGGTACGCTTCTTATCTCGAAGATTCGTGAGGAGTACCCAGACCGTATTATGTGCACGTACTCGGTCTGCCCGTCGCCCAAGGTGTCGGACACCGTCGTGGAGCCCTACAACGCCACGTTGTCGGTGCATCAGCTGGTCGAGAACGCCGATGAGGTCATGTGCCTGGATAACGAGGCCCTGTACGATATTTGCTTCCGTACGCTGAAGCTCACCACGCCCACCTACGGTGACCTGAACCACCTGGTGTGCGCCGCCATGTCCGGTATCACCACGTGCCTGCGTTTCCCCGGTCAGTTGAACTCGGACCTGCGTAAGCTGGCCGTGAACCTGATCCCGTTCCCGCGTCTCCACTTCTTTATGATTGGTTTCGCTCCGCTGACCTCGCGTGGATCTCAGCAGTACCGTGCTCTTACGGTGCCGGAGCTGACCCAGCAGCAGTTCGATGCCAAGAACATGATGTGCGCCGCTGACCCTCGCCACGGCCGCTATTTAACTGCCGCGTGTATGTTCCGCGGACGTATGAGCACGAAGGAGGTTGACGAGCAGATGCTGAACGTGCAGAACAAGAACTCATCGTACTTCGTCGAGTGGATCCCCAACAACATCAAGGCTAGCGTGTGTGATATTCCTCCCAAGGGTCTGAAGATGAGCACTACGTTCATCGGTAACTCGACTGCCATCCAGGAGATGTTCAAGCGTGTGTCTGAACAGTTTACGGCTATGTTCCGTCGTAAGGCTTTCTTGCACTGGTACACTGGTGAGGGTATGGATGAGATGGAGTTCACTGAGGCCGAGTCCAACATGAACGATCTGGTGTCTGAGTACCAGCAGTACCAGGACGCCACCG","order":18},{"start":1,"name":"Br\_lactucae/1-1288","end":1288,"id":"11834795","seq":"ATGAGAGAACTTGTTCACATCCAGGGTGGTCAATGCGGTAACCAAATTGGAGCTAAGTTCTGGGAAGTCATCTCTGACGAGCACGGCGTAGACCCGACTGGTTCGTATCACGGCGACTCAGACCTTCAACTTGAGCGCATCAACGTTTATTACAATGAGGCCACGGGTGGACGCTATGTGCCTCGCGCCATCCTAATGGATCTTGAACCTGGAACTATGGACTCAGTCCGCGCTGGTCCTTATGGCCAACTCTTCCGCCCAGACAATTTTGTGTTCGGCCAGACAGGCGCCGGTAACAATTGGGCGAAAGGACATTATACCGAGGGTGCCGAGCTTATTGATTCGGTGCTTGACGTCGTTCGCAAAGAGGCAGAGAGCTGTGATTGCCTTCAGGGTTTCCAAATCACGCACTCGCTTGGTGGCGGTACTGGTTCTGGGATGGGAACGCTTCTTATTTCAAAGATTCGTGAGGAATACCCCGACCGTATCATGTGCACATACTCCGTGTGCCCTTCGCCAAAGGTATCGGATACGGTCGTGGAGCCTTACAATGCTACGCTTTCCGTGCACCAGCTCGTTGAAAACGCTGATGAAGTCATGTGCTTGGACAATGAGGCCCTGTACGATATTTGCTTTCGAACGTTGAAGCTTACCACCCCAACTTATGGTGATTTGAACCACCTGGTGTGCGCCGCCATGTCTGGTATAACTACATGCCTGCGTTTTCCAGGCCAACTGAATTCTGATCTTCGTAAATTGGCTGTAAACCTTATTCCATTCCCGCGTCTTCACTTCTTTATGATTGGCTTTGCTCCGTTGACGTCGCGTGGCTCGCAGCAATACCGTGCCCTGACGGTTCCTGAATTAACTCAACAGCAGTTTGACGCGAAGAATATGATGTGTGCCGCTGACCCTCGTCATGGTCGCTATTTAACTGCCGCATGTATGTTTCGTGGACGCATGAGCACCAAGGAGGTTGACGAGCAGATGCTTAACGTGCAGAATAAGAATTCCTCCTACTTTGTCGAGTGGATTCCCAACAACATCAAAGCTAGTGTATGTGACATTCCGCCGAAGGGTTTGAAAATGAGCACCACATTTATTGGTAATTCAACAGCTATCCAAGAGATGTTCAAGCGAGTGTCAGAACAGTTCACGGCCATGTTCCGTCGTAAGGCTTTCTTGCACTGGTACACCGGCGAGGGCATGGACGAGATGGAGTTTACTGAAGCCGAGTCGAACATGAACGATTTGGTTTCTGAGTACCAGCAGTACCAAGACGCGACTG","order":19},{"start":1,"name":"Pl\_halstedii/1-1288","end":1288,"id":"1503893876","seq":"ATGAGAGAGCTTGTTCATATCCAGGGAGGTCAGTGTGGAAACCAGATCGGCGCAAAGTTCTGGGAAGTTATCTCTGATGAGCACGGAGTGGATCCCACTGGTTCGTACCACGGCGATTCGGATCTTCAATTAGAGCGCATCAATGTGTACTATAATGAAGCTACGGGTGGACGTTACGTTCCACGCGCCATTTTGATGGACCTGGAGCCTGGCACAATGGACTCGGTCCGCGCCGGCCCATATGGTCAGCTTTTCCGCCCAGACAATTTTGTTTTTGGCCAGACAGGCGCCGGTAACAATTGGGCGAAAGGACATTATACCGAGGGTGCCGAGCTGATTGATTCCGTGCTAGACGTTGTCCGCAAGGAAGCAGAAAGTTGCGATTGTCTTCAAGGTTTTCAAATTACGCACTCACTTGGTGGTGGTACCGGCTCCGGAATGGGCACACTTCTTATTTCGAAAATTCGGGAAGAGTATCCCGACCGAATCATGTGCACCTACTCGGTGTGCCCATCGCCTAAGGTATCAGATACGGTAGTAGAGCCTTACAATGCCACATTGTCGGTGCACCAGCTAGTCGAGAACGCTGATGAAGTTATGTGCTTAGACAACGAAGCTTTGTACGACATTTGCTTTCGTACGTTAAAACTCACCACGCCTACTTACGGTGACTTGAATCACTTGGTGTGTGCTGCCATGTCTGGTATTACCACATGCTTGCGATTTCCTGGTCAGCTAAATTCAGACTTGCGGAAGTTGGCCGTCAATCTGATTCCATTTCCCCGTCTTCATTTTTTTATGATTGGTTTTGCACCTTTGACATCACGTGGCTCGCAGCAGTATCGTGCGTTGACAGTACCTGAGTTGACCCAGCAGCAGTTTGATGCTAAAAACATGATGTGCGCCGCTGACCCTCGCCACGGTCGCTATTTAACTGCCGCGTGTATGTTTCGAGGGCGCATGAGCACGAAGGAAGTTGATGAGCAGATGCTGAATGTGCAAAACAAAAATTCTTCTTATTTCGTCGAGTGGATTCCCAACAACATCAAGGCGAGTGTGTGTGACATCCCGCCTAAGGGTTTAAAAATGAGCACCACGTTTATTGGTAATTCGACTGCTATTCAGGAGATGTTTAAGCGAGTGTCAGAACAGTTCACGGCAATGTTCCGTCGTAAGGCTTTCTTGCATTGGTACACTGGTGAGGGAATGGACGAAATGGAGTTTACTGAGGCTGAGTCCAACATGAACGATTTAGTTTCTGAATACCAACAGTACCAAGATGCCACCG","order":20}],"appSettings":{"globalColorScheme":"Nucleotide","webStartUrl":"https://www.jalview.org/services/launchApp","application":"Jalview","showSeqFeatures":"true","version":"2.11.4.1"},"seqGroups":[],"alignAnnotation":[],"svid":"1.0","seqFeatures":[{"fillColor":"#0000ff","score":0,"sequenceRef":"1497784736","featureGroup":"Jalview","description":"","xStart":530,"xEnd":621,"type":"BTUB E. coli-produced dsRNA"},{"fillColor":"#0000ff","score":0,"sequenceRef":"1497784736","featureGroup":"Jalview","description":"","xStart":531,"xEnd":804,"type":"BTUB E. coli-produced dsRNA"},{"fillColor":"#ff3333","score":0,"sequenceRef":"1497784736","featureGroup":"Jalview","description":"","xStart":680,"xEnd":710,"type":"Hpa-BTUB-SS-dsRNA"},{"fillColor":"#0000ff","score":0,"sequenceRef":"1436790884","featureGroup":"Jalview","description":"","xStart":530,"xEnd":621,"type":"BTUB E. coli-produced dsRNA"},{"fillColor":"#0000ff","score":0,"sequenceRef":"1436790884","featureGroup":"Jalview","description":"","xStart":531,"xEnd":804,"type":"BTUB E. coli-produced dsRNA"},{"fillColor":"#ff3333","score":0,"sequenceRef":"1436790884","featureGroup":"Jalview","description":"","xStart":680,"xEnd":710,"type":"Hpa-BTUB-SS-dsRNA"},{"fillColor":"#0000ff","score":0,"sequenceRef":"161094283","featureGroup":"Jalview","description":"","xStart":530,"xEnd":621,"type":"BTUB E. coli-produced dsRNA"},{"fillColor":"#0000ff","score":0,"sequenceRef":"161094283","featureGroup":"Jalview","description":"","xStart":531,"xEnd":804,"type":"BTUB E. coli-produced dsRNA"},{"fillColor":"#ff3333","score":0,"sequenceRef":"161094283","featureGroup":"Jalview","description":"","xStart":680,"xEnd":710,"type":"Hpa-BTUB-SS-dsRNA"}]}

  

xml version="1.0"?

ZebrafishArabidopsisHumanWormYeastHy\_brassicaePh\_ramorumPh\_cinnamomiPh\_agathacidaPh\_sojaePe\_v\_pisiPy\_aphanidermatumPe\_effusaH\_arabidopsidis\_Emoy2H\_arabidopsidis\_Noks1H\_arabidopsidis\_Cala2Ph\_infestansPh\_plurivoraBr\_lactucaePl\_halstediiConsensusZebrafishArabidopsisHumanWormYeastHy\_brassicaePh\_ramorumPh\_cinnamomiPh\_agathacidaPh\_sojaePe\_v\_pisiPy\_aphanidermatumPe\_effusaH\_arabidopsidis\_Emoy2H\_arabidopsidis\_Noks1H\_arabidopsidis\_Cala2Ph\_infestansPh\_plurivoraBr\_lactucaePl\_halstediiConsensusZebrafishArabidopsisHumanWormYeastHy\_brassicaePh\_ramorumPh\_cinnamomiPh\_agathacidaPh\_sojaePe\_v\_pisiPy\_aphanidermatumPe\_effusaH\_arabidopsidis\_Emoy2H\_arabidopsidis\_Noks1H\_arabidopsidis\_Cala2Ph\_infestansPh\_plurivoraBr\_lactucaePl\_halstediiConsensusZebrafishArabidopsisHumanWormYeastHy\_brassicaePh\_ramorumPh\_cinnamomiPh\_agathacidaPh\_sojaePe\_v\_pisiPy\_aphanidermatumPe\_effusaH\_arabidopsidis\_Emoy2H\_arabidopsidis\_Noks1H\_arabidopsidis\_Cala2Ph\_infestansPh\_plurivoraBr\_lactucaePl\_halstediiConsensusZebrafishArabidopsisHumanWormYeastHy\_brassicaePh\_ramorumPh\_cinnamomiPh\_agathacidaPh\_sojaePe\_v\_pisiPy\_aphanidermatumPe\_effusaH\_arabidopsidis\_Emoy2H\_arabidopsidis\_Noks1H\_arabidopsidis\_Cala2Ph\_infestansPh\_plurivoraBr\_lactucaePl\_halstediiConsensusZebrafishArabidopsisHumanWormYeastHy\_brassicaePh\_ramorumPh\_cinnamomiPh\_agathacidaPh\_sojaePe\_v\_pisiPy\_aphanidermatumPe\_effusaH\_arabidopsidis\_Emoy2H\_arabidopsidis\_Noks1H\_arabidopsidis\_Cala2Ph\_infestansPh\_plurivoraBr\_lactucaePl\_halstediiConsensusZebrafishArabidopsisHumanWormYeastHy\_brassicaePh\_ramorumPh\_cinnamomiPh\_agathacidaPh\_sojaePe\_v\_pisiPy\_aphanidermatumPe\_effusaH\_arabidopsidis\_Emoy2H\_arabidopsidis\_Noks1H\_arabidopsidis\_Cala2Ph\_infestansPh\_plurivoraBr\_lactucaePl\_halstediiConsensusATGAGGGAGATCGTGCATTTACAGGCTGGACAGTGCGGCAACCAGATTGGTGCCAAGTTCTGGGAAGTCATTAGTGACGAGCATGGAATTGACCCAACAGGCAGTTACCATGGCGACAGTGACCTTCAGCTGGACCGAATTAATGTTTATTATAATGAAGCCACAGGTGGAAAGTATGTTCCACGTGCCGTGCTGGTGGATTTGGAGCCATGAGAGAGATCCTTCATATCCAAGGCGGTCAATGTGGAAACCAGATCGGAGCAAAGTTCTGGGAAGTGATCTGCGACGAACACGGCATTGATCACACCGGTCAATACGTCGGCGATTCTCCGTTACAGCTTGAACGTATCGATGTCTATTTCAACGAAGCTAGCGGTGGAAAGTACGTTCCTCGCGCTGTTCTTATGGATCTGGAGCCATGAGGGAAATCGTGCACATCCAGGCTGGTCAGTGTGGCAACCAGATCGGTGCCAAGTTCTGGGAGGTGATCAGTGATGAACATGGCATCGACCCCACCGGCACCTACCACGGGGACAGCGACCTGCAGCTGGACCGCATCTCTGTGTACTACAATGAAGCCACAGGTGGCAAATATGTTCCTCGTGCCATCCTGGTGGATCTAGAACCATGCGTGAAATTGTTCATATCCAGGCAGGTCAGTGTGGTAACCAAATTGGGGCAAAGTTCTGGGAAGTTATTTCCGACGAGCACGGGATCGATCCCACCGGAGCATACAATGGAGACTCCGATTTGCAGTTGGAGAGAATCAATGTCTACTACAACGAAGCTAGCGGAGGAAAGTATGTCCCACGTGCTTGTCTTGTTGATTTGGAGCCATGAGAGAAATCATTCATATCTCGACAGGTCAGTGTGGTAACCAAATTGGTGCTGCATTCTGGGAAACTATCTGTGGTGAGCACGGTTTGGATTTCAATGGGACATATCACGGCCATGACGATATCCAGAAGGAGAGACTGAACGTGTACTTCAACGAGGCATCTTCTGGGAAGTGGGTTCCAAGATCTATTAACGTCGATCTAGAACCATGCGCGAGCTCGTCCACATTCAGGGCGGCCAGTGCGGCAACCAGATCGGCGCCAAGTTCTGGGAAGTCATTTCCGACGAGCACGGCGTGGACCCCACGGGCTCGTACCGCGGCGACTCGGACCTGCAGCTCGAGCGCATCAACGTGTACTACAACGAAGCGACGGGCGGACGCTATGTGCCCCGCGCGATTCTGATGGACCTGGAGCCATGAGAGAGCTCGTTCACATCCAGGGTGGCCAGTGCGGTAACCAGATCGGCGCCAAGTTCTGGGAGGTTATCTCCGACGAGCACGGCGTGGACCCCACGGGCTCGTACCACGGCGACTCGGACCTGCAGCTGGAGCGCATCAATGTGTACTACAACGAGGCCACGGGCGGCCGCTACGTGCCCCGCGCCATCCTCATGGACCTGGAGCCATGAGAGAGCTCGTTCACATCCAGGGTGGCCAGTGCGGTAACCAGATCGGCGCCAAGTTCTGGGAGGTCATCTCCGACGAGCACGGCGTGGACCCGACGGGATCCTACCACGGCGACTCGGACCTGCAGCTGGAGCGCATCAACGTGTACTACAACGAGGCCACGGGCGGCCGCTACGTGCCCCGCGCCATCCTCATGGACCTGGAGCCATGAGAGAGCTCGTTCACATCCAGGGCGGCCAGTGCGGTAACCAGATCGGCGCCAAGTTCTGGGAAGTCATCTCTGACGAGCACGGCGTGGACCCCACGGGCTCGTACCACGGCGACTCGGACCTGCAGCTCGAGCGCATCAACGTGTACTACAACGAGGCCACGGGCGGCCGCTACGTGCCCCGCGCCATCCTCATGGATCTCGAGCCATGAGAGAGCTCGTTCACATCCAGGGTGGCCAGTGCGGTAACCAGATCGGCGCCAAGTTCTGGGAGGTCATCTCCGACGAGCACGGCGTGGACCCCACGGGATCCTACCACGGCGACTCGGACCTGCAGCTGGAGCGCATCAACGTGTACTACAACGAGGCCACGGGCGGCCGCTACGTGCCGCGCGCCATCCTCATGGACCTGGAGCCATGAGAGAGCTTGTTCACATTCAGGGTGGCCAGTGCGGAAACCAGATCGGAGCCAAGTTTTGGGAGGTTATCTCCGACGAGCATGGTGTGGATCCCACTGGCTCGTACCACGGCGACTCGGACCTGCAGCTTGAGCGTATCAATGTGTACTATAACGAAGCCACAGGTGGACGTTACGTACCCCGCGCTATCCTGATGGATCTGGAGCCATGCGTGAGCTCGTTCACATTCAGGGTGGTCAGTGTGGTAACCAGATCGGTGCCAAGTTCTGGGAGGTGATCTCCGATGAGCACGGCGTGGACCCGACCGGTTCGTACCACGGTGACTCGGACCTTCAGCTTGAGCGCATCAACGTGTACTACAACGAAGCCACGGGTGGTCGCTATGTGCCTCGTGCCATTCTGATGGACTTGGAGCCATGAGAGAGCTCGTTCATATTCAGGGTGGTCAGTGCGGAAACCAAATTGGAGCCAAGTTCTGGGAGGTTATTTCCGACGAGCATGGTGTAGATCCCACCGGCTCGTACCACGGTGACTCGGACCTGCAGCTCGAGCGCATTAATGTGTATTATAACGAGGCCACAGGCGGACGCTACGTGCCCCGTGCTATTCTTATGGACCTGGAGCCATGAGAGAACTCGTCCACATTCAGGGTGGACAGTGCGGAAACCAGATCGGTGCCAAGTTTTGGGAAGTCATTTCCGACGAGCACGGCGTGGACCCCACGGGCTCGTACCACGGCGACTCGGACCTGCAGCTGGAACGCATTAACGTGTACTACAACGAAGCAACGGGTGGACGCTATGTGCCTCGTGCTATTCTAATGGATCTCGAGCCATGAGAGAACTCGTCCACATTCAGGGTGGACAGTGCGGAAACCAGATCGGTGCCAAGTTTTGGGAAGTCATTTCCGACGAGCACGGCGTGGACCCCACGGGCTCGTACCACGGCGACTCGGACCTGCAGCTGGAACGCATTAACGTGTACTACAACGAAGCAACGGGTGGACGCTATGTGCCTCGTGCTATTCTAATGGATCTCGAGCCATGAGAGAACTCGTCCACATTCAGGGTGGACAGTGCGGAAACCAGATCGGTGCCAAGTTTTGGGAAGTCATTTCCGACGAGCACGGCGTGGACCCCACGGGCTCGTACCACGGCGACTCGGACCTGCAGCTGGAACGCATTAACGTGTACTACAACGAAGCAACGGGTGGACGCTATGTGCCTCGTGCTATTCTAATGGATCTCGAGCCATGAGAGAGCTCGTTCACATTCAGGGTGGTCAGTGTGGTAACCAGATCGGTGCCAAGTTCTGGGAGGTCATCTCCGACGAGCACGGCGTGGACCCCACGGGCTCGTACCACGGCGACTCAGACCTGCAACTGGAGCGCATTAATGTGTACTACAACGAGGCCACGGGCGGCCGTTACGTGCCCCGCGCCATCCTCATGGATCTTGAGCCATGAGAGAGCTTGTTCACATCCAGGGTGGTCAGTGCGGTAACCAGATCGGTGCCAAGTTCTGGGAGGTCATCTCCGACGAGCACGGCGTGGACCCCACGGGATCGTACCACGGCGACTCGGACCTACAGCTGGAGCGCATCAATGTGTACTACAACGAAGCTACGGGCGGCCGTTACGTGCCCCGCGCCATCCTCATGGACCTGGAGCCATGAGAGAACTTGTTCACATCCAGGGTGGTCAATGCGGTAACCAAATTGGAGCTAAGTTCTGGGAAGTCATCTCTGACGAGCACGGCGTAGACCCGACTGGTTCGTATCACGGCGACTCAGACCTTCAACTTGAGCGCATCAACGTTTATTACAATGAGGCCACGGGTGGACGCTATGTGCCTCGCGCCATCCTAATGGATCTTGAACCATGAGAGAGCTTGTTCATATCCAGGGAGGTCAGTGTGGAAACCAGATCGGCGCAAAGTTCTGGGAAGTTATCTCTGATGAGCACGGAGTGGATCCCACTGGTTCGTACCACGGCGATTCGGATCTTCAATTAGAGCGCATCAATGTGTACTATAATGAAGCTACGGGTGGACGTTACGTTCCACGCGCCATTTTGATGGACCTGGAGCCATGAGAGAGCTCGTTCACATCCAGGGTGGTCAGTGCGGTAACCAGATCGGTGCCAAGTTCTGGGAAGTCATCTCCGACGAGCACGGCGTGGACCCCACGGGCTCGTACCACGGCGACTCGGACCTGCAGCTGGAGCGCATCAA+GTGTACTACAACGAAGCCACGGGTGGACGCTACGTGCCCCGCGCCATTCTCATGGATCTGGAGCCTGGTACAATGGACTCCGTGAGGTCTGGTCCATTTGGTCAGATCTTCAGACCAGACAACTTTGTGTTCGGCCAGAGTGGTGCTGGAAACAACTGGGCCAAGGGCCACTACACTGAAGGAGCTGAGCTGGTTGATTCCGTTCTGGATGTGGTCCGAAAAGAGGCTGAGAGCTGCGACTGTCTGCAGGGCTTCCAACTCACTCACTCACTGGTGGTACCATGGATTCTCTCAGATCTGGTCCGTTCGGTCAGATTTTCCGTCCTGATAACTTCGTCTTTGGTCAATCTGGTGCCGGAAATAACTGGGCGAAAGGTCATTACACCGAAGGTGCTGAGTTGATTGATTCTGTTCTCGATGTTGTGAGGAAGGAAGCTGAGAACAGCGATTGTCTTCAAGGTTTCCAAGTGTGTCATTCATTGGTGGGACCATGGACTCTGTTCGCTCAGGTCCTTTTGGCCAGATCTTTAGACCAGACAACTTTGTATTTGGTCAGTCTGGGGCAGGTAACAACTGGGCCAAAGGCCACTACACAGAGGGCGCCGAGCTGGTTGATTCTGTCCTGGATGTGGTACGGAAGGAGGCAGAGAGCTGTGACTGCCTGCAGGGCTTCCAGCTGACCCACTCACTGGAGGAACCATGGACTCCGTTAGGGCAGGTCCCTTTGGACAGCTCTTTCGACCAGACAACTTTGTTTTTGGTCAAAGCGGTGCCGGAAACAACTGGGCCAAAGGTCACTACACCGAGGGCGCCGAACTTGTAGACAATGTTCTCGACGTTGTCCGTAAAGAAGCTGAAAGCTGTGATTGTCTTCAGGGATTCCAAATGACTCACTCTTTGGTGGGACGATTGACGCAGTACGCAATTCTGCCATCGGGAATTTGTTTAGACCTGACAATTATATCTTTGGGCAAAGTTCTGCGGGCAACGTGTGGGCCAAGGGTCACTACACAGAAGGTGCTGAGCTTGTAGACAGCGTCATGGATGTTATTAGACGAGAGGCCGAAGGATGCGACTCCCTTCAAGGTTTCCAGATCACACATTCTCTTGCGGCACGATGGACTCTGTCCGCGCAGGTCCGTACGGCCAGCTCTTCCGTCCGGACAACTTTGTGTTTGGACAGACAGGCGCGGGTAACAACTGGGCCAAGGGACACTATACGGAGGGTGCGGAGCTCATTGATTCGGTGCTGGACGTTGTCCGTAAAGAGGCCGAGAGCTGTGACTGCCTACAGGGTTTCCAGTTCACGCACTCGCTTGCGGCACCATGGACTCGGTCCGCGCCGGCCCCTACGGCCAGCTCTTCCGCCCGGACAACTTCGTGTTCGGTCAGACCGGCGCCGGTAACAACTGGGCTAAGGGACACTACACGGAGGGTGCCGAGCTTATCGACTCGGTGCTCGACGTCGTCCGCAAGGAGGCCGAGAGCTGTGACTGCCTGCAGGGGTTCCAGATCACGCACTCTCTTGCGGCACCATGGACTCGGTGCGCGCCGGCCCCTACGGCCAGCTCTTCCGCCCGGACAACTTCGTGTTCGGCCAGACGGGCGCCGGTAACAACTGGGCCAAGGGACACTACACGGAGGGCGCCGAGCTCATCGACTCGGTGCTCGATGTCGTCCGCAAGGAGGCGGAGAGCTGCGACTGCCTGCAGGGATTCCAGATCACGCACTCGCTCGCGGCACCATGGACTCGGTCCGCGCCGGCCCCTACGGCCAGCTCTTCCGTCCGGACAACTTTGTGTTCGGCCAGACGGGCGCCGGTAACAACTGGGCCAAGGGACACTACACGGAGGGTGCCGAGCTTATCGACTCGGTGCTTGACGTCGTCCGTAAGGAGGCTGAGAGCTGTGATTGCCTGCAGGGTTTCCAGATCACCCACTCGCTTGCGGCACCATGGACTCGGTGCGCGCCGGCCCCTACGGCCAGCTCTTCCGCCCGGACAACTTCGTGTTCGGCCAGACGGGCGCCGGTAACAACTGGGCCAAGGGACACTACACGGAGGGTGCCGAGCTTATCGACTCGGTTCTCGACGTCGTCCGCAAGGAGGCTGAGAGCTGTGACTGCCTTCAGGGTTTCCAGATCACGCACTCGCTGGCGGCACCATGGACTCGGTTCGCGCTGGCCCCTACGGTCAGCTTTTCCGCCCCGACAATTTTGTGTTTGGCCAGACAGGTGCCGGTAATAACTGGGCCAAAGGACACTACACTGAGGGTGCTGAGCTCATTGACTCAGTGCTGGACGTCGTCCGTAAAGAGGCGGAAAGCTGTGACTGTTTGCAGGGTTTCCAGATCACGCACTCGCTTGTGGTACCATGGACTCGGTCCGCGCTGGTCCATACGGCCAGCTTTTCCGTCCGGACAACTTCGTGTTCGGCCAGACTGGTGCCGGTAACAACTGGGCCAAGGGTCACTACACGGAGGGTGCTGAGCTCATCGACTCGGTGCTTGACGTCGTCCGCAAGGAGGCTGAGAGCTGTGACTGCCTTCAGGGTTTCCAGATCACCCACTCGCTTGCGGCACCATGGACTCGGTTCGTGCTGGTCCCTACGGCCAGCTCTTCCGCCCCGACAATTTTGTGTTCGGCCAGACAGGTGCCGGTAACAATTGGGCTAAGGGACACTACACTGAGGGCGCCGAGCTCATTGACTCCGTTCTGGACGTCGTCCGTAAGGAGGCGGAAAGCTGTGACTGTTTGCAGGGTTTCCAGATCACGCACTCGCTTGCGGCACGATGGACTCGGTCCGCGCTGGTCCGTACGGCCAGCTCTTCCGTCCGGATAACTTTGTCTTTGGACAGACAGGAGCTGGTAACAATTGGGCCAAGGGACACTACACGGAGGGTGCAGAGCTTATTGACTCGGTGCTGGACGTTGTCCGCAAAGAGGCTGAGAGCTGTGACTGCCTACAAGGTTTCCAGTTCACGCACTCGCTCGCGGCACGATGGACTCGGTCCGCGCTGGTCCGTACGGCCAGCTCTTCCGTCCGGATAACTTTGTCTTTGGACAGACAGGAGCTGGTAACAATTGGGCCAAGGGACACTACACGGAGGGTGCAGAGCTTATTGACTCGGTGCTGGACGTTGTCCGCAAAGAGGCTGAGAGCTGTGACTGCCTACAAGGTTTCCAGTTCACGCACTCGCTCGCGGCACGATGGACTCGGTCCGCGCTGGTCCGTACGGCCAGCTCTTCCGTCCGGATAACTTTGTCTTTGGACAGACAGGAGCTGGTAACAATTGGGCCAAGGGACACTACACGGAGGGTGCAGAGCTTATTGACTCGGTGCTGGACGTTGTCCGCAAAGAGGCTGAGAGCTGTGACTGCCTACAAGGTTTCCAGTTCACGCACTCGCTCGCGGTACCATGGACTCCGTTCGCGCTGGCCCCTACGGTCAGCTTTTCCGCCCAGACAATTTCGTGTTCGGACAGACTGGCGCTGGTAACAACTGGGCCAAGGGACACTACACTGAGGGCGCTGAGCTGATTGACTCGGTGCTTGACGTCGTTCGCAAGGAGGCAGAGAGCTGTGATTGCCTTCAGGGTTTCCAGATCACGCACTCGCTTGCGGCACCATGGACTCGGTCCGTGCTGGCCCTTACGGCCAGCTCTTCCGTCCGGACAACTTCGTGTTCGGCCAAACAGGCGCTGGTAACAACTGGGCCAAGGGACACTACACAGAGGGTGCCGAGCTTATCGACTCGGTGCTTGATGTCGTCCGCAAGGAGGCTGAGAGCTGTGACTGCCTGCAGGGTTTCCAGATCACCCACTCGCTTGTGGAACTATGGACTCAGTCCGCGCTGGTCCTTATGGCCAACTCTTCCGCCCAGACAATTTTGTGTTCGGCCAGACAGGCGCCGGTAACAATTGGGCGAAAGGACATTATACCGAGGGTGCCGAGCTTATTGATTCGGTGCTTGACGTCGTTCGCAAAGAGGCAGAGAGCTGTGATTGCCTTCAGGGTTTCCAAATCACGCACTCGCTTGTGGCACAATGGACTCGGTCCGCGCCGGCCCATATGGTCAGCTTTTCCGCCCAGACAATTTTGTTTTTGGCCAGACAGGCGCCGGTAACAATTGGGCGAAAGGACATTATACCGAGGGTGCCGAGCTGATTGATTCCGTGCTAGACGTTGTCCGCAAGGAAGCAGAAAGTTGCGATTGTCTTCAAGGTTTTCAAATTACGCACTCACTTGCGGCACCATGGACTCGGTCCGCGCTGGTCCCTACGGCCAGCTCTTCCG+CCGGACAACTTTGTGTT+GGCCAGACAGGCGCCGGTAACAACTGGGCCAAGGGACACTACACGGAGGGTGCCGAGCTTATTGACTCGGTGCTGGACGTCGTCCGCAAGGAGGCTGAGAGCTGTGACTGCCT+CAGGGTTTCCAGATCACGCACTCGCTTGGTGGAGGTACAGGGTCTGGTATGGGCACCCTCCTCATTAGCAAAATCCGCGAGGAGTATCCCGACCGCATCATGAACACCTTCAGCGTGGTGCCCTCTCCTAAAGTCTCGGACACTGTGGTCGAGCCCTACAACGCCACACTGTCCGTCCATCAGCTAGTAGAGAACACAGACGAGACCTATTGTATTGATAACGAGGCCCTGTACGATGAGGAGGAACTGGATCTGGAATGGGAACTCTATTGATTTCTAAGATAAGAGAAGAGTATCCAGATCGTATGATGATGACTTTCTCAGTGTTTCCTTCTCCTAAGGTCTCTGACACTGTTGTTGAGCCATACAATGCAACTCTCTCTGTGCATCAGCTTGTCGAAAACGCTGACGAGTGTATGGTTTTGGACAATGAGGCTCTCTACGATGCGGGGGCACAGGCTCTGGAATGGGCACTCTCCTTATCAGCAAGATCCGAGAAGAATACCCTGATCGCATCATGAATACCTTCAGTGTGGTGCCTTCACCCAAAGTGTCTGACACCGTGGTCGAGCCCTACAATGCCACCCTCTCCGTCCATCAGTTGGTAGAGAATACTGATGAGACCTATTGCATTGACAACGAGGCCCTCTATGATGAGGAGGAACTGGTTCTGGAATGGGAACTCTTTTGATTTCCAAAATTCGTGAAGAATATCCAGATCGTATCATGATGACTTTCTCCGTAGTGCCAAGTCCAAAAGTGTCCGACACGGTCGTCGAGCCGTACAACGCAACTCTTTCTGTTCATCAACTTGTTGAAAACACCGACGAAACATTCTGTATTGACAACGAAGCCTTGTATGACGTGGTGGTACCGGTTCCGGTATGGGTACGCTTTTGATCTCGAAGATTAGGGAAGAGTTTCCTGATCGTATGATGGCCACCTTCTCCGTCTTGCCCTCTCCGAAGACTTCTGACACCGTTGTCGAACCATACAATGCCACGTTGTCTGTGCACCAATTGGTAGAACACTCTGATGAAACATTCTGTATCGATAACGAAGCACTTTATGACGCGGTGGTACTGGCTCGGGTATGGGCACGCTTCTTATCTCGAAGATCCGCGAGGAATACCCGGATCGCATCATGTGCACGTACTCCGTGTGCCCGTCCCCGAAGGTGTCGGACACGGTGGTGGAGCCCTACAATGCGACGCTGTCGGTGCACCAGCTCGTTGAAAACGCTGACGAAGTCATGTGCCTGGACAACGAGGCCCTGTACGACGTGGCGGTACCGGTTCTGGTATGGGCACGCTTCTGATCTCCAAGATCCGTGAGGAGTACCCGGACCGTATCATGTGCACGTACTCGGTGTGCCCGTCGCCCAAGGTGTCGGACACGGTCGTGGAGCCCTACAACGCCACGCTGTCGGTGCACCAGCTTGTCGAGAACGCCGACGAGGTCATGTGCCTGGATAACGAGGCGCTGTACGACGTGGCGGTACCGGTTCCGGTATGGGCACGCTTCTTATCTCCAAGATCCGTGAGGAGTACCCGGACCGTATCATGTGCACGTACTCGGTGTGCCCGTCGCCCAAGGTGTCGGACACGGTCGTGGAGCCCTACAACGCGACGCTGTCCGTGCACCAGCTTGTCGAGAACGCCGATGAGGTCATGTGCCTGGATAACGAGGCCCTGTACGACGTGGCGGTACCGGTTCCGGTATGGGCACGCTTCTTATTTCCAAGATCCGCGAAGAGTACCCCGACCGTATCATGTGCACGTACTCGGTGTGCCCGTCGCCCAAGGTGTCGGACACGGTCGTGGAGCCTTACAACGCGACGCTGTCGGTGCACCAGCTTGTCGAGAACGCCGATGAGGTCATGTGCCTGGATAACGAGGCCCTGTACGACGTGGCGGTACCGGTTCCGGTATGGGTACGCTTCTTATCTCCAAGATTCGTGAGGAGTACCCGGACCGTATCATGTGCACGTACTCGGTCTGCCCGTCGCCTAAGGTGTCGGACACGGTCGTCGAGCCCTACAACGCTACGCTGTCCGTCCACCAGCTCGTTGAGAACGCCGATGAGGTCATGTGCCTGGATAACGAGGCCCTGTACGACGTGGCGGGACCGGCTCCGGTATGGGAACGCTTCTTATCTCGAAGATCCGAGAAGAATACCCGGACCGTATCATGTGCACGTACTCGGTGTGTCCATCCCCCAAAGTGTCGGACACGGTTGTGGAGCCTTACAACGCTACGCTGTCTGTGCACCAGCTTGTCGAGAATGCCGATGAGGTCATGTGCCTCGATAACGAGGCCTTATACGACGTGGTGGTACCGGTTCTGGTATGGGTACCCTCCTCATCTCGAAGATTCGTGAAGAGTACCCAGACCGTATCATGTGCACGTACTCGGTGTGCCCATCGCCAAAGGTCTCGGATACCGTCGTTGAGCCATACAACGCCACGCTTTCGGTCCACCAGCTTGTCGAGAACGCCGATGAGGTGATGTGTCTTGACAACGAAGCCCTTTACGATGTGGCGGTACCGGCTCCGGTATGGGAACGCTTCTTATCTCAAAGATCCGAGAAGAGTACCCGGACCGTATCATGTGCACGTACTCGGTATGTCCATCTCCCAAGGTGTCGGACACTGTTGTAGAGCCCTACAATGCTACGCTGTCTGTACATCAGCTTGTCGAAAACGCCGATGAGGTCATGTGCCTGGATAATGAGGCCTTGTACGACGCGGTGGTACCGGCTCTGGTATGGGCACGCTTCTTATCTCAAAGATCCGAGAAGAGTATCCGGACCGTATTATGTGCACATACTCCGTGTGTCCGTCCCCCAAGGTGTCGGACACTGTCGTCGAGCCCTACAACGCTACGTTGTCGGTGCACCAGCTCGTGGAGAACGCTGATGAAGTCATGTGCCTTGACAATGAGGCCCTGTATGACCACTGTCGTCGAGCCCTACAACGCTACGTTGTCGGTGCACCAGCTCGTGGAGAACGCTGATGAAGTCATGTGCCTTGACAATGAGGCCCTGACTGTCGTCGAGCCCTACAACGCTACGTTGTCGGTGCACCAGCTCGTGGAGAACGCTGATGAAGTCATGTGCCTTGACAATGAGGCCCTGTATGACGCGGTGGTACCGGCTCTGGTATGGGCACGCTTCTTATCTCAAAGATCCGAGAAGAGTATCCGGACCGTATTATGTGCACATACTCCGTGTGTCCGTCCCCCAAGGTGTCGGACACTGTCGTCGAGCCCTACAACGCTACGTTGTCGGTGCACCAGCTCGTGGAGAACGCTGATGAAGTCATGTGCCTTGACAATGAGGCCCTGTATGACCACTGTCGTCGAGCCCTACAACGCTACGTTGTCGGTGCACCAGCTCGTGGAGAACGCTGATGAAGTCATGTGCCTTGACAATGAGGCCCTGACTGTCGTCGAGCCCTACAACGCTACGTTGTCGGTGCACCAGCTCGTGGAGAACGCTGATGAAGTCATGTGCCTTGACAATGAGGCCCTGTATGACGCGGTGGTACCGGCTCTGGTATGGGCACGCTTCTTATCTCAAAGATCCGAGAAGAGTATCCGGACCGTATTATGTGCACATACTCCGTGTGTCCGTCCCCCAAGGTGTCGGACACTGTCGTCGAGCCCTACAACGCTACGTTGTCGGTGCACCAGCTCGTGGAGAACGCTGATGAAGTCATGTGCCTTGACAATGAGGCCCTGTATGACCACTGTCGTCGAGCCCTACAACGCTACGTTGTCGGTGCACCAGCTCGTGGAGAACGCTGATGAAGTCATGTGCCTTGACAATGAGGCCCTGACTGTCGTCGAGCCCTACAACGCTACGTTGTCGGTGCACCAGCTCGTGGAGAACGCTGATGAAGTCATGTGCCTTGACAATGAGGCCCTGTATGACGTGGCGGTACCGGTTCCGGTATGGGTACGCTTCTTATCTCGAAGATTCGTGAGGAGTACCCCGATCGCATCATGTGCACATACTCGGTCTGCCCGTCGCCCAAGGTATCGGACACGGTCGTGGAGCCCTATAACGCTACGCTATCGGTACACCAGCTTGTCGAGAACGCCGATGAGGTCATGTGCCTGGACAATGAGGCCCTGTACGACGTGGCGGTACCGGTTCCGGTATGGGTACGCTTCTTATCTCGAAGATTCGTGAGGAGTACCCAGACCGTATTATGTGCACGTACTCGGTCTGCCCGTCGCCCAAGGTGTCGGACACCGTCGTGGAGCCCTACAACGCCACGTTGTCGGTGCATCAGCTGGTCGAGAACGCCGATGAGGTCATGTGCCTGGATAACGAGGCCCTGTACGATGTGGCGGTACTGGTTCTGGGATGGGAACGCTTCTTATTTCAAAGATTCGTGAGGAATACCCCGACCGTATCATGTGCACATACTCCGTGTGCCCTTCGCCAAAGGTATCGGATACGGTCGTGGAGCCTTACAATGCTACGCTTTCCGTGCACCAGCTCGTTGAAAACGCTGATGAAGTCATGTGCTTGGACAATGAGGCCCTGTACGATGTGGTGGTACCGGCTCCGGAATGGGCACACTTCTTATTTCGAAAATTCGGGAAGAGTATCCCGACCGAATCATGTGCACCTACTCGGTGTGCCCATCGCCTAAGGTATCAGATACGGTAGTAGAGCCTTACAATGCCACATTGTCGGTGCACCAGCTAGTCGAGAACGCTGATGAAGTTATGTGCTTAGACAACGAAGCTTTGTACGACGTGGCGGTACCGGTTCTGGTATGGGCACGCTTCTTATCTC+AAGATCCGTGAAGAGTACCCGGACCGTATCATGTGCACGTACTCGGTGTGCCCGTCGCCCAAGGTGTCGGACACGGTCGT+GAGCCCTACAACGCTACGCTGTCGGTGCACCAGCTTGTCGAGAACGCCGATGAGGTCATGTGCCTGGACAACGAGGCCCTGTACGACATCTGCTTCCGCACACTCAAACTCACAACCCCCACATACGGAGACCTCAACCATCTCGTCTCCGCCACAATGAGCGGTGTGACCACTTGCTTGAGGTTTCCAGGCCAGTTGAACGCTGATCTCCGTAAATTGGCGGTCAACATGGTGCCCTTCCCCCGACTGCACTTCTTCATGCCTGGCTTCGCGCCTCTGACTAGCAGGGGAAGCCAATCTGTTTCCGTACCCTCAAGCTCGCTAATCCTACCTTTGGTGATCTTAACCATCTCATCTCTGCTACAATGAGTGGTGTCACTTGCTGTCTTCGTTTCCCTGGCCAGCTTAACTCTGACCTTAGGAAACTCGCTGTGAACCTTATCCCATTCCCAAGGCTTCACTTCTTCATGGTTGGTTTCGCACCATTGACATCGAGAGGATCACAATCTGCTTCCGCACTCTGAAGCTGACCACACCAACCTACGGGGATCTGAACCACCTTGTCTCAGCCACCATGAGTGGTGTCACCACCTGCCTCCGTTTCCCTGGCCAGCTCAATGCTGACCTCCGCAAGTTGGCAGTCAACATGGTCCCCTTCCCACGTCTCCATTTCTTTATGCCTGGCTTTGCCCCTCTCACCAGCCGTGGAAGCCAATCTGTTTCCGCACGCTCAAGCTCACCACACCGACCTACGGAGATTTGAATCATCTCGTTTCGATGACGATGAGTGGTGTCACCACCTGTCTTCGTTTCCCGGGACAGCTGAATGCAGATTTGCGCAAATTAGCTGTTAACATGGTTCCGTTCCCACGTCTTCATTTCTTCATGCCCGGATTTGCTCCGCTCACATCCAGAGGAAGCCAATCTGTCAAAGGACCTTAAAGTTGAATCAACCTTCTTATGGAGATTTGAACAACTTGGTCTCGAGCGTCATGTCTGGTGTGACAACTTCATTGCGTTATCCCGGCCAATTGAACTCTGATTTGAGAAAGTTGGCTGTTAATCTTGTCCCATTCCCACGTTTACATTTCTTCATGGTCGGCTACGCTCCATTGACGGCAATTGGCTCTCAATTTGCTTCCGTACGCTGAAACTCACCACCCCTACGTATGGTGACCTGAACCACTTGGTGTGCGCTGCTATGTCTGGCATCACGACGTGTCTACGTTTTCCCGGCCAGCTGAACTCCGACCTGCGTAAGCTAGCCGTGAACCTGATTCCGTTTCCACGTCTCCACTTCTTCATGATCGGATTCGCTCCGCTGACGTCGCGAAGCTCGCAATTTGCTTCCGCACGCTCAAGCTCACCACCCCCACCTACGGTGACCTGAACCACCTGGTGTGCGCCGCTATGTCCGGCATCACCACGTGCCTGCGTTTCCCGGGTCAGCTGAACTCGGACCTGCGGAAGCTGGCGGTGAACTTGATTCCGTTCCCGCGTCTTCACTTCTTCATGATTGGTTTCGCCCCGCTGACCTCGCGTGGCTCGCAATTTGCTTCCGCACCTTGAAATTGACGAACCCGACGTACGGTGATCTGAACCACCTGGTGTGCGCCGCCATGTCCGGCATCACCACGTGCCTCCGTTTCCCCGGCCAACTGAACTCGGTCCTGAAGCTGTTTGCCGTTAACCTGATCCCGTTCCCCCGTCTCCATTTCTTCATGATCGGTTTCGCTCCGCTGACGTCGCGTGGATCTCAATTTGCTTCCGTACGCTCAAGCTCACCACCCCCACGTACGGTGACCTGAACCACTTGGTGTGCGCCGCCATGTCCGGTATCACGACGTGCCTGCGTTTCCCCGGTCAGCTGAACTCGGACCTGCGTAAGCTGGCCGTGAATCTGATTCCGTTCCCGCGTCTCCACTTCTTCATGATTGGTTTCGCCCCGCTGACGTCGCGTGGCTCGCAATTTGCTTCCGTACCCTGAAGCTCACGACCCCCACCTACGGTGACCTGAACCACCTGGTGTGCGCCGCCATGTCCGGCATTACCACGTGCCTGCGTTTCCCCGGTCAGCTGAACTCGGACCTGCGTAAGCTTGCCGTGAACCTGATCCCGTTCCCGCGTCTCCACTTCTTCATGATCGGTTTCGCCCCGCTGACGTCGCGCGGCTCGCAATTTGCTTTCGCACGCTGAAGCTCACCACCCCCACTTACGGTGACCTGAACCACTTGGTATGTGCCGCTATGTCTGGCATCACCACTTCCCTGCGATTCCCTGGACAGCTAAACTCGGACCTGCGGAAGCTGGCCGTGAACCTAATCCCTTTCCCCCGTCTCCATTTCTTCATGATTGGTTTTGCCCCGCTGACGTCACGTGGATCGCAATCTGTTTCCGTACCCTGAAGCTCACGACGCCAACCTACGGTGACCTGAACCACCTTGTGTGTGCTGCCATGTCGGGTATCACGACCAGTCTGCGTTTCCCAGGTCAGCTGAACTCGGATCTTCGTAAGCTTGCCGTCAACCTTATCCCGTTCCCACGTCTCCACTTCTTCATGGTCGGTTTCGCTCCGCTCACCTCGCGCGGCTCGCAATTTGCTTCCGTACGCTGAAGCTCACCACCCCCACCTATGGTGACCTGAACCACTTGGTGTGTGCCGCTATGTCGGGCATCACCACTTCCCTACGATTTCCGGGACAGCTGAACTCGGACCTGCGGAAGCTGGCCGTGAACCTGATCCCGTTCCCTCGTCTTCACTTCTTCATGATTGGTTTCGCGCCGCTGACGTCACGTGGTTCGCAATTTGCTTCCGCACGCTGAAGCTCACGACTCCTACTTACGGTGATCTGAATCACTTGGTGTGTGCTGCTATGTCTGGTATCACGACGTGCCTACGTTTTCCCGGTCAGCTGAACTCAGACCTCCGTAAGCTGGCTGTGAACTTGATTCCGTTTCCACGTCTTCACTTCTTCATGATTGGTTTCGCTCCGCTGACGTCGCGAAGCTCGCAATTTGCTTCCGCACGCTGAAGCTCACGACTCCTACTTACGGTGATCTGAATCACTTGGTGTGTGCTGCTATGTCTGGTATCACGACGTGCCTACGTTTTCCCGGTCAGCTGAACTCAGACCTCCGTAAGCTGGCTGTGAACTTGATTCCGTTTCCACGTCTTCACTTCTTCATGATTCTTGGTGTGTGCTGCTATGTCTGGTATCACATTTGCTTCCGCACGCTGAAGCTCACGACTCCTACTTACGGTGATCTGAATCACTTGGTGTGTGCTGCTATGTCTGGTATCACGACGTGCCTACGTTTTCCCGGTCAGCTGAACTCAGACCTCCGTAAGCTGGCTGTGAACTTGATTCCGTTTCCACGTCTTCACTTCTTCATGATTGGTTTCGCTCCGTTGACGTCGCGAAGCTCGCAATTTGCTTCCGCACGCTGAAGCTCACGACTCCTACTTACGGTGATCTGAATCACTTGGTGTGTGCTGCTATGTCTGGTATCACGACGTGCCTACGTTTTCCCGGTCAGCTGAACTCAGACCTCCGTAAGCTGGCTGTGAACTTGATTCCGTTTCCACGTCTTCACTTCTTCATGATTCTTGGTGTGTGCTGCTATGTCTGGTATCACATTTGCTTCCGCACGCTGAAGCTCACGACTCCTACTTACGGTGATCTGAATCACTTGGTGTGTGCTGCTATGTCTGGTATCACGACGTGCCTACGTTTTCCCGGTCAGCTGAACTCAGACCTCCGTAAGCTGGCTGTGAACTTGATTCCGTTTCCACGTCTTCACTTCTTCATGATTGGTTTCGCTCCGTTGACGTCGCGAAGCTCGCAATTTGCTTCCGCACGCTGAAGCTCACGACTCCTACTTACGGTGATCTGAATCACTTGGTGTGTGCTGCTATGTCTGGTATCACGACGTGCCTACGTTTTCCCGGTCAGCTGAACTCAGACCTCCGTAAGCTGGCTGTGAACTTGATTCCGTTTCCACGTCTTCACTTCTTCATGATTCTTGGTGTGTGCTGCTATGTCTGGTATCACATTTGCTTCCGCACATTGAAGCTCACCACCCCCACTTATGGTGACCTGAACCACTTGGTGTGTGCCGCCATGTCCGGTATTACCACGTGCCTTCGTTTCCCCGGTCAGCTGAACTCGGACCTGCGTAAGCTGGCCGTGAACCTGATCCCGTTCCCGCGTCTCCACTTCTTTATGATTGGTTTCGCTCCTCTGACATCGCGCGGCTCGCAATTTGCTTCCGTACGCTGAAGCTCACCACGCCCACCTACGGTGACCTGAACCACCTGGTGTGCGCCGCCATGTCCGGTATCACCACGTGCCTGCGTTTCCCCGGTCAGTTGAACTCGGACCTGCGTAAGCTGGCCGTGAACCTGATCCCGTTCCCGCGTCTCCACTTCTTTATGATTGGTTTCGCTCCGCTGACCTCGCGTGGATCTCAATTTGCTTTCGAACGTTGAAGCTTACCACCCCAACTTATGGTGATTTGAACCACCTGGTGTGCGCCGCCATGTCTGGTATAACTACATGCCTGCGTTTTCCAGGCCAACTGAATTCTGATCTTCGTAAATTGGCTGTAAACCTTATTCCATTCCCGCGTCTTCACTTCTTTATGATTGGCTTTGCTCCGTTGACGTCGCGTGGCTCGCAATTTGCTTTCGTACGTTAAAACTCACCACGCCTACTTACGGTGACTTGAATCACTTGGTGTGTGCTGCCATGTCTGGTATTACCACATGCTTGCGATTTCCTGGTCAGCTAAATTCAGACTTGCGGAAGTTGGCCGTCAATCTGATTCCATTTCCCCGTCTTCATTTTTTTATGATTGGTTTTGCACCTTTGACATCACGTGGCTCGCAATTTGCTTCCGCACGCTGAAGCTCACCACCCCCAC+TACGGTGACCTGAACCAC+TGGTGTGTGCCGCCATGTCTGGTATCACCACGTGCCTGCGTTTCCCCGGTCAGCTGAACTCGGACCTGCGTAAGCTGGCCGTGAACCTGATCCCGTTCCCACGTCT+CACTTCTTCATGATTGGTTTCGCTCCGCTGACGTCGCGTGGCTCGCAGCAGTATCGTGCACTTACCGTTCCCGAACTCACCCAGCAGATGTTCGATGCCAAAAACATGATGGCTGCCTGCGACCCACGTCACGGCCGTTATCTGACGGTCGCCGCCGTCTTCCGTGGTCGCATGTCCATGAAGGAGGTGGACGAGCAGATGCTCAACGTCCAGAACAAGAACAGCAGCTACTTCGTTGAATGGATCCCAAACAACGGCAATACAGTGCCTTGAGTGTTCCTGAACTGACCCAGCAGATGTGGGATGCAAAGAACATGATGTGTGCTGCTGACCCTCGTCATGGACGTTACTTGACTGCATCCGCTGTGTTCCGTGGAAAGCTGAGCACCAAAGAGGTTGACGAGCAGATGATGAACATTCAGAACAAGAACTCATCCTACTTTGTGGAATGGATCCCAAACAACGGCAGTATCGAGCTCTCACAGTGCCGGAACTCACCCAGCAGGTCTTCGATGCCAAGAACATGATGGCTGCCTGTGACCCCCGCCACGGCCGATACCTCACCGTGGCTGCTGTCTTCCGTGGTCGGATGTCCATGAAGGAGGTCGATGAGCAGATGCTTAACGTGCAGAACAAGAACAGCAGCTACTTTGTGGAATGGATCCCCAACAATGGCAATACAGATCGCTCACCGTTCCAGAGCTCACACAACAAATGTTCGACGCCAAGAATATGATGGCAGCCTGCGATCCAAGACACGGTCGCTACCTGACAGTTGCTGCAATGTTCCGCGGAAGAATGAGCATGAAAGAAGTCGACGAGCAAATGCTCAATGTGCAAAATAAAAACTCCTCGTACTTTGTTGAATGGATTCCAAACAACGATCATTTAGATCTTTGACTGTCCCTGAATTAACACAGCAAATGTTTGATGCCAAGAACATGATGGCTGCTGCCGATCCAAGAAACGGTAGATACCTTACCGTTGCAGCCTTCTTTAGAGGTAAAGTTTCCGTTAAGGAGGTGGAAGATGAAATGCATAAAGTGCAATCTAAAAACTCAGACTATTTCGTGGAATGGATCCCCAACAATGGCAGTACCGTGCGCTGACGGTGCCCGAGCTCACGCAGCAGCAGTTTGATGCGAAGAACATGATGTGCGCTGCAGACCCTCGCCATGGCCGTTATTTGACCGCCGCGTGTATGTTCCGCGGTCGCATGAGTACGAAGGAGGTGGACGAGCAAATGCTGAACGTGCAGAACAAGAATTCGTCGTACTTCGTCGAGTGGATTCCGAATAACAGCAGTACCGTGCCCTGACGGTGCCCGAGCTGACCCAGCAGCAGTTCGACGCAAAGAACATGATGTGCGCCGCCGACCCTCGTCACGGCCGCTATTTAACTGCCGCGTGTATGTTCCGCGGACGTATGAGCACGAAGGAGGTTGATGAGCAGATGCTGAACGTGCAGAACAAGAACTCGTCGTACTTCGTCGAGTGGATCCCTAACAACAGCAGTACCGTGCCCTGACGGTGCCGGAGCTGACGCAGCAGCAGTTCGATGCTAAGAACATGATGTGCGCCGCGGACCCTCGCCACGGCCGCTATTTAACTGCCGCGTGTATGTTCCGCGGACGTATGAGCACGAAGGAGGTCGATGAGCAGATGCTCAACGTGCAGAACAAGAACTCGTCGTACTTCGTCGAGTGGATCCCCAACAACAGCAGTACCGTGCCCTTACGGTGCCCGAGCTGACCCAGCAGCAATTCGATGCTAAGAACATGATGTGCGCCGCCGACCCTCGCCACGGCCGCTATTTAACTGCCGCGTGTATGTTCCGCGGACGTATGAGTACGAAGGAGGTTGATGAGCAGATGCTGAACGTGCAGAACAAGAACTCGTCGTACTTCGTTGAGTGGATCCCCAACAACAGCAGTACCGTGCCCTGACGGTGCCCGAGCTGACCCAGCAGCAGTTCGATGCTAAGAACATGATGTGTGCCGCCGACCCTCGCCACGGCCGCTATTTAACTGCCGCGTGTATGTTCCGCGGACGTATGAGCACGAAGGAGGTTGACGAGCAGATGCTCAACGTGCAGAACAAGAACTCGTCGTACTTCGTCGAGTGGATCCCCAACAACAGCAATACCGTGCCCTGACGGTGCCTGAGCTGACGCAGCAGCAGTTCGATGCTAAGAACATGATGTGCGCCGCTGACCCTCGTCACGGCCGCTATTTAACTGCCGCCTGTATGTTCCGCGGACGTATGAGCACGAAGGAGGTTGATGAGCAAATGCTGAACGTGCAGAACAAGAACTCGTCGTACTTTGTCGAGTGGATCCCCAACAATAGCAGTACCGTGCCCTTACGGTGCCAGAGCTGACCCAGCAGCAGTTCGACGCCAAGAACATGATGTGTGCCGCTGATCCTCGTCACGGTCGTTACCTGACCGCCGCTTGTATGTTCCGTGGTCGTATGAGCACCAAGGAGGTCGACGAGCAGATGCTCAACGTCCAGAACAAGAACTCGTCGTACTTCGTTGAGTGGATCCCGAACAACAGCAGTACCGTGCCTTGACGGTGCCTGAGCTGACACAGCAGCAGTTTGATGCTAAAAACATGATGTGCGCCGCCGACCCTCGCCACGGTCGCTATTTAACTGCCGCTTGTATGTTCCGCGGACGCATGAGCACGAAGGAGGTTGATGAGCAAATGCTGAACGTGCAGAACAAAAACTCGTCGTACTTTGTCGAGTGGATTCCTAACAACAGCAGTACCGTGCGTTGACGGTGCCCGAGCTGACGCAGCAGCAGTTCGATGCTAAAAACATGATGTGTGCCGCCGACCCCCGCCATGGCCGCTATTTAACTGCCGCGTGTATGTTCCGCGGACGTATGAGTACGAAGGAGGTGGATGAGCAAATGCTGAACGTGCAAAACAAGAACTCGTCGTACTTCGTCGAGTGGATTCCCAACAACAGCAGTACCGTGCGTTGACGGTGCCCGAGCTGACGCAGCAGCAGTTCGATGCTAAAAACATGATGTGTGCCGCCGACCCCCGCCATGGCCGCTATTTAACTGCCGCGTGTATGTTCCGCGGACGTATGAGTACGAAGGAGGTGGATGAGCAAATGCTGAACGTGCAAAACAAGAACTCGTCGTACTTCGTCGAGTGGATTCCCAACAACAGCAGTACCGTGCGTTGACGGTGCCCGAGCTGACGCAGCAGCAGTTCGATGCTAAAAACATGATGTGTGCCGCCGACCCCCGCCATGGCCGCTATTTAACTGCCGCGTGTATGTTCCGCGGACGTATGAGTACGAAGGAGGTGGATGAGCAAATGCTGAACGTGCAAAACAAGAACTCGTCGTACTTCGTCGAGTGGATTCCCAACAACAGCAGTACCGCGCCCTGACGGTGCCCGAGCTGACCCAGCAGCAGTTCGATGCTAAGAACATGATGTGTGCCGCCGACCCTCGCCATGGCCGCTATTTAACTGCAGCTTGTATGTTCCGCGGACGCATGAGCACGAAGGAGGTTGATGAGCAGATGCTGAACGTGCAGAACAAGAACTCGTCATACTTCGTCGAGTGGATCCCCAACAACAGCAGTACCGTGCTCTTACGGTGCCGGAGCTGACCCAGCAGCAGTTCGATGCCAAGAACATGATGTGCGCCGCTGACCCTCGCCACGGCCGCTATTTAACTGCCGCGTGTATGTTCCGCGGACGTATGAGCACGAAGGAGGTTGACGAGCAGATGCTGAACGTGCAGAACAAGAACTCATCGTACTTCGTCGAGTGGATCCCCAACAACAGCAATACCGTGCCCTGACGGTTCCTGAATTAACTCAACAGCAGTTTGACGCGAAGAATATGATGTGTGCCGCTGACCCTCGTCATGGTCGCTATTTAACTGCCGCATGTATGTTTCGTGGACGCATGAGCACCAAGGAGGTTGACGAGCAGATGCTTAACGTGCAGAATAAGAATTCCTCCTACTTTGTCGAGTGGATTCCCAACAACAGCAGTATCGTGCGTTGACAGTACCTGAGTTGACCCAGCAGCAGTTTGATGCTAAAAACATGATGTGCGCCGCTGACCCTCGCCACGGTCGCTATTTAACTGCCGCGTGTATGTTTCGAGGGCGCATGAGCACGAAGGAAGTTGATGAGCAGATGCTGAATGTGCAAAACAAAAATTCTTCTTATTTCGTCGAGTGGATTCCCAACAACAGCAGTACCGTGCCCTGACGGTGCCCGAGCTGACCCAGCAGCAGTTCGATGCTAAGAACATGATGTGTGCCGCCGACCCTCGCCACGGCCGCTATTTAACTGCCGCGTGTATGTTCCGCGGACGTATGAGCACGAAGGAGGTTGATGAGCAGATGCTGAACGTGCAGAACAAGAACTCGTCGTACTTCGTCGAGTGGATCCCCAACAACATCAAGACCGCCGTCTGCGACATTCCACCACGAGGCCTCAAGATGGCCGCCACCTTCATCGGCAACAGCACCGCCATCCAGGAGCTCTTCAAGCGCATCTCAGAGCAGTTCACGGCTATGTTCAGGCGCAAGGCTTTCCTGCATTGGTACACCGGAGAGGGCATGGATGAGATGGAGTTCACAGAGGCCGAGAGCAACATGAACGACCTGTCAAGTCCAGTGTCTGTGATATTGCACCAAAGGGTTTGAAAATGGCGTCTACTTTCATTGGTAACTCAACCTCAATCCAGGAGATGTTTAGGCGTGTGAGCGAACAGTTCACAGCTATGTTCAGGAGAAAGGCTTTCCTTCATTGGTACACAGGAGAAGGCATGGACGAGATGGAGTTCACTGAAGCAGAGAGTAACATGAATGATCTTTCAAGACAGCCGTCTGTGACATCCCACCTCGTGGCCTCAAGATGGCAGTCACCTTCATTGGCAATAGCACAGCCATCCAGGAGCTCTTCAAGCGCATCTCGGAGCAGTTCACTGCCATGTTCCGCCGGAAGGCCTTCCTCCACTGGTACACAGGCGAGGGCATGGACGAGATGGAGTTCACCGAGGCTGAGAGCAACATGAACGACCTCTCAAAACTGCTGTTTGTGATATTCCGCCAAGAGGAGTAAAGATGGCGGCAACATTCGTCGGAAATTCCACTGCAATTCAAGAGCTTTTCAAGAGAATCAGTGAGCAATTCACAGCTATGTTCCGTAGAAAAGCTTTCCTTCATTGGTACACTGGAGAAGGTATGGACGAAATGGAGTTTACTGAAGCTGAGAGTAACATGAATGACTTGTGCAAACTGCTGTGTGTTCTGTCGCTCCTCAAGGTTTGGACATGGCTGCTACTTTCATTGCTAACTCCACATCTATTCAAGAGCTATTCAAGAGAGTTGGTGACCAATTTTCCGCTATGTTCAAAAGAAAAGCTTTCTTGCACTGGTATACTAGTGAAGGTATGGACGAATTGGAATTCTCTGAGGCTGAATCTAATATGAATGATCTGTTAAGGCTAGCGTGTGCGACATCCCGCCCAAGGGCCTGAAGATGAGTACGACGTTTATCGGCAACTCGACCGCGATCCAGGAGATGTTCAAGCGTGTGTCTGAACAGTTTACGGCTATGTTTCGTCGTAAGGCTTTCTTGCACTGGTACACTGGCGAGGGTATGGATGAGATGGAGTTCACGGAGGCCGAGTCGAACATGAACGATCTCTCAAGGCTAGCGTGTGTGACATCCCGCCCAAGGGCCTGAAGATGAGCACTACGTTCATCGGTAACTCGACCGCTATCCAGGAGATGTTCAAGCGTGTGTCTGAGCAGTTTACGGCTATGTTCCGTCGTAAGGCTTTCTTGCACTGGTACACGGGCGAGGGTATGGACGAGATGGAGTTCACGGAGGCCGAGTCCAACATGAACGATCTTTCAAGGCCAGCGTGTGTGACATCCCGCCTCAGGGTCTCAAGATGAGCACCACGTTCATCGGCAACTCCACTGCCATCCAGGAGATGTTCAAGCGTGTTTCCGAACAGTTCACGGCGATGTTCCGTCGTAAGGCTTTCTTGCACTGGTACACGGGCGAGGGTATGGATGAGATGGAGTTCACGGAGGCTGAGTCCAACATGAACGATCTTTCAAGGCTAGCGTGTGTGACATCCCGCCCAAGGGTCTCAAGATGAGCACCACGTTCATTGGTAACTCGACTGCCATCCAGGAGATGTTCAAGCGTGTGTCCGAACAGTTCACTGCTATGTTCCGTCGTAAGGCTTTCTTGCACTGGTACACTGGTGAGGGTATGGATGAGATGGAGTTCACGGAGGCTGAGTCCAACATGAACGACCTGTCAAGGCTAGCGTGTGTGACATCCCGCCCAAGGGTCTCAAGATGAGCACCACGTTCATCGGTAACTCGACCGCTATCCAGGAGATGTTCAAGCGCGTGTCCGAACAGTTCACGGCTATGTTCCGTCGTAAGGCTTTCTTGCACTGGTACACGGGTGAGGGTATGGACGAGATGGAGTTCACGGAGGCCGAGTCCAACATGAACGATCTTTCAAGGCTAGCGTGTGTGACATCCCGCCCAAGGGCCTGAAGATGAGCACTACCTTTATCGGTAATTCGACAGCTATCCAAGAGATGTTCAAGCGTGTGTCTGAACAGTTCACGGCTATGTTTCGTCGCAAGGCTTTCTTGCACTGGTACACTGGTGAGGGTATGGATGAGATGGAGTTCACGGAGGCTGAGTCCAACATGAACGATCTTTCAAGGCCAGCGTTTGTGACATCCCGCCGAAGGGTCTCAAGATGAGCGCCACGTTCATCGGTAACTCGACTGCCATCCAGGAGATGTTCAAGCGTGTCAGCGAGCAGTTCACGGCCATGTTCCGTCGTAAGGCCTTCTTGCACTGGTACACTGGTGAGGGTATGGACGAGATGGAGTTCACTGAAGCCGAGTCGAACATGAACGATCTCTCAAGGCTAGTGTGTGTGACATCCCACCCAAGGGCCTGAAGATGAGCACCACCTTTATTGGTAATTCGACAGCCATCCAAGAGATGTTCAAGCGTGTGTCTGAACAGTTCACGGCTATGTTCCGTCGTAAGGCTTTCTTGCATTGGTACACTGGTGAGGGTATGGACGAGATGGAGTTCACGGAGGCTGAGTCTAACATGAACGATCTCTCAAGGCTAGCGTGTGTGACATCCCGCCCAAGGGGTTGAAGATGAGTACCACCTTCATCGGTAACTCGACCGCAATCCAGGAGATGTTCAAGCGCGTGTCTGAGCAATTTACGGCCATGTTTCGTCGTAAGGCTTTCTTGCACTGGTACACTGGTGAGGGCATGGATGAGATGGAGTTCACGGAGGCCGAGTCCAACATGAACGACCTCTCAAGGCTAGCGTGTGTGACATCCCGCCCAAGGGGTTGAAGATGAGTACCACCTTCATCGGTAACTCGACCGCAATCCAGGAGATGTTCAAGCGCGTGTCTGAGCAATTTACGGCCATGTTTCGTCGTAAGGCTTTCTTGCACTGGTACACTGGTGAGGGCATGGATGAGATGGAGTTCACGGAGGCCGAGTCCAACATGAACGACCTCTCAAGGCTAGCGTGTGTGACATCCCGCCCAAGGGGTTGAAGATGAGTACCACCTTCATCGGTAACTCGACCGCAATCCAGGAGATGTTCAAGCGCGTGTCTGAGCAATTTACGGCCATGTTTCGTCGTAAGGCTTTCTTGCACTGGTACACTGGTGAGGGCATGGATGAGATGGAGTTCACGGAGGCCGAGTCCAACATGAACGACCTCTCAAGGCTAGCGTGTGTGACATCCCGCCCAAGGGTCTGAAGATGAGCACTACGTTCATTGGTAACTCTACTGCTATCCAAGAGATGTTCAAGCGTGTGTCCGAACAGTTTACGGCTATGTTCCGTCGTAAGGCTTTCTTGCACTGGTACACCGGTGAGGGTATGGACGAGATGGAGTTCACTGAGGCTGAGTCCAACATGAACGATCTGTCAAGGCTAGCGTGTGTGATATTCCTCCCAAGGGTCTGAAGATGAGCACTACGTTCATCGGTAACTCGACTGCCATCCAGGAGATGTTCAAGCGTGTGTCTGAACAGTTTACGGCTATGTTCCGTCGTAAGGCTTTCTTGCACTGGTACACTGGTGAGGGTATGGATGAGATGGAGTTCACTGAGGCCGAGTCCAACATGAACGATCTGTCAAAGCTAGTGTATGTGACATTCCGCCGAAGGGTTTGAAAATGAGCACCACATTTATTGGTAATTCAACAGCTATCCAAGAGATGTTCAAGCGAGTGTCAGAACAGTTCACGGCCATGTTCCGTCGTAAGGCTTTCTTGCACTGGTACACCGGCGAGGGCATGGACGAGATGGAGTTTACTGAAGCCGAGTCGAACATGAACGATTTGTCAAGGCGAGTGTGTGTGACATCCCGCCTAAGGGTTTAAAAATGAGCACCACGTTTATTGGTAATTCGACTGCTATTCAGGAGATGTTTAAGCGAGTGTCAGAACAGTTCACGGCAATGTTCCGTCGTAAGGCTTTCTTGCATTGGTACACTGGTGAGGGAATGGACGAAATGGAGTTTACTGAGGCTGAGTCCAACATGAACGATTTATCAAGGCTAGCGTGTGTGACATCCCGCCCAAGGGTCTGAAGATGAGCACCACGTTCATCGGTAACTCGACCGC+ATCCAGGAGATGTTCAAGCGTGTGTCTGAACAGTTCACGGCTATGTTCCGTCGTAAGGCTTTCTTGCACTGGTACACTGGTGAGGGTATGGACGAGATGGAGTTCACGGAGGCCGAGTCCAACATGAACGATCT+GTGTCCGAGTACCAGCAGTACCAGGACGCCACCGGTCGCAGAGTACCAGCAGTACCAAGATGCTACAGGTCTCTGAGTATCAGCAGTACCAGGATGCCACCGGTCTCAGAGTATCAACAGTATCAAGAAGCAACCGGTTAGCGAATACCAACAATACCAAGAGGCTACTGGTGTCTGAGTACCAACAGTACCAGGATGCCACCGGTGTCTGAGTACCAGCAGTACCAGGACGCCACCGGTGTCTGAGTACCAGCAGTACCAGGACGGCACCGGTGTCTGAGTACCAGCAGTACCAGGACGCGACCGGTGTCTGAGTACCAGCAGTACCAGGACGCTACCGGTGTCTGAGTACCAGCAATATCAGGATGCGACCGGTGTCGGAGTACCAGCAGTACCAGGATGCTACGGGTGTCTGAGTACCAGCAATACCAGGATGCGACCGGTGTCCGAGTACCAGCAGTACCAGGATGCGACCGGTGTCCGAGTACCAGCAGTACCAGGATGCGACCGGTGTCCGAGTACCAGCAGTACCAGGATGCGACCGGTGTCTGAGTACCAGCAGTACCAGGACGCCACTGGTGTCTGAGTACCAGCAGTACCAGGACGCCACCGGTTTCTGAGTACCAGCAGTACCAAGACGCGACTGGTTTCTGAATACCAACAGTACCAAGATGCCACCGGTGTCTGAGTACCAGCAGTACCAGGATGCCACCG11111111111111111111209209209209209209209209209209209209209209209209209209209209102030405060708090100110120130140150160170180190200210210210210210210210210210210210210210210210210210210210210418418418418418418418418418418418418418418418418418418418418210220230240250260270280290300310320330340350360370380390400410419419419419419419419419419419419419419419419419419419419419627627627627627627627627627627627627627627627627627627627627420430440450460470480490500510520530540550560570580590600610620628628628628628628628628628628628628628628628628628628628628836836836836836836836836836836836836836836836836836836836836630640650660670680690700710720730740750760770780790800810820830837837837837837837837837837837837837837837837837837837837837104510451045104510451045104510451045104510451045104510451045104510451045104510458408508608708808909009109209309409509609709809901000101010201030104010461046104610461046104610461046104610461046104610461046104610461046104610461046125412541254125412541254125412541254125412541254125412541254125412541254125412541050106010701080109011001110112011301140115011601170118011901200121012201230124012501255125512551255125512551255125512551255125512551255125512551255125512551255125512881288128812881288128812881288128812881288128812881288128812881288128812881288126012701280
