## Supplementary figures and images for "Advancing RNAi-Based Strategies Against Downy Mildews: Insights Into dsRNA Uptake and Gene Silencing"

### Supplemental Figure 3

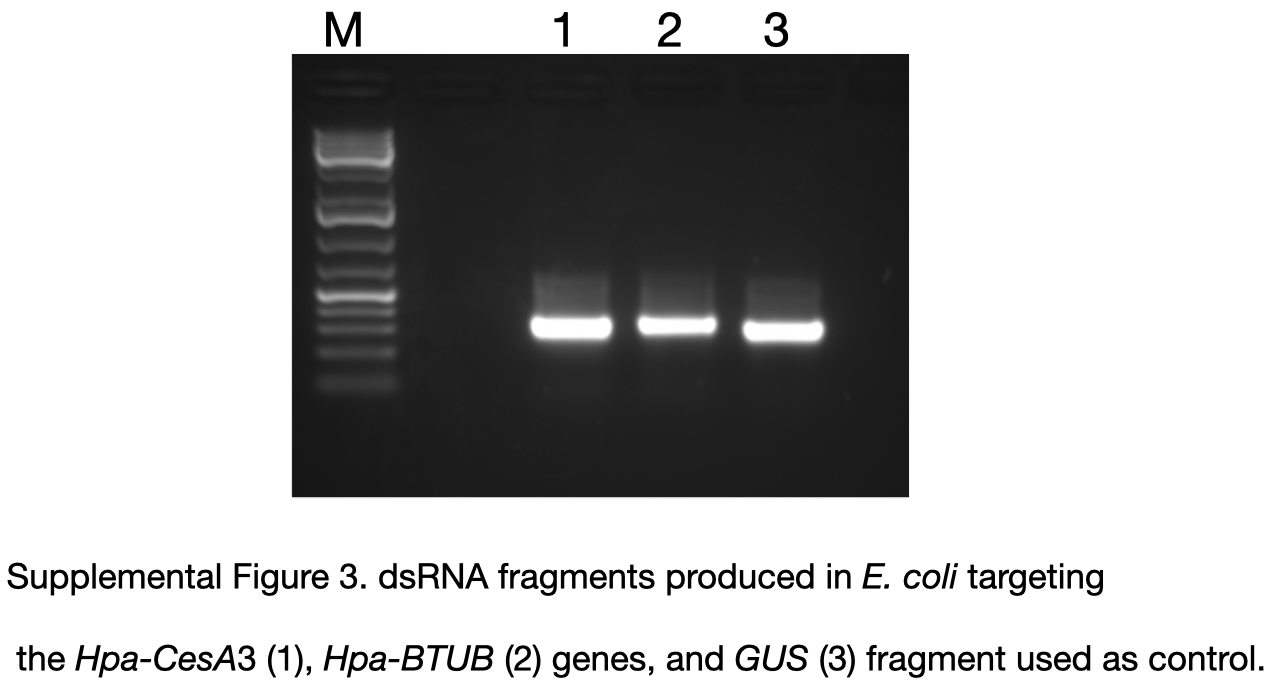

### Supplemental Figure 4

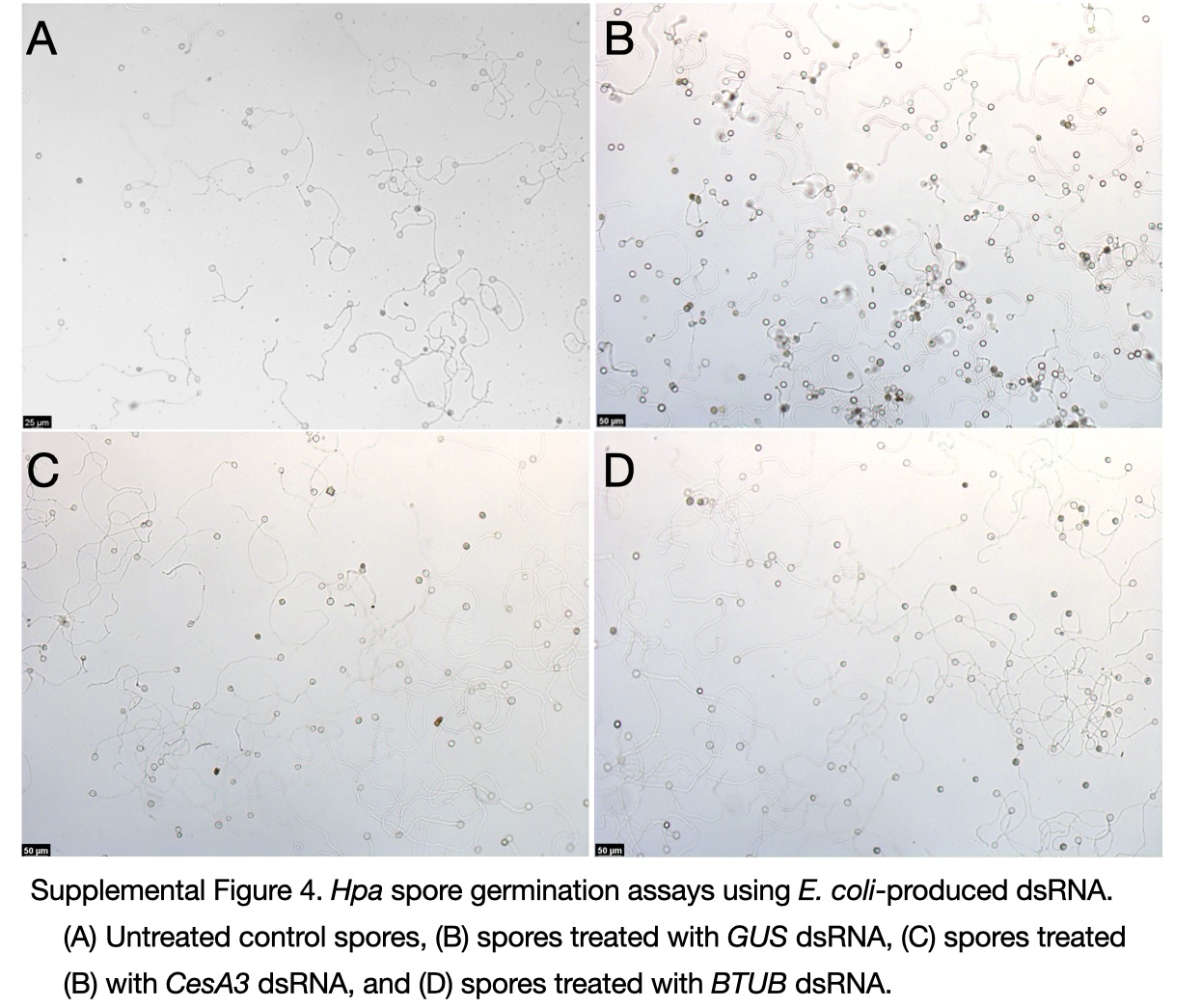
